# Mechanical regulation of cellular energy metabolism in cancer microenvironments Short title: Mechano-metabolism of metastatic cells

**DOI:** 10.1101/2024.04.30.591879

**Authors:** Joshua M. Toth, Anuja Jaganathan, Ramin Basir, Laurent Pieuchot, Yihui Shen, Cynthia A. Reinhart-King, Vivek B. Shenoy

## Abstract

Cells dynamically regulate their morphology, contractility, and metabolism in response to the mechano-chemical properties of their microenvironment. Here, we show matrix stiffness and ligand density jointly govern the bioenergetics of contractile cells through a nonequilibrium active chemo-mechanical model built around a newly introduced cellular metabolic potential. This concept links ATP hydrolysis to mechanosensitive signaling, quantifies the energetic cost of stress fiber assembly, and determines mechanically stable states. The metabolic potential enables quantitative prediction of cell contractility, morphology, and ATP consumption in different stiffness 2D and 3D environments, and we find quantitative agreement with experimental measurements in MDA-MB-231 breast cancer cells. The model further predicts activation of AMPK accompanies increased energetic demands in stiffer microenvironments which we experimentally validate and correlate with increased mitochondrial membrane potential, glucose uptake, and intracellular ATP levels. Together, these findings establish a predictive quantitative framework unifying mechanosensitive control of cell shape and contractility with the metabolic pathways sustaining cellular function across diverse mechanical environments.

**Teaser:** Matrix stiffness reshapes the cellular energy budget, driving metabolic adaptation to mechanical demand.

## INTRODUCTION

Cells sense and respond to mechanical cues through cytoskeletal reorganization and regulation of actomyosin contractility (*1*). Changes in contractility alter cell stiffness (*2*), shape (*3*), and differentiation (*4,5*) via mechanosensitive biochemical pathways. Mechanical inputs also regulate cellular metabolism through diverse mechanisms, including cytoskeletal sequestration of glycolytic enzymes (*6–9*), modulation of lipid synthesis (*10,11*), alterations in mitochondrial morphology and function (*12,13*), and YAP/TAZ signaling (*14,15*). A rapid consequence of increasing matrix stiffness is AMP-activated protein kinase (AMPK)-dependent upregulation of ATP production (*16–18*). As a master energy sensor, AMPK monitors intracellular ATP levels and coordinates glucose uptake (*19,20*), glycolysis (*21,22*), oxidative phosphorylation (*23*), protein synthesis (*24*), and autophagy (*25*), thereby maintaining energy homeostasis. Since generating and sustaining contractile forces is energetically costly, mechanotransduction and metabolic regulation are necessarily coupled. This coupling is particularly relevant in cancer progression and metastasis (*26,27*), where tumor cells reprogram metabolism in response to mechanical cues from their microenvironment (*28*). Despite extensive evidence linking matrix mechanics to metabolic adaptation, this interplay remains largely qualitative. A predictive quantitative framework that integrates mechanosensitive signaling with metabolic regulation is lacking.

Here, we develop a nonequilibrium chemo-mechanical model of contractility and adhesion centered on a newly introduced cellular “metabolic potential.” This nonequilibrium thermodynamic quantity explicitly links ATP hydrolysis by myosin-II motors to mechanosensitive stress-fiber assembly, accounting for the partitioning of chemical energy into contractile work and dissipation. Within this framework, cells adopt contractile states and morphologies that minimize their metabolic potential in a given mechanical microenvironment. The model predicts cell shape, contractility, and ATP consumption across substrates of varying stiffness and reveals that differences between 2D and 3D morphologies emerge solely from geometry-dependent stress distributions that reshape downstream chemo-mechanical signaling. We further incorporate AMPK-mediated energy replenishment and show that homeostatic regulation can sustain elevated intracellular ATP levels in highly contractile cells. These predictions are validated experimentally through measurements of cell morphology, ATP:ADP ratios, phosphorylated AMPK levels, glucose uptake, and mitochondrial membrane potential in MDA-MB-231 cancer cells cultured across stiffness gradients. Together, this work establishes a predictive quantitative framework linking cellular mechanics, mechanosignaling, and metabolic adaptation.

## RESULTS

### The Metabolic Potential Links ATP Hydrolysis to Cellular Tension

To elucidate the key principles underlying mechanosensitive metabolism, we begin with a 1D formulation, which applies to mesenchymal cells adhered between deformable microposts (Figure 1C). In contractile cells, myosin II motors bind to F-actin filaments and generate contractile forces by forming stress fibers, thereby converting the chemical energy released by ATP hydrolysis into mechanical work (*29,30*). We model individual myosin motors as force dipoles (Figure S1). The contractile stress generated by motors engaged in stress fibers is represented by ρ, while the cytoskeletal stress and strain are denoted by σ and ε, respectively. In a quiescent, non-adherent cell, we assume a baseline level of contractility, ρ = ρ₀, determined by the balance between the decrease in enthalpy upon motor binding to actin and the associated loss of entropy relative to unbound motors in the cytosol. Upon establishing adhesions, mechanosensitive signaling increases both myosin motor activity and stress fiber assembly. Specifically, cell-generated tension activates signaling pathways including Rho–ROCK (*31,32*) and Ca²⁺/calmodulin (*33*) (Figure 1A) which increase motor recruitment and contraction via ATP hydrolysis. A portion of the energy released by ATP hydrolysis is converted into (i) chemical energy associated with actin–myosin binding (U_binding_(ρ, ε)), (ii) mechanical work done by active motors (U_motor_(ρ, ε)), and (iii) elastic strain energy stored in the cytoskeleton (Ustrain^cell^ (ε)) and matrix (Ustrain^ECM^ (ε)) (Figure 1B).

**Figure 1:**
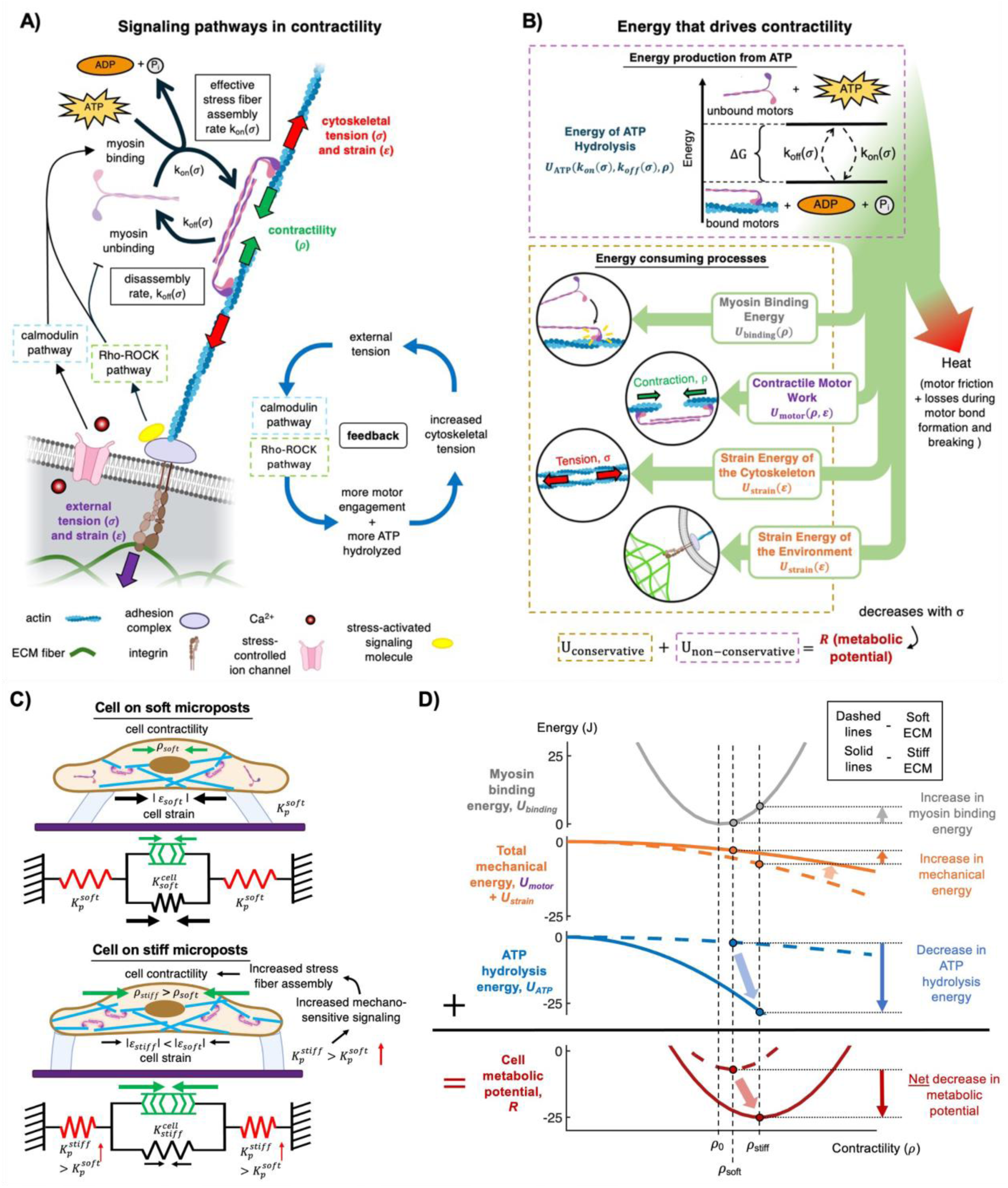
A model for the establishment of stress-regulated contractility driven by ATP hydrolysis. (**A**) Schematic illustrating the stress-dependent pathways that regulate cell contractility. Tensile forces transmitted from the micro-environment act through adhesion complexes at the cell-matrix interface and trigger: (I) Stress-assisted activation of signaling molecules (like Src Family Kinases (SFKs)), which act on Rho-GTPases and promote ROCK-mediated phosphorylation of myosin light chain (MLC) and consequently, stress fiber assembly. ROCK activation also leads to inhibition of myosin phosphatase (MYPT) which downregulates myosin unbinding and stress fiber disassembly. (II) Generation of cell membrane tension leading to the opening of stress-controlled ion-channels, an influx of Ca^2+^ into the cytoplasm, and upregulation of motor activation by myosin light chain kinase (MLCK) activation. (**B**) Schematic showing how the energy released from ATP hydrolysis by active motors in stress fibers depends on the kinetics of stress fiber assembly and comprises the dissipative part of the energy, some of which is converted into other conservative forms while the rest is dissipated as heat. We define the sum of all energies as the metabolic potential. (**C**) A cell adhered between microposts is modeled with active contractile element representing myosin motors in parallel with a linear elastic spring that describes passive cytoskeleton components. (**D**) Contributions to the metabolic potential for cells adhered to microposts as a function of contractility. Plots indicate how the various energies evolve as microposts become stiffer. The metabolic potential shows an optimum at a higher contractility on stiff (ρ_stiff_) than on soft microposts (ρ_soft_).

To describe the energy available from ATP hydrolysis, we incorporate the mechanosensitive kinetics of stress fiber assembly and disassembly through stress-dependent rate constants k_on_(σ) and k_off_(σ) (Figure 1A). These rates represent the timescale of stress fiber assembly and the lifetime of existing fibers, respectively (Figure S2). A lower disassembly rate corresponds to longer-lived stress fibers, such that k_off_ ∝ 1/τ_off_, with τ_off_ the fiber lifetime. The fraction of time that myosin motors remain engaged with actin and hydrolyze ATP is then 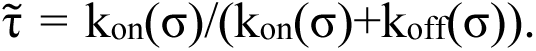 When assembly is fast and disassembly is slow, motor engagement—and thus ATP consumption—is maximized. Therefore, the time-averaged energy released by ATP hydrolysis at steady state is U_ATP_ 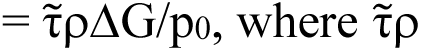 is the time-averaged density of ATP-hydrolyzing stress fibers, ΔG is the free energy released per ATP hydrolyzed, and p_0_ is the dipole strength of a single myosin dipole.

ATP hydrolysis supplies the energy needed to drive otherwise unfavorable processes, including increases in contractility and cytoskeletal and ECM deformation (Figure 1B). Because mechanosensitive signaling modulates myosin ATPase activity and stress fiber organization, it also determines how much ATP-derived energy is accessible to the cell. To capture this relationship, we introduce a metabolic potential, R, defined as the sum of conservative energies and the energy released by ATP hydrolysis (Equation S1.11). As shown in SI Section 1.1.4, minimizing R with respect to contractility and strain is equivalent to predicting the nonequilibrium steady state arising from stress fiber assembly and disassembly kinetics. Thus, the cell selects the configuration (ρ, σ, ε) that minimizes the metabolic potential—corresponding to maximal ATP hydrolysis.

To illustrate this concept, we examine a cell adhered between two microposts (Figure 1C). The cell consists of a passive elastic element of bulk modulus K and an active contractile element representing stress fibers, while each micropost is modeled as an elastic element of modulus K_p_ in series with the cell (parameter values listed in Table S1). Minimizing the metabolic potential (SI Section 1.2) predicts that steady-state contractility increases with micropost stiffness (Figure 1D). The predicted cellular traction force (SI Section 1.3) also increases with post stiffness and agrees well with experimental measurements of myoblasts contracting against deformable substrates (*34,35*). Stiffer microposts elevate mechanosensitive signaling, activate more myosin motors, and increase ATP hydrolysis, which in turn drives stronger cell contraction. The detailed dependence of mechanical energies on K, K_p_, and other parameters is provided in SI Section 1.2 and Figure S3.

Stress regulates the coupling between signaling and contractility such that k_on_(σ) increases and k_off_(σ) decreases with stress. This formulation is consistent with experimentally measured kinetics of myosin II isoforms. Myosin IIB exhibits strong catch-bond behavior, with tension decreasing its detachment rate (lower k_off_ at higher σ) and thus extending stress fiber lifetime (*36*). However, its force-dependent recruitment (increase in k_on_) is modest and cell-type specific, depending on co-assembly with myosin IIA (*37,38*). Conversely, myosin IIA is strongly recruited under tension (higher k_on_), although its catch-bond behavior is weaker (*38*). The stress-dependent forms of k_on_ and k_off_ (Equation S1.22) therefore capture both mechano-signaling–dependent motor activation and isoform-specific recruitment and dissociation kinetics.

### A 3D Chemo-mechanical formulation of the metabolic potential

While the 1D model captures stiffness sensing, real cells exhibit spatially heterogeneous and anisotropic stress distributions. We therefore generalize the framework to three dimensions, where contractility, stress, and strain become tensors, (ρ_ij_(**x**), σ_ij_(**x**), and ɛ_ij_(**x**)) that can vary with spatial coordinate, **x**, in the cell. In elongated cells, stress concentrates near elongated edges (Figure S4A), triggering mechanosensitive signaling and promoting preferential stress fiber formation along the long axes, producing a polarized contractility field (ρ₁₁(**x**) ≠ ρ₂₂(**x**) ≠ ρ₃₃(**x**)) (*39*). To capture this anisotropy, the assembly and disassembly rates, k_on,ij_ and k_off,ij_, are also treated as spatially dependent tensor quantities. Their volumetric components (k^v^ (σ_kk_(**x**)), k^v^ (σ_kk_(**x**))) represent mean-stress–dependent changes in assembly/disassembly, while their deviatoric components 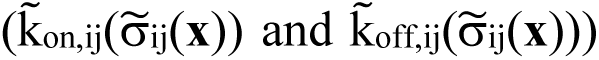 capture anisotropic stress-driven polarization of stress fibers. This tensorial description is necessary to reproduce experimentally observed spatial polarization of myosin densities in cells with anisotropic morphologies. Because anisotropic stress also upregulates contractility in elongated cells (*39*), the 3D model includes an additional anisotropy-dependent term in the assembly rate (Equation S2.14). We further introduce a cell–matrix interfacial energy, Γ(Θ), which accounts for membrane/cortical tension and adhesion energy, and depends on cell shape, described by Θ (Figure 2A).

**Figure 2:**
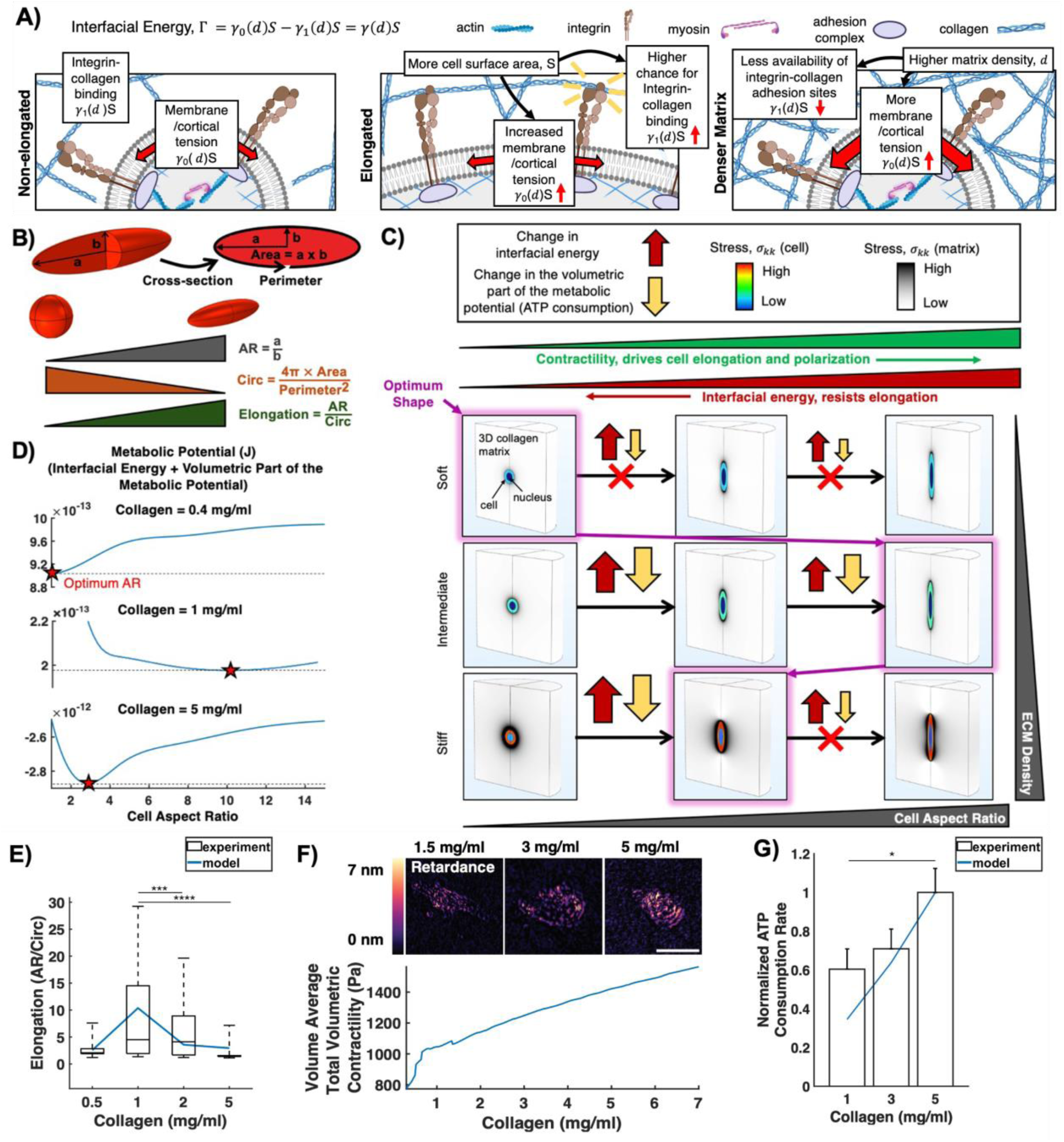
Minimizing the metabolic potential predicts a biphasic variation in cell shape with increasing collagen density. (**A**) Interfacial energy comprised of membrane/cortical tension, γ_0_, which resists increases in cell surface area, and adhesion energy of integrin-collagen binding, γ_1_, which favors cell elongation as increased surface area allows for more binding interactions. (**B**) Schematic description of cell body aspect ratio (AR), circularity (circ) and their variation with cell shape. (**C**) Optimum cell shape shows a biphasic transition from spherical in low density collagen to elongated (spindle-shaped) in medium density collagen, back to nearly spherical in high density collagen resulting from the competition between the volumetric part of the metabolic potential (red block arrows) and the interfacial energy (yellow block arrows). (**D**) Metabolic potential as a function of cell shape for low (0.4 mg/mL), medium (1 mg/mL), and high density (5 mg/mL) collagen, where the minimum in metabolic potential corresponds to the steady state optimum cell shape. Simulation predictions as compared to reported experimental measurements of MDA-MB-231 cells in 3D collagen for (**E**) cell elongation (*16*) and (**G**) normalized ATP consumption rate (*42*). The experimentally measured optical retardance signal (*45*) indicating contractile strain on f-actin is displayed in (**F**) along with model predictions of steady state contractility vs collagen density. Bar plots show mean ±SE, and box plots show median, 25^th^/75^th^ percentiles, and 5^th^/95^th^ percentiles *p<0.05, **p<0.01, ***p<0.001, ****p<0.001 with Kruskal-Wallis one-way ANOVA (**E**) or Tukey’s HSD post hoc test (**G**). Scale bar = 25 μm.

Performing a spatial integration of all contributions over their respective domains yields the generalized metabolic potential:

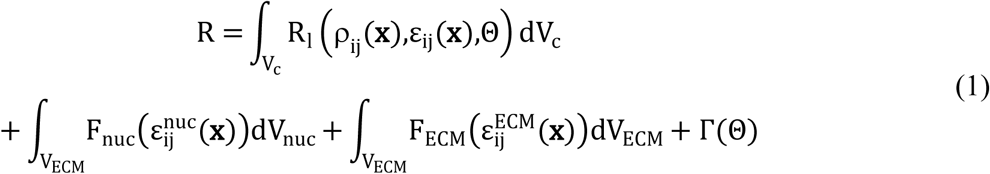

where R_l_ is the local (non-integrated) metabolic potential at every coordinate, **x** (Equation 3 in Materials and Methods), and F_ECM_ and F_nuc_ are strain energy densities of the ECM and nucleus. As in the 1D formulation, minimizing R corresponds to the steady state of the full chemo-mechanical system governed by stress fiber kinetics (SI Section 2.2.3). Thus, both the metabolic potential and the ATP hydrolysis energy follow directly from nonequilibrium thermodynamic arguments.

### Metabolic potential predicts biphasic changes in cell morphology with increasing collagen ECM density

We implemented the generalized 3D model within a finite element framework (Figure S4C) to investigate the physical mechanisms that govern the establishment of cell morphology, cytoskeletal contractility, and metabolic activity in MDA-MB-231 breast cancer cells embedded in three-dimensional collagen matrices. The cell body is represented as an ellipsoid characterized by two semiaxes of equal radius and a third distinct radius (a = c ≠ b), and the cell aspect ratio is defined as AR = a/b (Figure 2B). Thus, a spherical cell has AR = 1, whereas AR > 1 denotes an elongated cell. To consistently parameterize changes in cellular geometry, we normalized the aspect ratio by the cell circularity, circ=4π(area)/(perimeter)^2^, resulting in an elongation factor (EF) that captures shape variations across conditions (Figure 2B). Although cellular volume can vary with ECM stiffness, we assume that shape change is a volume-conserving process and set the volume of each cell in the model to 1500 µm³ (*40*). This assumption is appropriate because changes in cell volume are regulated by water transport, which requires osmotic pressures on the order of ∼1 MPa (*41*) - approximately three orders of magnitude larger than typical cytoskeletal tension (∼1–10 kPa) - making volume changes negligible relative to mechanically driven shape changes within collagen matrices.

Cell morphology, cytoskeletal contractility, and energy expenditure in 3D collagen matrices depend strongly on collagen density (*16,42*). To capture this mechano-regulation, we modeled the collagen network using a nonlinear fiber formulation (*43*) (see SI Section 2.2.5 for details; model parameters are listed in Table S2), which accounts for tension-induced fiber alignment and the resulting anisotropic matrix stiffening (Figure S4B). For each collagen density, we determined the steady-state cell shape, strain field, and contractility by solving the constitutive equations (Equations S2.20–S2.21) obtained from minimizing the metabolic potential, R (Equation 1), with respect to these variables. We repeated this analysis across a range of matrix densities and iteratively compared model predictions of cell elongation and contractility with experimental measurements for MDA-MB-231 cells to calibrate the mechanical and biochemical model parameters (Table S3).

### Actomyosin contractility and ATP hydrolysis rates are higher in elongated cells

Having established the computational framework, we examined how cell shape and collagen density jointly influence emergent contractility and energetics. As predicted in the 1D micropost setting, the steady-state volume-averaged contractility, 1/V_c_ ∫ρ_kk_(**x**)dVc, increases with collagen density because initial cell contraction generates larger volumetric stresses, 1/V_c_ ∫σ_kk_(**x**)dVc in denser matrices, which in turn lead to stronger stress-dependent actomyosin upregulation. Similarly, for any fixed matrix density, elongated cells exhibit higher steady-state contractility than spherical cells (Figure S5). This enhancement arises because elongated geometries generate larger stresses at the cell–matrix interface (Figure 2C), thereby increasing the activation of mechanosensitive signaling pathways and promoting the recruitment and activation of actomyosin machinery (Figure 1A), analogous to the effects observed in stiffer matrices.

Interestingly, in high-density collagen matrices (≥ 4 mg/ml), a different mode of stress-dependent regulation emerges. While the volume-averaged contractility increases monotonically with elongation, the volumetric stress exhibits a biphasic trend (Figure S5). This occurs because, at constant cell volume, further elongation eventually produces diminishing increases in the stress along the elongated axis (σ_33_) that are outweighed by reductions in the stresses along the transverse axes (σ_22_ and σ_11_). In soft collagen matrices, this behavior only appears at unrealistically large elongation (EF > 50), but in stiff collagen it occurs within physiologically relevant elongation levels (0 < EF < 10). However, our 3D model also includes stress-dependent actomyosin regulation arising from stress anisotropy (Equation S2.14), and anisotropy increases continuously with elongation at all collagen densities. Consequently, in high-density collagen, anisotropy-dependent signaling dominates over mean-stress–dependent signaling, causing the volume-averaged contractility to increase monotonically with elongation even when the axial stress, σ_33_ begins to decrease.

Consistent with this interpretation, our model captures the emergence of stress polarization in anisotropic cell shapes. Examination of the deviatoric stress components reveals that the component aligned with the elongated axis 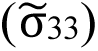 is significantly greater than those along the other axes. This polarization drives the preferential alignment of stress fibers along the first principal stress direction (Figure S6). These trends in stress and contractility determine the observed behavior of the metabolic potential and its volumetric portion, v^cell^ (all terms in Equation 1 except Γ(Θ)), reflecting the central role of stress-dependent signaling in modulating actomyosin activity.

### The optimum cell aspect ratio minimizing metabolic potential exhibits biphasic variation with collagen density

Metabolic potential dictates the optimum cell morphology with changing cell shape and collagen density through competition between its volumetric component, v^cell^, and the interfacial energy, Γ(Θ). Although all energetic terms in the volumetric metabolic potential closely track the changes in contractility across matrix densities and cell shapes, the decrease in ATP hydrolysis energy shows the largest variation with cell elongation (Figure S7). Since elongated cells develop higher contractility and thus higher ATP consumption, they exhibit lower v^cell^. However, elongation also increases cell surface area thereby increasing interfacial energy, which incorporates both membrane tension and adhesion-related contributions (Figure 2A). As detailed in SI Section 2.2.5, membrane tension is approximately two orders of magnitude larger than receptor–ligand adhesion energy and increases with both cell surface area and matrix density. Moreover, dense collagen matrices present fewer accessible adhesion sites because matrix degradation is limited (*44*). To capture these effects, the interfacial energy, Γ(Θ), is implemented as an increasing function of both cell surface area and collagen density and introduces an energetic penalty for elongation (Figure 2C). The specific functional form was determined through sensitivity analysis (Figure S8 and Table S3).

The competition between the decrease in the volumetric part of the metabolic potential, v^cell^, and the increase in the interfacial energy, Γ(Θ), generates a biphasic trend in the optimum cell aspect ratio as collagen density increases:

*Low collagen density (< 0.5 mg/ml):* In soft collagen matrices, cells cannot generate substantial stresses even when elongated (Figure 2C). The resulting stress-regulated enhancement of stress fiber assembly, and thus the ATP hydrolysis energy, is small for both spherical and elongated shapes (Figure S7). Consequently, the small decrease in the volumetric part of the metabolic potential, v^cell^, associated with elongation is insufficient to overcome the increase in the interfacial energy, Γ(Θ). The metabolic potential therefore predicts a spherical shape (Figure 2D), consistent with experimental observations (Figure 2E).

*Intermediate collagen density (0.5–3 mg/ml):* At moderate collagen densities, the matrix can support the generation of larger stresses and higher contractility when the cell elongates (Figure 2C). As a result, the reduction in the volumetric part of the metabolic potential due to elongation becomes large enough to overcome the interfacial energy penalty, and the model predicts elongated cell morphologies (Figure 2D), again matching experimental data (Figure 2E).

*High collagen density (> 3 mg/ml):* In dense matrices, the decrease in the volumetric part of the metabolic potential, v^cell^, that accompanies elongation is insufficient to compensate for the increase in interfacial energy beyond an aspect ratio of approximately 3, and the cell therefore adopts a moderately elongated shape (Figure 2D). This prediction is in close agreement with experimental measurements (Figure 2E). Notably, the difference in v^cell^ between spherical and elongated cells is smaller in dense matrices, since high matrix stiffness already enables cells of nearly any shape to generate substantial contractility (Figure 2C). Thus, the metabolic potential does not provide a strong driving force for elongation. For a clear visualization of how the variation in the volumetric part of the metabolic potential and interfacial energy produces the trends described here, we plot and compare their exact values in Figure S9.

In summary, only at intermediate collagen densities is the stress-and contractility-dependent reduction in the volumetric portion of the metabolic potential large enough to offset the interfacial energy penalty associated with elongation. This competition produces an energy landscape that leads to a biphasic dependence of the optimum cell aspect ratio on collagen density (Figure 2D). Indeed, we find that without interfacial energy, the shape that minimizes the volumetric portion of the metabolic potential at any density is always very highly elongated (Figure S10).

### Metabolic and mechanical consequences of morphology selection

Despite the nonmonotonic dependence of cell shape on collagen density, both the steady-state volume-averaged contractility and the ECM strain energy of the optimal cell shape increase monotonically with collagen density because the amount of stress generated by the cell (and thus the amount of stress-related signaling the cell experiences) increases with ECM stiffness (Figures 2F, S11A). These predictions agree with quantitative polarization microscopy (QPOL) measurements, which report optical retardance due to F-actin strain and increase with collagen density (Figure 2F). The model also predicts a monotonic increase in ATP consumption rate at steady state (Equation S2.27), consistent with PercevalHR measurements of ATP depletion following inhibition of oxidative phosphorylation and glycolysis (*40*) (Figure 2G).

Finally, to further validate our framework, we simulated myosin inhibition by reducing the stress fiber assembly rate k_on,ij_. This perturbation significantly reduced steady-state cytoskeletal stress, mirroring the effects of Y-27632 treatment in 3D collagen (Figure S12A,B) (*45*). Myosin inhibition also prevented cells from adopting elongated shapes in intermediate-density collagen (1.5 mg/ml), reflecting the fact that reduced contractility weakens the reduction in the volumetric portion of the metabolic potential, v^cell^, normally associated with elongation.

### Metabolic potential predicts cell flattening and elongation on stiffer 2D substrates

We next asked whether the metabolic potential framework could accurately predict single-cell morphologies in environments beyond 3D collagen, such as on two-dimensional polyacrylamide (PA) substrates commonly used to study mechanosensitivity as a function of matrix stiffness (*46*). To address this, we predicted MDA-MB-231 cell shape and contractility on 2D substrates of varying stiffness (Young’s modulus) using *the same biophysical parameters* determined for cells in 3D collagen (Table S3), without introducing any additional fitting parameters. The only modification is the interfacial energy term, Γ(Θ), which must account for the fact that cells on 2D substrates interact with the ECM only along their basal surface rather than being fully encapsulated (SI Section 2.2.5). In addition, because cells form larger and stronger adhesions on stiffer substrates (*47*), the receptor–ligand adhesion energy is scaled with substrate stiffness, rather than implemented as a function decreasing with matrix density as in 3D collagen.

Cells on 2D substrates are modeled as half-ellipsoids with semiaxes a, b, and c (Figure 3A), where in general a ≠ b ≠ c. When all three semiaxes are equal, the cell adopts a hemispherical shape. To parameterize cell morphology, we define two aspect ratios: a/b, which captures elongation in the plane of the cell–matrix interface, and c/b, which characterizes the cell height relative to its in-plane width (Figure 3A). Cell volume is held constant, so the spreading area (A=πab) increases as the vertical axis, c, shortens. Thus, a decrease in c/b indicates cell flattening, while an increase in a/b indicates cell elongation. Although a continuous family of shapes was explored, we found that three representative morphologies capture the dominant energetic effects: (i) hemispherical (a/b=1, c/b=1), (ii) moderately elongated spindle-like (a/b>1,c/b<1), and (iii) highly elongated and flattened spindle-like (a/b≫1,c/b≪1) (Figure 3B). The PA substrate is treated as a linear elastic material (*46*) with stiffness varying between 1 and 200 kPa.

**Figure 3:**
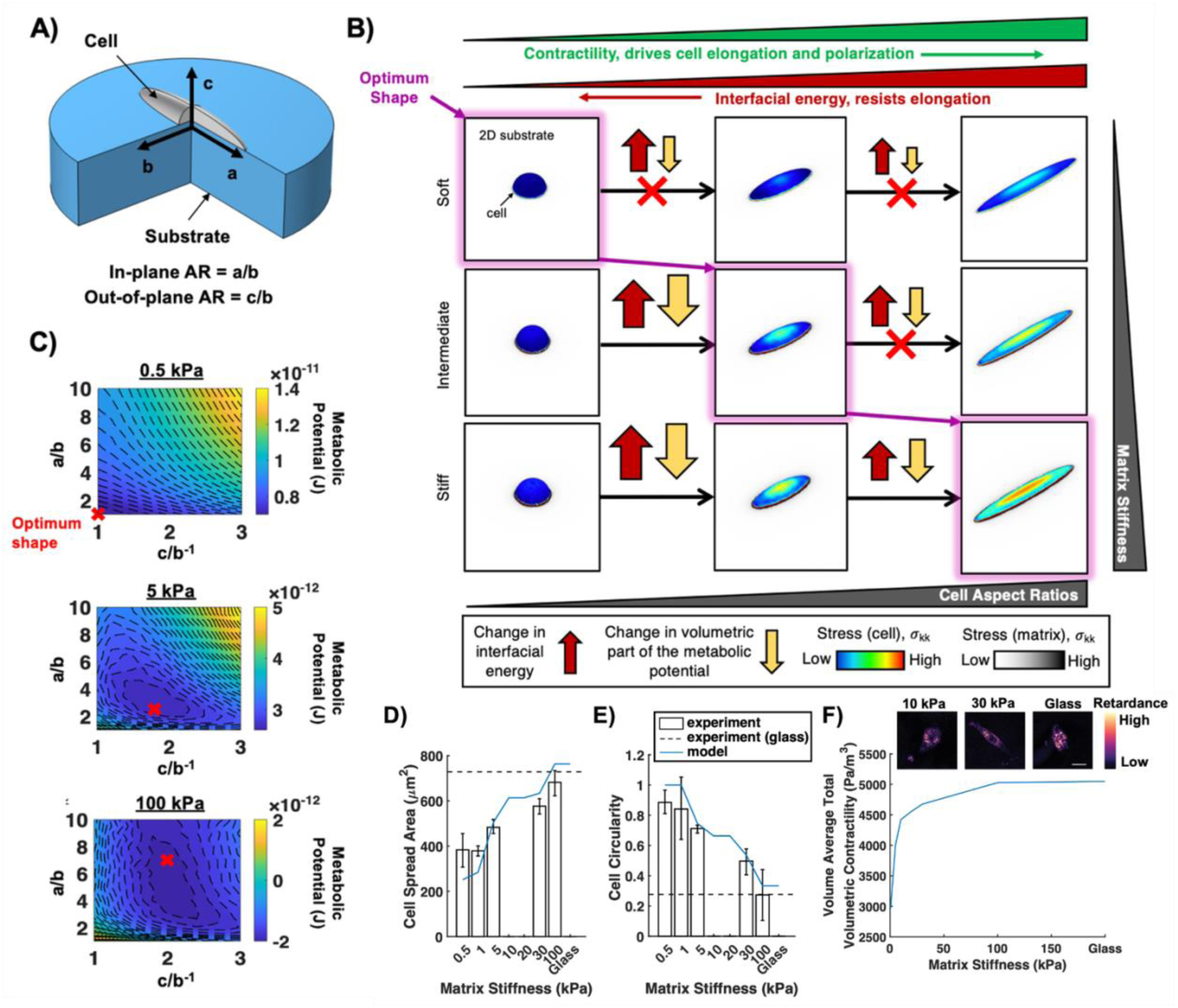
Minimizing the metabolic potential predicts increasing cell elongation and flattening with 2D matrix stiffness. (**A**) Schematic depiction of the geometry of a cell on a 2D substrate. (**B**) Optimum cell shape shows a transition from hemispherical on soft gels to increasingly elongated and flattened on intermediately and very stiff PA gels as determined by the interaction between the volumetric part of the metabolic potential (red block arrows) and the interfacial energy (yellow block arrows). (**C**) Contour plots of the metabolic potential as a function of the cell aspect ratios a/b and c/b with the optimum cell shape indicated. Comparison of (**D**) cell spread area, (**E**) cell circularity, and (**F**) cell contractility at steady state predicted by the model with the corresponding data from experiments with MDA-MB-231 cells seeded on 2D substrates reported in (*45,48*). Bar plots show mean ±SD. Scale bar = 25 μm.

### Anisotropic cell shapes on 2D substrates develop elevated and polarized cytoskeletal stresses

Cells on 2D substrates exhibit mechanosensitive trends similar to those observed in 3D collagen. As substrate stiffness increases, they develop larger volume-averaged stresses, which lead to higher steady-state contractility and increased ATP hydrolysis by active motors (Figure S13). Likewise, as the cell elongates within the 2D plane, both the stress field and the contractility field become increasingly polarized due to preferential stress fiber assembly along the direction of elongation (Figure S14A,B). The optimum cell shape on 2D substrates is still determined by the competition between the volumetric portion of the metabolic potential, v^cell^, and the interfacial energy, Γ(Θ). Substrate stiffness modulates this competition by regulating the magnitude of stress the cell can generate at steady state, and thus the degree of mechanosensitive upregulation of stress fiber assembly.

However, the predicted dependence of optimal cell morphology on matrix stiffness differs substantially from the biphasic trend observed in 3D collagen. In 2D, cells *monotonically elongate and flatten* with increasing substrate stiffness (Figure 3B). This monotonic trend arises because the cell can only adhere to and transmit traction forces through its basal surface, meaning the region of highest stress is always located near the cell–matrix interface regardless of shape (Figure 3B). Flattening the cell increases the fraction of the cytoskeletal network that lies close to this high-stress region and prevents stress relaxation in regions farther from the interface. Consequently, flattened morphologies support larger stresses throughout the cell body. These larger stresses increase ATP hydrolysis and reduce the volumetric portion of the metabolic potential, v^cell^, providing a strong energetic incentive for both flattening and elongation as stiffness increases.

Interfacial energy opposes both flattening and elongation because each increases cell surface area. However, as substrate stiffness rises, the cell’s ability to develop larger contractility more effectively reduces v^cell^, enabling it to overcome the increasing interfacial energy penalty. As a result, the competition between v^cell^ and Γ(Θ) produces a minimum in the metabolic potential that shifts systematically toward shapes with higher a/b (more elongated) and lower c/b (more flattened) as stiffness increases (Figure 3C). The resulting morphology ranges from a nearly hemispherical cell on soft substrates (< 1 kPa) to a highly elongated, spindle-like cell on stiff substrates (> 100 kPa). This predicted monotonic elongation and flattening is in excellent agreement with experimental measurements of MDA-MB-231 spread area and cell circularity on PA substrates (*48*) (Figure 3D,E). Consistent with the predictions from 3D collagen, the volume-averaged stress, contractility, and ATP hydrolysis all monotonically increase with substrate stiffness (Figure S14C–E). These predictions also match experimental assessments of cell contractility via optical retardance measurements (*45*) (Figure 3F).

In summary, we find cells do not exhibit a biphasic morphology trend in 2D because, unlike in 3D matrices, a flattened configuration always provides the most favorable means of maintaining high stresses on stiff substrates. Therefore, there is no energetic advantage to returning to a less flattened shape once stiffness becomes large. Interestingly, this driving force for a cell to adopt the shape which maximizes contractility and minimizes metabolic potential can accurately predict the array of morphologies observed when cells are placed in environments with different stiffness and different geometry.

### A Predictive Framework for Mechanosensitive ATP Replenishment in Microenvironments with Variable Matrix Stiffness

Thus far, we have shown that despite differences in optimum morphologies, cells in stiffer environments in both 2D and 3D develop higher contractility and correspondingly exhibit greater ATP consumption. We now extend our model to develop a theoretical framework describing how cells regulate their energetic budget in response to the increased bioenergetic demands imposed by stiff microenvironments. Our objective is to quantitatively predict key metabolic indicators—including ATP levels, ATP:ADP ratios, glucose uptake, and oxidative phosphorylation (OXPHOS) rates—as functions of ECM stiffness. We first provide a quantitative description of ATP consumption in adherent, contractile cells and then outline the proposed mechanosensitive ATP replenishment pathways, along with the model parameters used to quantify them. Full derivations and parameter determination are provided in SI Section 3.

Living cells maintain a high, nonequilibrium ATP:ADP ratio, typically in the range of 1–10 (*49*). To model ATP consumption, we consider the ATP hydrolysis reaction (reaction 1 in Table S4), in which the rate constants k_1_ ^+^ and k_1_ ^-^ govern ATP consumption and ATP synthesis, respectively. Although cells may adjust many energy-intensive processes in different microenvironments, here we focus on ATP consumption associated with changes in motor activity driven by stress fiber assembly. We therefore take the ATP consumption rate constant k_1_ ^+^ to depend on the rates of stress fiber assembly and disassembly, which themselves depend on cytoskeletal stress. Accordingly, k_1_ ^+^ increases with the total volumetric stress (see SI Section 3 for the functional form).

Once the finite element formulation is solved for a given cell shape on a 2D substrate or in 3D collagen, we compute the total stress by integrating σ_kk_(**x**) over the cell volume. We also incorporate the conversion of ADP to AMP (reaction 3 in Table S4) by introducing the ratio γ′ of the reverse and forward rate constants (k_2_^+^ and k_2_ ^-^). Increased cytoskeletal tension and ATP consumption on stiff substrates lead to a greater reduction in ATP:ADP ratio and a corresponding rise in the AMP:ATP ratio compared with soft substrates.

### Mechanosensitive, calcium-dependent activation of AMPK drives ATP replenishment

A central regulator of cellular metabolism is AMP-activated protein kinase (AMPK), which integrates signals related to cellular energy levels and controls major catabolic and anabolic pathways, including OXPHOS and glycolysis (*17*). AMPK is a heterotrimeric complex composed of the catalytic AMPKα subunit and regulatory AMPKβ and AMPKγ subunits. The activation of AMPK is initiated by the allosteric binding of AMP to AMPKγ. As cellular energy levels fall, AMP concentration increases, promoting higher levels of AMP-bound AMPK. This effect is amplified in stiffer microenvironments, where greater contractility and ATP consumption elevate AMP levels (Figure 4A).

**Figure 4:**
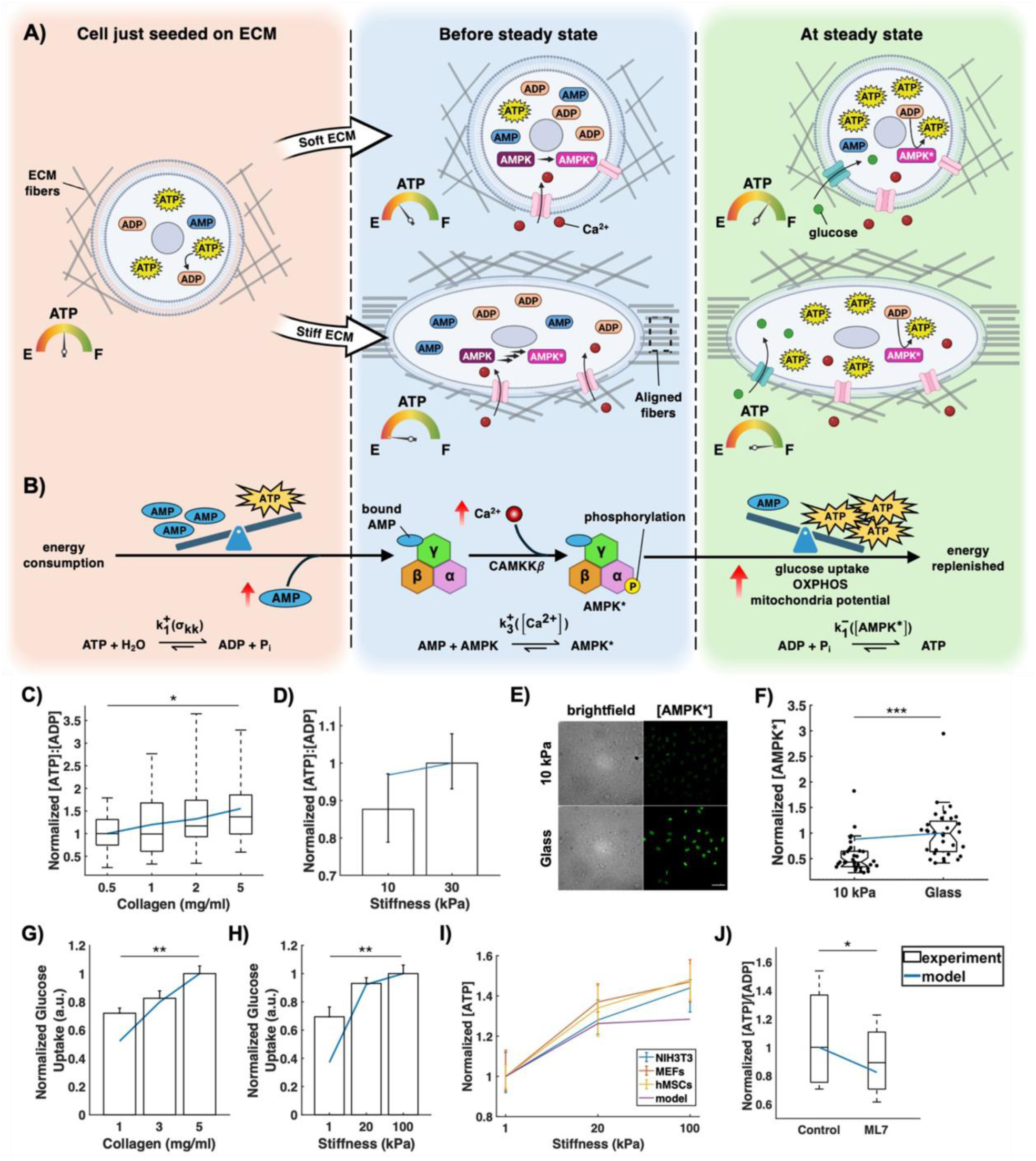
Model predicts ATP replenishment based on the mechano-sensitive calcium-mediated activation of AMPK. (**A**) Schematic representation of the changing ATP, ADP and AMP levels immediately after a cell is seeded, before reaching steady state, and at steady state on soft and stiff ECMs. (**B)** Schematic depiction of AMPK activation in response to AMP and calcium levels. ECM stiffness regulates AMPK activation by affecting both cell energy consumption and calcium signaling as the cell evolves towards steady state. (**C**) Predicted ATP:ADP ratio vs. collagen density and (**D**) PA gel stiffness compared to experimental measurements with PercevalHR from (*16*). (**E**) Confocal max intensity projections of phosphorylated AMPK in MDA-MB-231 cells cultured on 10kPa 2D PA gel and glass. (**F**) Quantification of (**E**) compared with model predictions of [AMPK*]. Model predictions of [AMPK*] are compared with change in rate of glucose uptake for MDA MB-231 cells in (**G**) 3D collagen and (**H**) hMSCs on 2D gels from references (*18,42*). (**I**) Predicted ATP levels compared with experimental measures in different cell types from (*18*). (**J**) Predicted ATP:ADP levels in cells with normal and lowered stress fiber assembly rates compared against experimental measurements for MDA MB-231 cells in 3D collagen treated with contractility inhibitor ML7 from (*42*). Bar and line plots show mean ±SE, and box plots show median, 25^th^/75^th^ percentiles, and 5^th^/95^th^ percentiles *p<0.05, **p<0.01, ***p<0.001, ****p<0.001 with Kruskal-Wallis one-way ANOVA (**C,D,F**), Tukey’s HSD post-hoc test (**G,H**) or Wilcoxon rank test (**J**). Scale bar = 50 μm.

AMP binding induces a conformational change that enhances phosphorylation of AMPKα by upstream kinases. Two kinases—LKB1 (liver kinase B1) and CaMKKβ (Ca²⁺/calmodulin-dependent protein kinase kinase β)—promote phosphorylation of AMPKα at Thr172. Although LKB1 expression does not appear to depend on ECM stiffness, cytoplasmic calcium levels increase on stiffer substrates (*50–52*), especially in mesenchymal cells where elevated RhoA signaling and increased membrane tension both enhance calcium influx (*52,53*). Higher intracellular calcium leads to higher CaMKKβ activation and thus higher levels of phosphorylated (activated) AMPK, denoted here as AMPK*(Figure 4B).

AMPK activation is represented by reaction 4 in Table S4, where k_3_ ^+^ denotes the activation rate constant and k_3_ ^-^ the deactivation rate constant. To capture the mechanosensitive nature of AMPK activation, we take k_3_ ^+^ to be a function of cytoskeletal stress (and thus intracellular calcium), i.e., k_3_ ^+^ = k_3_ ^+^(σ_kk_). The ATP synthesis rate constant k_1_ ^-^ is an increasing function of AMPK*, reflecting AMPK-mediated upregulation of ATP production (reaction 2 in Table S4).

Mathematical expressions for steady-state concentrations of ATP, ADP, AMP, and AMPK* are obtained by setting time derivatives to zero and enforcing conservation of total nucleotide concentration and total AMPK. This yields a system of five nonlinear equations that we solve simultaneously.

### Cells sustain higher ATP:ADP ratios and AMPK activation on stiffer matrices

Simplifying our system of equations results in a predicted relationship between the ATP:ADP ratio and total volumetric stress:

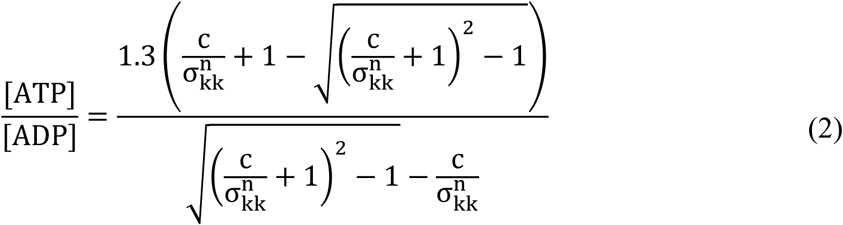

where c and n represent combinations of other model parameters that arise while solving (see SI Section 3.1). Equation 2 predicts that the steady state ATP:ADP ratio will increase with increasing stress (or increasing ECM stiffness) if both c and n are positive. Parameter c is always positive since all its factors such as k_2_^+^, k_2_^-^, and the total AMPK concentration, [AMPKtot], are all positive by definition. However, n describes the difference between the change in the rate parameters k_1_^+^ and k_3_^+^ in response to a change in stress and will only be positive if k_3_^+^ (the rate of calcium-dependent AMPK activation) shows a larger increase in response to stress than k_1_^+^ (the rate of ATP consumption by motors). In other words, Equation 2 reaffirms that the ATP:ADP ratio will increase with ECM stiffness only when cells experience sufficient AMPK activation and subsequent ATP production to overcome the rate at which ATP is consumed by motors. We fitted Equation 2 to experimentally measured ATP:ADP ratios for collagen densities ranging from 0.5 to 5 mg/mL (Figure S15A) to determine the value of parameters c and n (Table S5). Our framework captures the trend that ATP:ADP ratios increase with collagen density (Figure 4C) and accurately predicts that cells maintain steady-state nucleotide concentrations with [ATP]>[ADP]>[AMP] (Figure S15B,C).

To validate the generality of the model, we predicted ATP:ADP ratios for cells cultured on 2D PA substrates using the same parameters. The predicted ATP:ADP ratio increases monotonically with substrate stiffness, in agreement with experimental observations (Figure 4D). Although cell morphology responds differently to stiffness in 3D collagen and 2D PA gels, the ATP:ADP ratio increases in both environments because the optimum configuration adopted by cells on stiffer substrates supports higher stress and contractility (Figures 2F and 3F), leading to greater ATP consumption and a corresponding increase in ATP replenishment.

Our framework also predicts that AMPK activation levels increase as a function of matrix stiffness (Equation S3.18). Measurements of phosphorylated AMPK (Thr172) from immunostained cells cultured on soft versus stiff substrates show significant increases in AMPK activation on stiffer substrates, in good agreement with our predictions (Figure 4F). These findings are consistent with reports showing that AMPK activation increases with stiffness on both short and long timescales (*18*). We find that activated AMPK localizes strongly to the nucleus (Figure 4E), consistent with other reports of nuclear accumulation of AMPK under conditions of metabolic stress (*54,55*).

Model predictions of increased AMPK activation in stiff microenvironments correlate with experimentally observed increases in glucose uptake (Figure 4G,H), elevated mitochondrial membrane potential (a proxy for OXPHOS rate) (Figure S16), and increased ATP levels across multiple cell types (Figure 4I) (*16,18,42*). Increased ATP levels with stiffness are a general feature of mesenchymal cells, and the model captures this trend well. To assess whether changes in ATP:ADP ratio indeed reflect changes in motor activity, we simulated myosin inhibition by reducing the stress fiber assembly rate k_on,ij_ over 200-fold. Motor inhibition reduced steady-state cytoskeletal stress (Figure S12B), AMPK activation, and ATP replenishment, producing significantly lower ATP:ADP ratios consistent with experimental results using myosin light chain kinase inhibitors such as ML7 (*42*) (Figure 4J). These findings support the idea that cytoskeletal contractility is a major determinant of energy budget regulation across stiffness environments.

In summary, we find a mathematical analysis accounting for the mechanosensitive chemical reactions governing the generation and interconversion of ATP, ADP, and AMP is sufficient to explain experimental measurements of intracellular ATP:ADP ratio and AMPK activation in contractile cells in different stiffness environments. On stiffer substrates, motor ATP consumption and calcium mechanosignaling work in parallel to upregulate AMPK activation and ensure cellular energy production meets energy demand.

### Restricted spreading reduces cytoskeletal stress and metabolic activity

Previous work has shown that constraining cells to micropatterned substrates that limit their ability to spread or elongate alters their metabolic state, including the ATP:ADP ratio (*16*). Using our model, we examined how cytoskeletal stress and ATP consumption are modulated in cells confined to restricted geometries. For cells adhered to PA substrates of 10 or 30 kPa, we considered two cases: 1) fixed spread area with varying elongation, and 2) fixed elongation with varying spread area. For each shape, we solved for the steady-state stress field and computed the volume-averaged cytoskeletal stress (1/V_c_ ∫σ_kk_dV_c_) followed by the predicted ATP:ADP ratio using Equation 2 and the biophysical parameters in Table S5.

Our model predicts that ATP:ADP ratios increase with cell elongation when spread area is held constant, consistent with experimental measurements (Figure 5A). We obtain similar results when modeling cells on patterns of increasing spread area at a constant elongation (Figure 5C). Confining patterns that restrict elongation or spreading force the cell into more hemispherical shapes, which exhibit greater stress relaxation and lower volume-averaged cytoskeletal stress, ultimately reducing contractility. Cells with lower contractility consume less ATP and consequently exhibit reduced AMPK activation and ATP replenishment at steady state. For a given elongation, ATP:ADP ratios are higher on stiffer substrates, reflecting the enhanced stress-supported contractility (Figure 5A). Model predictions of AMPK activation also agree closely with observed mitochondrial membrane potential changes across micropattern geometries that limit elongation (Figure 5B). When spread area increases at fixed elongation, ATP:ADP ratios again increase, matching experimental findings, as does mitochondria membrane potential (Figure S17).

**Fig. 5.**
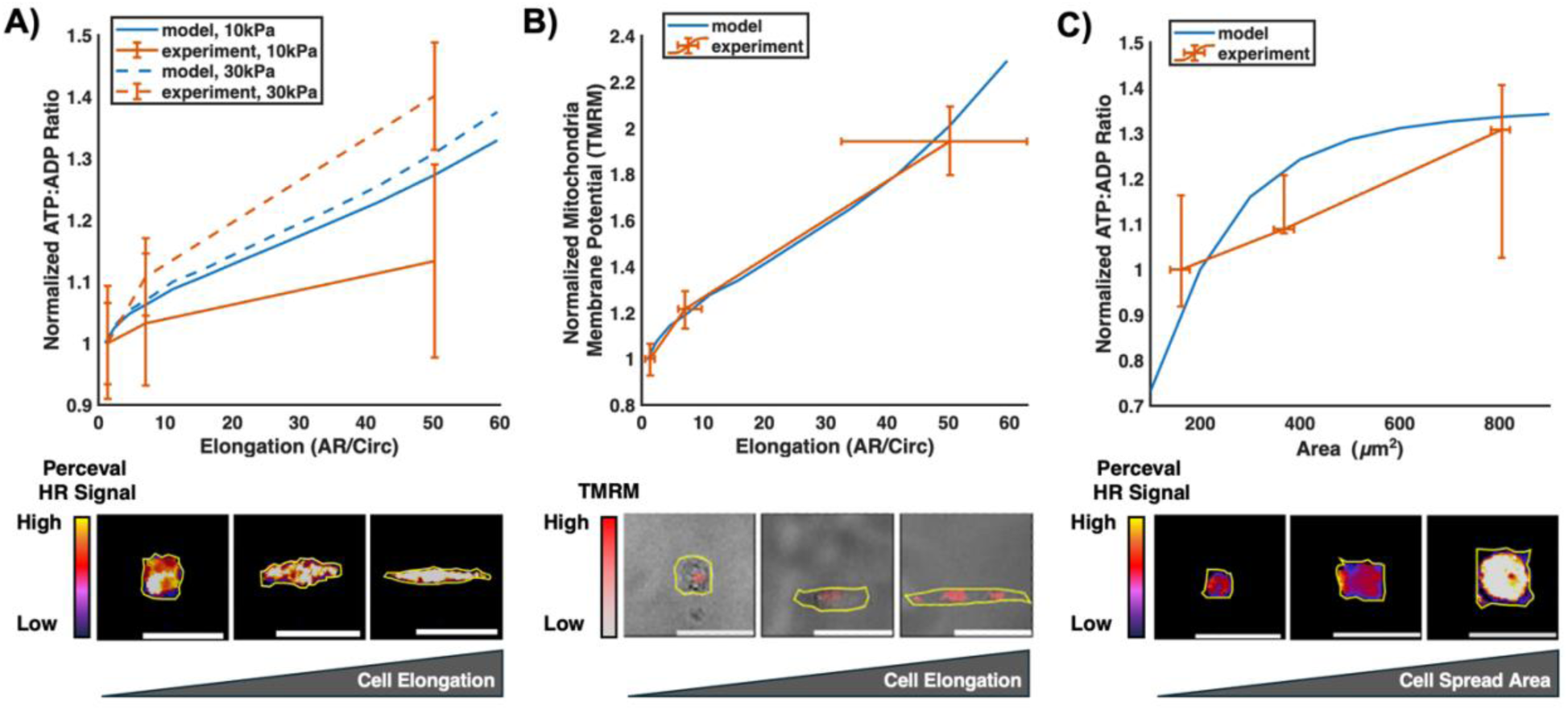
Model predicts decreases in intracellular energy levels and mitochondria membrane potential observed in cells on confining micropatterns. (**A**) Normalized [ATP]:[ADP] ratio as measured with PercevalHR and (**B**) mitochondria membrane potential as measured with tetramethylrhodamine methyl ester (TMRM) in MDA-MB-231 cells cultured on collagen micropatterns of fixed area fabricated on PA gels of 10 kPa or 30 kPa along with model predictions. (**C**) Normalized ATP:ADP ratio measured in cells on micropatterns of fixed elongation (elongation ∼1) compared to model predictions. Plots show mean ±SD. Scale bar = 50 μm. Experimental data and images obtained from (*16*).

Together, these results further validate the proposed mechanosensitive ATP replenishment model in contexts where cell spreading is externally controlled and where long-term cytoskeletal remodeling dominates cellular energy expenditure.

## DISCUSSION

In this study, we developed a nonequilibrium chemo-mechanical model for adherent cells that quantitatively connects mechanosensitive signaling to the regulation of cellular energy budgets and morphological adaptations in response to microenvironment stiffness (Figure 6). Central to this framework is the concept of a metabolic potential, defined as the sum of energies associated with establishing contractility, including myosin binding energy, motor work, cytoskeletal and matrix strain energies, energy released from ATP hydrolysis, and the cell–matrix interfacial energy. We demonstrate that minimizing this metabolic potential yields contractility and strain fields identical to those derived from the kinetics of mechanosensitive stress fiber assembly, consistent with nonequilibrium thermodynamics.

**Figure 6.**
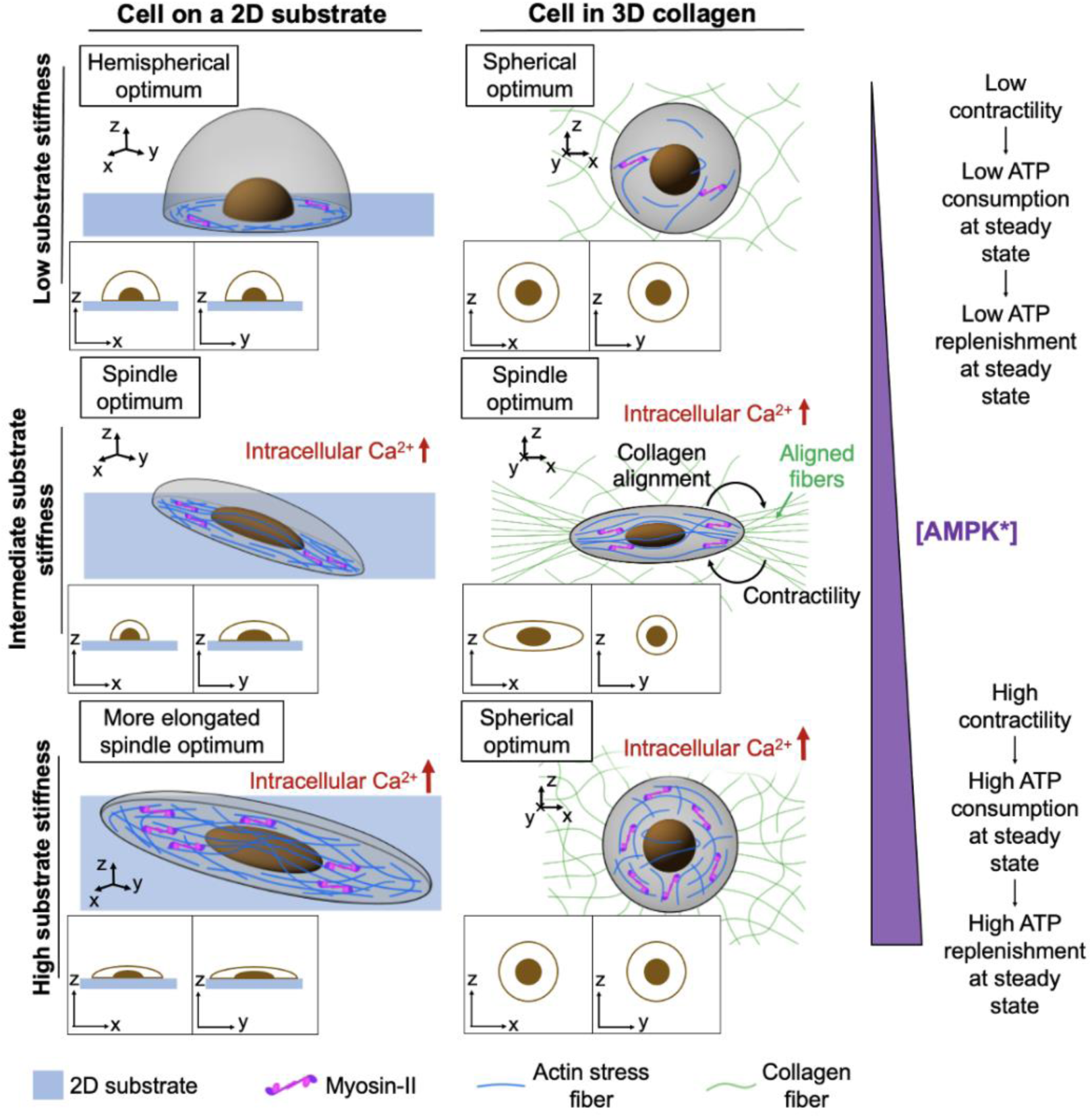
A model to predict how cell shape, contractility, and metabolism depend on mechanosensitive stress fiber assembly in adherent cells. Cell shape and ECM stiffness determine the distribution and magnitude of stresses generated in an adherent cell which, through mechanosensitive pathways, regulate the establishment of stress fibers. The decrease in energy achieved by coupling the ATP hydrolysis with stress fiber assembly drives the cell towards a more contractile state and to adopt morphologies that maximize ATP consumption and minimize interfacial energy. Simultaneously, the same mechanosensitive pathways upregulate cytosolic calcium levels which leads to the activation of AMPK through a calmodulin mediated pathway, replenishing cellular ATP stores.

Using this formulation, we predicted the steady-state cell shape and contractility of MDA-MB-231 cells in two distinct contexts: embedded within 3D collagen matrices and adhered to 2D polyacrylamide substrates. Across both environments, elongated cells exhibit increased contractility and stress-fiber polarization, which lower the metabolic potential, while elongation simultaneously increases interfacial energy, creating a competition that determines the cell’s optimal morphology. The model predicts a biphasic dependence of cell elongation on collagen density in 3D matrices and a monotonic elongation and flattening with increasing stiffness on 2D substrates, matching experimental observations. These results highlight the central role of geometry and stress distribution in determining cell morphology under different mechanical constraints.

Our predictions agree with experimentally observed cell shapes in different collagen densities and PA stiffnesses (*16*), and they are consistent with behaviors reported for fibroblasts and other mesenchymal cells (*56*). Previous reports of increased collagen fiber alignment with cell contraction in dense matrices support our finding that cells generate larger contractile forces in stiffer microenvironments (*57,58*). Very high-density matrices can suppress traction when they restrict matrix degradation, as observed for hMSCs in covalently crosslinked hydrogels (*59*), but this effect is unlikely to influence MDA-MB-231 cells at the range of densities examined here. Our model also predicts matrix strain energies that are consistent with reported experimental measurements (*60,61*). Differences in strain energy between nonlinear collagen gels and linear PA gels are captured by the model’s incorporation of tension-driven fiber alignment, which enables long-range force transmission (*43*).

We further introduced a theoretical description of ATP replenishment mediated by mechanosensitive activation of AMPK. Because ATP hydrolysis fuels contractility, cells increase metabolic output to meet the demands imposed by stiffer microenvironments. The model predicts increases in ATP:ADP ratio, glucose uptake, mitochondrial potential, and AMPK activation with ECM stiffness, consistent with experimental data in both 2D and 3D environments (*16,42*). The reduction in ATP:ADP ratio observed upon myosin inhibition provides additional support for the direct coupling between cytoskeletal tension, AMPK activation, and cellular energy homeostasis. Predictions for cells on micropatterns further demonstrate how reduced spreading or elongation lowers cytoskeletal stress, metabolic activity, and AMPK signaling.

An important aspect of our framework is that all ATP consumed at steady state is dissipated as heat, since there is no net addition of stress fibers and no net change in conservative energy. The heat dissipation rate therefore represents the maximum mechanical power output of the cell. Using experimentally constrained parameters (Table S6), we estimate that a cell in 0.5 mg/mL collagen generates approximately 0.81 fW of power. Since 1 fW corresponds to the hydrolysis of roughly 20,000 ATP molecules per second (*62*), this power output implies that such a cell consumes approximately 16,000 ATP molecules per second to maintain its steady-state configuration. On soft 1 kPa PA gels, the predicted power output is around 0.38 fW. In dense collagen (5 mg/mL), this value increases to approximately 6.5 fW, corresponding to more than 100,000 ATP molecules per second. These predictions are comparable to reported single-cell power measurements and illustrate the substantial energetic investment required to sustain high contractility in stiff environments (*35*).

The results of this study suggest several natural extensions of the framework. The metabolic potential formulation can be generalized to describe dynamic behaviors such as cell spreading, migration, durotaxis, and mechanically induced transitions in contractile state. Such processes involve time-dependent changes in stress fiber organization, signaling, and AMPK activation, and incorporating these dynamics could provide deeper insight into transient and adaptive mechanosensitive responses. The model could also be adapted to account for cell-type-specific mechanosensitive programs, enabling comparisons across different cancer cell lines, stem cells, or immune cells that experience distinct mechanical and metabolic constraints. Differences in motor kinetics, adhesion dynamics, or metabolic capacity could be incorporated to explore how distinct populations balance mechanical demand with energetic supply.

More broadly, the framework provides a foundation for linking single-cell energetics to tissue-level mechanical environments. Mechanical heterogeneity within tumors, fibrotic tissues, or regenerating matrices may impose spatial patterns of energetic demand that govern collective behavior, metabolic adaptation, and invasiveness. Understanding these relationships may help identify mechanically regulated vulnerabilities in cancer cells or inform the design of engineered microenvironments that direct stem cell function or modulate immune activation.

Altogether, by establishing a quantitative link between mechanical signaling, cytoskeletal organization, and energetic regulation, this model offers a predictive and extensible approach for understanding how cells sense, respond to, and adapt to the mechanical properties of their surroundings. These insights provide a basis for future explorations of mechanosensitive metabolic regulation across diverse physiological and pathological settings

## METHODS

### A 3D chemo-mechanical formulation of the metabolic potential

Here, we briefly outline (refer to SI Section 2 for details) a general 3D model for the metabolic potential of a contractile cell to describe the effect of stress-activated signaling pathways on the metabolic budget. The key mechanical components of this model are a) stress fibers composed of active contractile elements in parallel with passive components representing the cell cytoskeleton, b) a poroelastic nucleus, and c) the extracellular matrix or substrate.

#### A) Stress fibers

In the 3D model, contractility, stress, and strain are treated as tensor fields, reflecting the cell’s ability to assume nonspherical shapes with spatially nonuniform anisotropic stresses and locally varying motor orientations (Figure S4A). In this case, the contractility field (motor density associated with stress fibers) is anisotropic, i.e., ρ_11_≠_22_≠_33_, as is the stress field, and both can vary spatially across the cell body necessitating a dependence on spatial coordinates represented by vector 𝐱. The volumetric part of the contractility tensor ρ_kk_(**x**) = (ρ_11_(**x**)+ρ_22_(**x**)+ρ_33_(**x**))/3 represents contraction averaged over all directions and hence is a measure of the density of motors per unit volume in the cell. The deviatoric part, 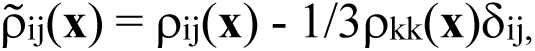 is a measure of the deviation of each component of ρ_ij_ from the mean and represents the degree of motor polarization and, by extension, stress fiber polarization.

The cell experiences activation of mechanosensitive pathways dependent on the strength of the stress generated, σ_ij_. These pathways regulate myosin motor activity and associated ATP consumption by affecting the stress-dependent rates of stress fiber assembly (k_on,ij_(σ_ij_)) and disassembly (k_off,ij_(σ_ij_)). k_on,ij_ and k_off,ij_ are tensorial quantities which allow for anisotropy in stress fiber assembly and disassembly governed by anisotropy in the stress field. The exact functional dependence of the components of k_on,ij_ and k_off,ij_ on the components of σ_ij_ is informed by the biophysical mechanisms that underlie motor activation and stress fiber formation. For example, we assume that stress fiber kinetics are linearly proportional to the average stress at every point such that the volumetric parts of the on and off rates (k^v^_on_ and k^v^_off_) are proportional to the volumetric stress (i.e., k^v^ (σ_kk_(**x**)) = (k_on,11_(σ_kk_(**x**)) + k_on,22_(σ_kk_(**x**)) + k_on,33_(σ_kk_(**x**)))/3) while the deviatoric parts 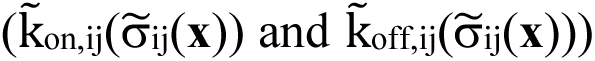 allow for upregulated stress fiber formation rates in the direction of larger stresses. This captures the fact that mechanosignaling and an increase in the total number of active motors may arise from an increase in stress along any direction, but the cell will experience a greater rate of stress fiber formation in the direction of higher stress. However, we have previously shown that contractility regulated solely by volumetric (average) stress is insufficient to explain experimentally observed increases in cell contractility in elongated cell morphologies characterized by anisotropic stress distributions (*39*). This discrepancy arises from the formation of anisotropic adhesions aligned with polarized stress fibers, which establish bidirectional mechano-chemical feedback between cell shape and signaling (*63*). To model this process, we introduce a non-linear term to the rate of stress fiber assembly, k^a^, that accounts for increases in cell contractility but only when the stress field is anisotropic. This is readily accomplished by expressing tensor quantities in terms of their principal components to identify stress anisotropy independent of any coordinate system. The principal components of stress, σ_i_(**x**) where σ_1_(**x**) ≥ σ_2_(**x**) ≥ 0, reveal stress anisotropy exists whenever the first principal component, σ_1_(**x**), is larger than the second principal component, σ_2_(**x**). We introduce this mathematically by assigning k^a^ a functional dependence on the term (σ_1_(**x**)/σ_2_(**x**)-1) such that k^a^ is nonzero and leads to increased stress fiber assembly only when σ_1_(**x**) > σ_2_(**x**), or when the stress fields are sufficiently anisotropic (see SI Section 2 for a full description).

Using these quantities, we define the conservative energy densities at every point in the cell, including the myosin binding energy (Equation S2.8), motor work (Equation S2.9), strain energy of the passive components (Equation S2.10), and the energy from ATP hydrolysis (Equation S2.14) (see SI Section 2.2 for a full tensorial description of these energies). Physically, this formulation captures the fact that ATP hydrolysis is energetically downhill and can therefore drive otherwise unfavorable processes, such as myosin binding and the straining of the cytoskeleton and extracellular matrix. However, because the release of this energy is directly coupled to the mechanosensitive activity of myosin motors within stress fibers, the kinetics of stress-dependent and stress-anisotropy-dependent signaling pathways regulate the rate at which ATP energy can be utilized. We incorporate this effect by defining the stress fiber assembly rates, k_on,i_ and k^a^, as increasing functions of the local principal stress, σ_i_, and the disassembly rate, k_off,i_, as a decreasing function of stress. Consequently, when the cell experiences high levels of cytoskeletal tension - such as on stiff extracellular matrices - the energy available from ATP consumption increases due to both an elevated rate of myosin motor engagement and an increased motor binding lifetime.

#### B) Adhesions at the cell-matrix interface

Transmembrane proteins at the cell-ECM interface, such as integrins, form a bond with ligands on the ECM and play an important role in transmitting forces to the cell cytoskeleton. To capture the cellular morphology in 2D and 3D microenvironments, we also introduce an interfacial energy, Γ, which is a function of the cell shape. There are two contributions to the interfacial energy: the cell surface tension, γ_0_ which resists any increase in the cell area or elongation, and an adhesive contribution from membrane protein‒ligand binding interactions, γ_1_, which favors an increase in the cell surface area along the cell-matrix interface (Figure 2A). Cell membranes consist of lipid bilayers tethered to an underlying rigid actin–myosin cortex. Hence, any change in the cell membrane area is accompanied by unfavorable stretching of membrane-engaged proteins that resist expansion in the cell area as well as by sustained contraction of the underlying actomyosin cortex. We consider these effects to comprise the energetic cost of stretching the cell membrane. The adhesive element represents the reduction in energy achieved due to bond formation between receptors such as integrin at the cell-matrix interface and ligands present in the matrix. The relative strengths of these two quantities determine whether the total interfacial energy Γ = (γ_0_ - γ_1_)S will increase or decrease with cell surface area, S. Additionally, we account for changes in the adhesive energy with changing ECM ligand density and changing membrane protein expression with ECM stiffness, as explained further in SI Section 2.2.5.

#### C) The nucleus

We assume that the viscous relaxation of the nucleus is negligible (see SI Section 2.1.2) and model the nucleus as a linear elastic ellipsoidal inclusion encapsulated within the cell with elastic properties K_nuc_ and μ_nuc_ denoting its bulk and shear modulus, respectively. We assign the nucleus a volume 1/5 of the cell volume and vary its aspect ratios in the same manner as cell aspect ratios when considering different shape cells.

#### D) Mechanical response of the microenvironment

We consider two types of cell microenvironments in this work: 1) cells encapsulated in 3D collagen and 2) cells seeded on 2D PA gels. Following our previous work, the mechanical behavior of collagen is described using a fibrous nonlinear constitutive model that captures the strain-stiffening effect of fiber alignment under tension (*43*) (see SI Section 2.2.6). The bulk and shear moduli of collagen as a function of its density were determined from experimental measurements of the equilibrium modulus of collagen gels (*16*). 2D PA gel substrates are modeled as linear elastic, as AFM measurements have shown that gels with stiffness values of physio-pathological relevance exhibit ideal linear elastic behavior even at the nanometer scale (*46*).

### Generalized 3D expression for the metabolic potential of an adherent cell

We define the local metabolic potential, R_l_, at every point **x** in the cell in terms of principal components by summing all energetic contributions:

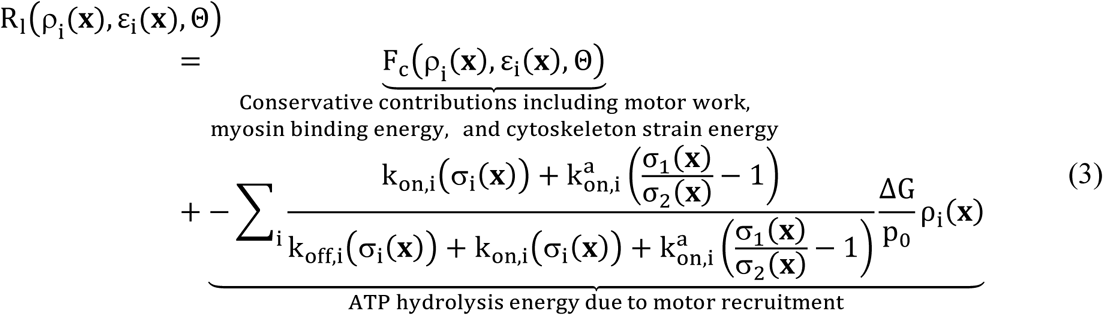

The total metabolic potential, R, is then calculated by integrating R_l_ over the cell volume and adding the strain energy from the nucleus and ECM as well as the interfacial energy, Γ(Θ) (Equation 1). The metabolic potential represents the potential of a cell to enhance its contractility and alter its morphology/shape in response to the mechanical properties of its microenvironment by considering mechanosensitive cytoskeletal kinetics. Minimization of the metabolic potential with respect to the contractility, strain, and shape yields the rate of ATP consumption in the cell at steady state (Equation S2.25).

To determine the optimum cell shape and contractility as a function of matrix stiffness, we employed a finite element procedure, as described in SI Section 2.3 (Figure S4C). Briefly, we construct cell geometries with varying cell body aspect ratios and discretize the geometry within a finite element framework. At every spatial coordinate **x**, we determine the steady-state cellular contractility and strain by minimizing the local metabolic potential R_l_ with respect to ρ_ij_(**x**) and 𝜖_ij_(**x**) through calculation of functional derivatives of R with respect to these variables. This yields a set of equations that we solve using simulations for a given cell body aspect ratio (Equations S2.20-S2.21). Then, knowing the spatial variation of the contractility and strain, we determine the cell metabolic potential, R (Equation 1), corresponding to a particular cell body aspect ratio by integrating the local metabolic potential, R_l_, over the cell and matrix volume and adding the interfacial energy corresponding to that shape. This process is repeated for different cell body aspect ratios, and the optimum cell shape is identified as the cell body aspect ratio that has minimum metabolic potential. We repeated this procedure for different matrix stiffness values to generate the observed variations in the optimum cell shape and contractility with stiffness.

### Experimental measurement of AMPK activation

To compare with model results, we performed immunostaining and confocal imaging of cells cultured on 2D polyacrylamide substrates to determine activation of AMPK at the single-cell level in response to changing microenvironment stiffness. MDA-MB-231 cells (ATCC HTB-26) were cultured in growth media consisting of DMEM supplemented with glutamax, sodium pyruvate, Anti-Anti (Gibco), and 10% FBS (Cytiva) before trypsinization and resuspension for growth on substrates. Polyacrylamide substrates were fabricated on pre-treated glass coverslips by mixing different volumes of acrylamide, bis-acrylamide, tetramethylethylenediamine (TEMED), and ammonium persulfate (APS) as previously described (*56,64*). Final percentages by volume were 7.7%:0.26%:0.27%:0.9% (10kPa) in dH_2_O. Coverslips were pretreated to promote polyacrylamide binding by incubation in 0.1M NaOH for 3 minutes, 0.5% (3-Aminopropyl)trimethoxysilane (3-APTMS, Sigma) for 30 minutes, and 0.5% glutaraldehyde (Sigma) overnight. All incubations were performed at room temperature and preceded by multiple washes with dH_2_O. Coverslips were dried, and the acrylamide mixture was polymerized for 30 minutes at room temperature while sandwiched between the coverslips and siliconized glass, the latter of which could be easily removed with gentle force afterward. Resulting PA gels were surface coated with collagen to promote cell adherence. This was accomplished by incubation in 0.05% sulfo-sanpah (Sigma) in PBS during treatment with UV light for 10 minutes. Gels were then washed multiple times with PBS and incubated in 0.1 mg/ml rat tail type I collagen (Corning) for 2 hours at room temperature. Gels were again washed multiple times with PBS and sterilized under UV for 1 hour. After at least 24 hours, cells were seeded in growth media at 25% confluency. For the glass stiffness condition, coverslips simply underwent the final incubation step in 0.1 mg/ml collagen before sterilization and cell seeding.

Samples were cultured for 24 hours prior to 15 minute fixation with 4% paraformaldehyde, permeabilization with 0.5% triton X-100 (Sigma) for 20 minutes, and blocking with 3% bovine serum albumin (Sigma) for 1 hour with repeated washing between each step. Samples were then stained overnight at 4°C with phosphorylated AMPK primary antibody (Cell Signaling Technology #2531) and 1 hour at room temperature with secondary antibodies (Abcam 150077) prior to confocal imaging. For each cell, a bright field image as well as a confocal z-stack at 235 x 235 μm (512 x 512 pixels) and 0.5 μm vertical spacing was obtained using a Zeiss LSM980 point scanning confocal microscope equipped with a 40x water immersion lens and a Prime BSI Express sCMOS camera. Samples were imaged by inverting the gels into a No. 0 35 mm glass-bottom dish (MatTek) containing Fluoromount-G mounting media (Invitrogen) to allow imaging of the cells through the glass bottom of the dish. Relative levels of phosphorylated AMPK were quantified for each cell by finding the total fluorescence signal using ImageJ software (*65*). Confocal images were z-projected for max intensity and the extent of each cell body was manually determined by using reference bright field images. Total expression for each cell (Figure 4F) was then calculated by normalizing the sum of all pixel values to cell area and the mean expression of all cells on glass. Full details of how model predictions were compared with previously published experimental data are available in SI Section 3.2.

## Statistical Analysis

All box and whisker plots show median (center line) ± interquartile ranges (box edges) and 5^th^ and 95^th^ percentiles (whiskers). Bar plots show mean and standard error or mean and standard deviation where indicated. Statistical analysis was performed with MATLAB version 2023b (*66*) and significance was found using Kruskal-Wallis ANOVA (*67*) with Dunn’s post hoc analysis (*68*). Experiments were reproduced independently at least three times. Nonlinear least squares regression was performed using MATLAB’s lsqcurvefit algorithm.

## Acknowledgments

We express our gratitude to X. Chen for assistance with development of early versions of our models, E. Sorokina for assistance in maintaining cell cultures, and C. Madl for allowing generous use of his imaging system.

## Author contributions

Conceptualization: VBS, CRK, JT, AJ Methodology: VBS, JT, AJ Investigation: JT, AJ

Visualization: JT, AJ Supervision: VBS Writing—original draft: JT, AJ

Writing—review & editing: VBS, CRK, LP, YS

## Data and materials availability

All data are available in the main text or the supplementary materials. We provide open access to all files used to generate and run our models as well as custom written code used to analyze model results in a public GitHub repository: https://github.com/ShenoyLab/Metabolism. Within, readers will find instructions to download, install, and reproduce model data. Unprocessed images from novel experiments are also available within this repository.

