## Supplementary Information for "Mechanical regulation of cellular energy metabolism in cancer microenvironments Short title: Mechano-metabolism of metastatic cells"

Joshua Toth *et al.*

**This PDF file includes:**

Supplementary Text
Figs. S1 to S17
Tables S1 to S6
References (69 to 81)

### Supplementary Text

1. Active, non-equilibrium model for the metabolic potential of adherent contractile cells

*1.1. Development of the 1D metabolic potential for cells adhered between microposts:*

We present here a model for the metabolic potential of cells as a function of cytoskeletal strain $\epsilon$ and contractility $\rho$ and show that minimizing it (with respect to $\rho$ and $\epsilon$) can yield the steady state conditions predicted by considering the dynamic equilibrium of stress fibers in adherent cells. We first formulate a metabolic potential for a cell held between two microposts (*34*) (Figure 1C) with the goal of predicting how the cytoskeletal contractility is modulated by the stiffness of the extracellular micro-environment and how the change in contractility is directly correlated to the amount of ATP consumed. This analysis lays the groundwork for the more general case of cells in 3D collagen networks and 2D gels. The chief components of the model considered here are: a) stress fibers, which consist of myosin motors engaged to actin filaments, modelled as an active contractile element, and b) the cytoskeleton represented as an isotropic linear elastic material. This can be represented as a linear elastic spring (represents the passive components) in parallel with the contractile element (see Figure 1C). The mechanical properties of the spring represent the mechanical properties of the cytoskeleton. The molecular myosin motors are modelled as force dipoles (see Figure S1) and the contractile force exerted by the motors on the actin filaments generates an internal stress in the cell, which we represent by the scalar variable $\rho$.

*1.1.1. Conservative energy contributions to the metabolic potential:*

The conservative energy has the following contributions, namely a) the binding energy of myosin motors, b) motor work and c) the strain energy of the cytoskeleton and the ECM.

*Myosin Binding Energy:* In the quiescent state, the isotropic contractility in the cell is denoted by $\rho_0$. This is the baseline level of contractility determined by the balance between reduction in enthalpic energy due to formation of bonds between actin and the decreased entropy of motors in the cytosol. When forces are generated and transmitted from the cell to the matrix, stress fiber assembly is upregulated through stress-dependent signaling pathways (see Figure 1A) and hence the contractility increases. Any deviation from $\rho_0$ is, however, accompanied by an increase in the "chemical" energy of the actomyosin system, which represents the increase in energy when free-floating myosin motors from the cytosol bind to actin filaments. This energy is called the myosin binding energy ($U_{binding}$) and is given as:

$$U_{binding} = \frac{\beta}{2}(\rho - \rho_0)^2 V_c \tag{S1.1}$$

In the above equation, $\beta$ is a measure of the chemical stiffness of the cell, and $V_c$ is the cell volume. Physically, $\beta$ represents the penalty for any change in contractility from its quiescent state value $\rho_0$. The larger $\beta$ is, the more energetically expensive it is for the cell to increase/decrease its contractility from the quiescent state level.

*Mechanical Energy:* The mechanical energy consists of three terms which describe the work done on the cell by the myosin motors, the strain energy of the cell, and work done by the mechanical stresses at the cell-matrix interface. As the motors contract, they apply internal forces to the cytoskeleton and consequently, do mechanical work, which we call motor work, $U_{motor}$:

$$U_{motor} = \rho \epsilon V_c \tag{S1.2}$$

While we provide a precise mathematical derivation for $U_{motor}$ in a generalized 3D framework in SI Section 2.1, note that this 1D non-tensor form arises from the fact that the contractility, $\rho$, is a measure of motor density (or density of force dipoles) in the cell that when multiplied by cytoskeletal strain, $\epsilon$, and cell volume, $V_c$, results in the total work done by motors. The contracting motors deform the cytoskeleton, and the strain energy associated with the deformation is given as:

$$U_{strain}^{cell} = \frac{1}{2} K \epsilon^2 V_c \tag{S1.3}$$

where $K$ is the Young's modulus of the cell and $\epsilon$ is the strain induced in the cell cytoskeleton.

To illustrate the ability of the model to capture the mechanical features of the cell's environment, we choose to examine the simple case of a cell adhered to microposts (Figure 1C). We can account for the strain energy of the two microposts by including the term:

$$U_{strain}^{posts} = K_p \epsilon_p^2 l_p^2 \tag{S1.4}$$

where $K_p$ is the stiffness of the microposts, $\epsilon_p$ is the strain in the posts, and $l_p$ is the distance between the microposts. The sum of the binding energy and the mechanical energy represents the conservative part of the energy which we denote as $U_c(\rho, \epsilon)$.

*1.1.2.* *Kinetics of stress fiber assembly and steady state energy dissipation:*

We now derive the condition for the dynamic steady state of the cell constrained by the posts from a first-principles consideration of cytoskeletal kinetics and contraction based on the second law of thermodynamics. If the rate of assembly and disassembly per unit volume of stress fibers is denoted $J_{on} > 0$ and $J_{off} > 0$, respectively, we can write the rate of change of $\rho$ as $\frac{\partial \rho}{\partial t} = J_{on} - J_{off}$. For each motor that binds to actin filaments forming stress fibers, ATP is hydrolyzed to release energy $(\Delta G / p_0)$ which we denote as $\Delta \tilde{G}$. Here, $\Delta G$ is the energy released from the hydrolysis of one molecule of ATP, and $p_0$ is the dipole strength of one myosin motor. The energy available through ATP hydrolysis per unit time due to stress fiber formation throughout the entire cell is then $J_{on} \Delta \tilde{G} V_c$. Part of this energy is dissipated as heat, while the rest is converted into the conservative forms of energy (Figure 1B). The heat dissipated per unit time $\dot{D}$, can be found as the energy available through ATP hydrolysis minus the energy stored in conservative forms as a result of stress fiber assembly and disassembly:

$$\dot{D} = \underbrace{J_{on}\Delta\tilde{G}V_c}_{\substack{energy\\ from\\ ATP\ per\ unit\ time}} - \underbrace{J_{on}\left(\frac{\partial U_c}{\partial \rho}\right)}_{\substack{rate\ of\ conservative\\ energy\ change\\ upon\ stress\ fiber\ formation}} + \underbrace{J_{off}\left(\frac{\partial U_c}{\partial \rho}\right)}_{\substack{rate\ of\ conservative\\ energy\ change\\ upon\ stress\ fiber\ disassembly}}$$

$$= -J_{on}\left(\frac{\partial U_c}{\partial \rho} - \Delta\tilde{G}V_c\right) + J_{off}\left(\frac{\partial U_c}{\partial \rho}\right) \geq 0.$$

(S1.5)

We invoke the second law of thermodynamics which states that the dissipation rate should always be greater than zero, and hence choose the following forms for $J_{on}$ and $J_{off}$:

$$J_{on} = -k_{on}(\sigma)\left(\frac{\partial U_c}{\partial \rho} - \Delta\tilde{G}V_c\right) \text{ and } J_{off} = k_{off}(\sigma)\left(\frac{\partial U_c}{\partial \rho}\right).$$

(S1.6)

where $k_{on}$ and $k_{off}$ are positive rate parameters that govern the rates of stress fiber assembly and disassembly, respectively (Figure S2). A functional dependence of these parameters on the cytoskeletal stress $\sigma$ is introduced to account for the mechano-sensitive upregulation in stress fiber formation and their subsequent stabilization under stress (Eq S1.22). At steady state, a dynamic equilibrium reflecting the elevated rates of mechano-signaling is established and the rate of assembly is balanced by the rate of disassembly, i.e., $J_{on} = J_{off}$, and the contractility remains constant over time. Enforcing this condition leads to:

$$J_{on} - J_{off} = -\frac{\partial U_c}{\partial \rho} + \frac{k_{on}(\sigma)}{k_{on}(\sigma) + k_{off}(\sigma)}\Delta\tilde{G}V_c = 0,$$

(S1.7)

The above equation relates contractility to stress by accounting for the kinetics of these processes. As we show below, choosing $k_{on}(\sigma)$ to be an increasing function of stress and $k_{off}(\sigma)$ to be a decreasing function of stress implies that rates of stress fiber formation and their lifetime increases with stress (e.g. on stiff substrates). It should be noted that the steady state heat dissipation rate obtained from Eq S1.5 is $J_{on}\Delta\tilde{G}V_c$, implying that all the energy released by myosin ATP consumption is dissipated as heat at steady state. We find a Hill-like relation between rate of contraction and stress (Eq S1.20) that emerges as a natural consequence of the Second Law of Thermodynamics. Note that the energy dissipated due to motor friction during contraction can also be calculated, but since motors cease to move in the "stall" state (or steady state) when the cytoskeletal stress $\sigma$ equals the stall stress, $\sigma_m$ (Eq S1.21c), the velocity of contraction, $\dot{\epsilon}$, is zero, and there is no contribution to overall dissipation in this case. Next, we describe the energy available from ATP hydrolysis during cytoskeletal contraction considering the mechano-sensitive character of the rate parameters $k_{on}$ and $k_{off}$.

*1.1.3. Energy of ATP hydrolysis:*

The hydrolysis of ATP is downhill energetically, and studies have estimated the value of energy released at ~30 kJ/mol (*62*). Active non-muscle myosin-II motors utilize ATP hydrolysis to power a conformational change in the motor that facilitates the formation of crosslinks between actin and myosin and subsequent force generation (*30*). On stiff substrates, there is an upregulation in the number of stress fibers formed per unit volume due to stress-induced mechanosensitive signaling through the Rho-ROCK and calcium pathways (Figure 1A). These pathways lead to both an increase in the rate of stress fiber formation and a decrease in the rate of stress fiber disassembly such that fibers remain intact for longer times on stiffer substrates. Consequently, a higher steady state contractility and higher ATP consumption are achieved on stiff substrates. To capture the stress regulated kinetics of stress fiber formation, we introduce the rate parameters $k_{on}$ and $k_{off}$, that govern the rate of stress fiber assembly and disassembly, respectively. As shown in Figure S2, if we denote $\tau_{off}$ as the average stress fiber lifetime in the assembled state and $\tau_{on}$ as the average lifetime in the disassembled state then:

$$\tau_{off}(\sigma) = \frac{1}{k_{off}(\sigma)}$$

and (S1.8a)

$$\tau_{on}(\sigma) = \frac{1}{k_{on}(\sigma)}$$

(S1.8b)

Here, $\tau_{off}(\sigma)$ is assumed to be an increasing function of stress, which represents the physical phenomenon that stress fibers are reinforced by cytoskeletal stress on stiff substrates. Also, $\tau_{on}(\sigma)$ is taken to be a decreasing function of stress which represents the upregulation in the rate of stress fiber formation on stiff substrates (Figure S2). Thus, the time for which a stress fiber remains intact and exerts contractile force over a cycle of assembly and disassembly events, denoted by $\tilde{\tau}$, can be written as:

$$\tilde{\tau}(\sigma) = \frac{\tau_{off}(\sigma)}{\tau_{off}(\sigma) + \tau_{on}(\sigma)} = \frac{k_{on}(\sigma)}{k_{on}(\sigma) + k_{off}(\sigma)}$$

(S1.9)

As noted in the Main Text, the specific properties of different myosin II isoforms will determine how the rate of stress fiber formation and the cell's contractility changes in response to stiffness but assuming $k_{on}(\sigma)$ to be an increasing function of stress and $k_{off}(\sigma)$ to be a decreasing function of stress accounts for the known properties of the most common isoforms, IIA and IIB. If we denote $(\Delta G/p_0)$ as the energy released by ATP hydrolysis due to a single motor, and recalling that $\rho$ represents the number density of motors in stress fibers per unit volume, we can write the following simple expression for the energy available through ATP hydrolysis:

$$U_{ATP} = -\tilde{\tau}(\sigma)\left(\frac{\Delta G}{\rho_0}\right)\rho V_c = -\frac{k_{on}(\sigma)}{k_{on}(\sigma) + k_{off}(\sigma)}\left(\frac{\Delta G}{p_0}\right)\rho V_c$$

(S1.10)

The above equation provides the important link between the energy released through ATP hydrolysis and the dynamic process of stress fiber assembly/disassembly.

*1.1.4. Metabolic Potential:*

We now define the expression for the metabolic potential of the cell, $R$, as the sum of all contributions, both conservative and the free energy of ATP hydrolysis:

$$R(\rho,\epsilon) = U_{binding} + U_{motor} + U_{strain}^{cell} + U_{strain}^{posts} + U_{ATP} = U_c + U_{ATP}$$
$$= \left[\frac{\beta}{2}\left(\rho - \rho_0\right)^2 + \rho\epsilon + \frac{1}{2}K\epsilon^2 + -\left(\frac{\Delta G}{p_0}\right)\frac{k_{on}(\sigma)}{k_{on}(\sigma) + k_{off}(\sigma)}\rho\right]V_c + K_p\epsilon_p^2 l_p^2$$

(S1.11)

Note the last term in Equation S1.11 which represents the micropost strain energy could be expressed in terms of $\sigma$ and $\epsilon$ since force balance dictates the stresses generated in the posts and cell must be the same (i.e., $\sigma = \sigma_p = \left(K_p\epsilon_p l_p\right)/A_c$), where $A_c$ is the cross-sectional area of the posts. Additionally, the bases of the microposts are fixed which gives a geometrical constraint leading to a mathematical relationship between the strain in the cell and the strain in the microposts: $\epsilon + 2\epsilon_p = 0$. Furthermore, if we assume that $V_c = \frac{1}{2}A_c l_p$ then the last term of Equation S1.11 can be rewritten as $K_p\epsilon_p^2 l_p^2 = -\sigma\epsilon V_c$. If we consider the temporal evolution of the metabolic potential and enforce the condition that it should satisfy the Second Law of Thermodynamics as it evolves towards steady state, i.e., $dR/dt \leq 0$, we find:

$$\frac{dR(\rho,\epsilon)}{dt} = \frac{\partial R}{\partial \rho}\frac{\partial \rho}{\partial t} + \frac{\partial R}{\partial \epsilon}\frac{\partial \epsilon}{\partial t} \leq 0$$

(S1.12)

The above Equation S1.12 can be written as:

$$\frac{dR(\rho,\epsilon)}{dt} = \frac{\partial \rho}{\partial t}\left(\epsilon + \beta(\rho - \rho_0) - \left(\frac{\Delta G}{p_0}\right)\frac{k_{on}(\sigma)}{k_{on}(\sigma) + k_{off}(\sigma)}\right)V_c + \frac{\partial \epsilon}{\partial t}(K\epsilon - \sigma + \rho)V_c$$

(S1.13)

We therefore choose the following time evolution laws for contractility and strain which will ensure that the Second Law of Thermodynamics (Equation (S1.12)) is always satisfied:

$$\frac{\partial \rho}{\partial t} = -k_\rho\frac{\partial R}{\partial \rho} = -k_\rho\left(\epsilon + \beta(\rho - \rho_0) - \left(\frac{\Delta G}{p_0}\right)\frac{k_{on}(\sigma)}{k_{on}(\sigma) + k_{off}(\sigma)}\right)V_c$$

(S1.14)

and,

$$\frac{\partial \epsilon}{\partial t} = -k_\epsilon\frac{\partial R}{\partial \epsilon} = -k_\epsilon(K\epsilon - \sigma + \rho)V_c$$

(S1.15)

where $k_\rho > 0$ and $k_\epsilon > 0$ are the kinetic constants that relate the rate of change of the metabolic potential with the corresponding change in contractility or strain, respectively. Given these relations, it follows that we may predict the steady state optimum configuration for the cell by minimizing the metabolic potential with respect to contractility and strain since at steady state ($\frac{\partial \rho}{\partial t} = 0$ and $\frac{\partial \epsilon}{\partial t} = 0$) Equations S1.14 and S1.15 yield the same result as minimizing the metabolic potential, i.e., setting ${\partial R}/{\partial \rho} = 0$ and ${\partial R}/{\partial \epsilon} = 0$. Minimizing with respect to contractility yields:

$$\frac{\partial R}{\partial \rho} = \left(\epsilon + \beta(\rho - \rho_0) - \left(\frac{\Delta G}{p_0}\right)\frac{k_{on}(\sigma)}{k_{on}(\sigma)+k_{off}(\sigma)}\right)V_c = 0,$$

(S1.16)

which is equivalent to Equation (S1.14) when $\frac{\partial \rho}{\partial t} = 0$. Similarly, minimizing with respect to strain yields:

$$\frac{\partial R}{\partial \epsilon} = (K\epsilon - \sigma + \rho)V_c = 0,$$

(S1.17)

which is equivalent to Equation (S1.15) when $\frac{\partial \epsilon}{\partial t} = 0$. Equation S1.17 is the condition for balance of forces; the stress generated by the cell is equal to the sum of active (contractility, $\rho$) and passive stresses ($K\epsilon$). Further, Equation S1.16 is the same as Equation S1.7 that predicted the dynamic steady state based on the kinetics of myosin motor contraction. *Hence, minimizing the metabolic potential of the cell with respect to contractility and strain is equivalent to predicting dynamic steady state on the basis of stress fiber assembly and disassembly consistent with the Second Law of Thermodynamics.*

By solving the system of equations (S1.16-S1.17), we find the following relations for the steady state optimum contractility and strain:

$$\rho = \left(\frac{\beta K}{K\beta - 1}\right)\rho_0 + \left(\frac{K\alpha - 1}{K\beta - 1}\right)\sigma$$

(S1.18(a))

and

$$\epsilon = \left(\frac{-\beta}{K\beta - 1}\right)\rho_0 + \left(\frac{\beta - \alpha}{K\beta - 1}\right)\sigma$$

(S1.18(b))

where for simplicity we have written these equations in terms of the parameter $\alpha$ as defined in Equation S1.22 $\left(\frac{k_{on}(\sigma)}{k_{on}(\sigma)+k_{off}(\sigma)}\left(\frac{\Delta G}{p_0}\right) = \alpha\sigma\right)$. In the limit of very rapid assembly (i.e. $k_\rho \to \infty$), the cell contractility is found to be:

$$0 = \epsilon + \beta(\rho - \rho_0) - \left(\frac{\Delta G}{p_0}\right)\frac{k_{on}(\sigma)}{k_{on}(\sigma) + k_{off}(\sigma)}$$

(S1.19)

By substituting equation (S1.19) into (S1.15) and rearranging, we obtain:

$$\frac{\hat{\sigma}}{\sigma_m} - \frac{\dot{\epsilon}}{\dot{\epsilon}_m} = 1$$

(S1.20)

The above is a linearized version of the Hill Law, where $\sigma_m$ is the stall stress and $\dot{\epsilon}_m$ is the maximum rate of contraction and are given by:

$$\dot{\epsilon}_m = k_\epsilon\left(\left(\frac{K\beta - 1}{\beta}\right)\epsilon + \rho_0\right)V_c$$

(S1.21a)

$$\hat{\sigma} = \sigma - \left(\frac{\Delta G}{p_0 \beta}\right) \frac{k_{on}(\sigma)}{k_{on}(\sigma) + k_{off}(\sigma)}$$

(S1.21b)

$$\sigma_m = \left(\frac{K\beta - 1}{\beta}\right)\epsilon + \rho_0$$

(S1.21c)

Thus, the Hill's Law is valid only when the contraction is very rapid, and in all other instances, the coupled equations (S1.14) and (S1.15) govern the rate of stress fiber assembly and deformation.

*1.2.* *<u>Steady state solution of a cell adhered to microposts:</u>*

To present a simple analysis without losing any physical insight, we assume the following simple functional stress dependence for the rate parameters that govern stress fiber assembly and disassembly:

$$\frac{k_{on}}{k_{off}} = \frac{\alpha\sigma}{1 - \alpha\sigma}$$

(S1.22)

which ensures that as the stress increases, the rate of assembly ($k_{on}$) increases while the rate of disassembly ($k_{off}$) decreases. The energy of ATP hydrolysis can therefore be written as:

$$U_{ATP} = -\alpha\sigma\rho V_c.$$

(S1.23)

We construct the expression for the metabolic potential as:

$$R(\rho, \epsilon) = \left(\frac{\beta}{2}\left(\rho - \rho_0\right)^2 + \rho\epsilon + \frac{1}{2}K\epsilon^2 - \alpha\sigma\rho\right) V_c + K_p l_p^2 \epsilon_p^2$$

(S1.24)

To find steady state values, we minimize by setting $\frac{\partial R}{\partial \epsilon} = 0$ and $\frac{\partial R}{\partial \rho} = 0$ and get the following two simultaneous equations in contractility and strain:

$$\frac{\partial R}{\partial \rho} = \beta\left(\rho - \rho_0\right) + \epsilon + \alpha \frac{K_p l_p \epsilon}{2A_c} = 0$$

(S1.25a)

$$\frac{\partial R}{\partial \epsilon} = (\rho + K\epsilon)V_c + \frac{K_p l_p^2 \epsilon}{2} = 0$$

(S1.25b)

Here we have made use of the relationship between $\sigma$ and $\epsilon$ explained in the previous SI Section 1.1.4: $\sigma = \sigma_p = \left(K_p \epsilon_p l_p\right)/A_c = -\frac{K_p l_p \epsilon}{2A_c}$. We can use Equation (S1.25b) to substitute for $\epsilon$ in the metabolic potential Equation (S1.24) and find how the different contributions change with $\rho$ to predict the steady state optimum. The following expressions are obtained:

$$U_{strain}^{cell+post} = \frac{1}{2}\frac{\rho^2 V_c}{\left(K + \frac{K_p l_p^2}{2V_c}\right)}$$

(S1.26)

$$U_{motor} = -\frac{\rho^2 V_c}{\left(K + \frac{K_p l_p^2}{2V_c}\right)}$$

(S1.27)

$$U_{ATP} = -\frac{\left(\frac{l_p}{2A_c}\right)\alpha K_p \rho^2 V_c}{\left(K + \frac{K_p l_p^2}{2V_c}\right)}$$

(S1.28)

These equations (S1.26)–(S1.28) along with the myosin binding energy (already in terms of $\rho$; $U_{binding} = \frac{\beta}{2}\left(\rho - \rho_0\right)^2)V_c$ and the metabolic potential $R$ are plotted in Figure 1D for two different values of post stiffness, $K_p$. The two different values of post stiffness represent a soft (~1.5 nN/µm) and a stiff (~30 nN/µm) system, which fall within the stiffness range reported for microposts fabricated to examine the effect of rigidity on mesenchymal cells (*69*). We observe the following trends:

- *Myosin Binding Energy*: The binding energy represents the energetic cost of increasing or decreasing the contractility from its quiescent value, $\rho_0$. In the case of cells adhered to stiff microposts, the upregulation of stress fiber assembly due to stress-activated signaling pathways (Figure 1A) leads to a large increase in the contractility from its quiescent state minimum value, $\rho_0$, and is associated with higher myosin binding energy (Figure 1D). By contrast, in cells adhered to soft microposts, the cytoskeletal stress developed is low and the increase in contractility from the quiescent state is not as large, leading to lower binding energy values (Figure 1D).
- *Mechanical Energy*: This represents the sum of the motor work and the elastic strain energy of the cell. The former is a measure of the potential energy of myosin motors, and by extension, the work done by motors in deforming the cytoskeleton and microposts. As cell contraction increases, the work done by the motors increases and is associated with a decrease in their potential energy. This mechanical work is stored as strain energy of the passive components including the cytoskeleton and microposts which increases with cell contractility. Note that when the cell deforms, the motor potential energy always decreases by twice the amount that the strain energy increases (Equations S1.26-S1.27). This arises from the fact that motors act as force dipoles, so any amount of strain energy they impart translates to their potential energy decreasing by double the amount. Thus, greater levels of contractility of the cell leads to lower potential energy and hence also lower overall mechanical energy for a given post stiffness (Figure 1D). On stiffer posts, the mechanical energy increases for all levels of contractility ($\rho$), leading to an upward shift in the curve (Figure 1D).
- *Energy of ATP hydrolysis*: While the mechanical energy and myosin binding energy is higher for cells adhered to stiffer microposts, more energy is available from ATP

hydrolysis in this case due to increased motor activity. On soft microposts, stresses experienced by the cell are small as is the stress-dependent increase in contractility since there is only a modest increase in the on-rate and modest decrease in the off-rate of stress fiber formation. By contrast, on stiff microposts, due to the high magnitude of cytoskeletal stress developed by the cell as it contracts, the upregulation of stress fiber assembly through mechanosensitive signaling pathways is enhanced (Figure 1A), leading to a much larger density of stress fibers and larger contractility at steady state. The magnitude of energy available through ATP hydrolysis is correspondingly larger on stiff microposts, as a larger amount of ATP is consumed to maintain the steady state contractility (Figure 1D).

Despite the unfavorable increases in binding energy and mechanical energy associated with establishing higher contractility, when coupled with ATP hydrolysis, which is an exothermic process, increases in contractility are favorable on stiffer posts. As discussed in SI Section 1.1.3, the stress-dependent kinetics of stress fiber assembly and disassembly determine the time-averaged amount of energy available from ATP hydrolysis. Stiffer environments favor the formation of stress fibers because these rates depend on cytoskeletal stress ($k_{on}(\sigma)$ increases with $\sigma$ and $k_{off}(\sigma)$ decreases with $\sigma$), establishing positive feedback between stress-activated signaling and contractility in our model. We find the minimum in the metabolic potential predicts an increasing level of steady state contractility with micropost stiffness driven by the larger rate of decrease in ATP hydrolysis energy with contractility on stiffer posts. While both the contractility on soft posts ($\rho_{soft}$) and the contractility achieved on stiff posts ($\rho_{stiff}$) are larger than the quiescent value ($\rho_0$), the level of steady state contractility is higher for cells on stiff posts ($\rho_{stiff} > \rho_{soft}$) due to more stress-dependent mechanosensitive upregulation of myosin motor recruitment (Figure 1D).

If we solve Equations (S1.25a) and (S1.25b) for $\epsilon$ and $\rho$ we obtain the following expressions for the steady state strain, contractility and stress (denoted as $\epsilon^*$,$\rho^*$ and $\sigma^*$respectively) as a function of post stiffness:

$$\epsilon^* = \frac{-\beta \rho_0}{K_p(\bar{\beta} - \bar{\alpha}) + K\beta - 1}$$

(S1.29(a))

$$\rho^* = \frac{\left(\frac{K_p l_p^2}{2V_c} + K\right)\beta \rho_0}{K_p(\bar{\beta} - \bar{\alpha}) + K\beta - 1}$$

(S1.29(b))

$$\sigma^* = \frac{-K_p \beta \rho_0}{K_p(\bar{\beta} - \bar{\alpha}) + (K\beta - 1)}\left(\frac{l_p}{2A_c}\right)$$

(S1.29(c))

where we define the normalized parameters $\bar{\beta} = \left(\frac{l_p^2}{2V_c}\right)\beta$ and $\bar{\alpha} = \left(\frac{l_p}{2A_c}\right)\alpha$. Next, we find the strain energy ($U_{strain}^{cell+post}$), energy of ATP hydrolysis ($U_{ATP}$) and motor work ($U_{motor}$) of the steady state configuration as a function of post stiffness by substituting equation (S1.29(b)) into equations (S1.26)-(S1.28). These equations are now given as:

$$U_{strain}^{*cell+post} = \frac{\beta^2 \rho_0^2 \left( \frac{K_p l_p^2}{2V_c} + K \right)}{2\left(K_p(\bar{\beta} - \bar{\alpha}) + K\beta - 1\right)^2} \tag{S1.30}$$

$$U_{ATP}^{*} = \frac{-\bar{\alpha} K_p \beta^2 \rho_0^2 \left( \frac{K_p l_p^2}{2V_c} + K \right) V_c}{\left(K_p(\bar{\beta} - \bar{\alpha}) + K\beta - 1\right)^2} \tag{S1.31}$$

$$U_{motor}^{*} = -\frac{\beta^2 \rho_0^2 \left( \frac{K_p l_p^2}{2V_c} + K \right)}{\left(K_p(\bar{\beta} - \bar{\alpha}) + K\beta - 1\right)^2} \tag{S1.32}$$

Note that the monotonicity of the trends in steady state strain energy and motor work depend on the relative stiffness between the cell and post. By examining the equation for the steady state cytoskeletal strain given in equation (S1.29a), we observe that in the stable parameter regime (where $\tilde{\beta} > \tilde{\alpha}$ and $\beta > 1/_K$), the denominator is an increasing function of post stiffness. Thus, the cytoskeletal strain decreases as post stiffness is increased (Figure S3A). Due to mechanosensitive pathways that upregulate stress fiber assembly on stiffer posts, the steady state contractility (Equation S1.29b, Figure S3B) and consequently the energy of ATP hydrolysis increases with post stiffness (Figure 1D). To better understand how the strain energy and the motor work of the steady state configuration vary with post stiffness, consider this function comprised of factors that appear in the expression for these energies:

$$f(K_p) = \frac{\left( \frac{K_p l_p^2}{2} + K V_c \right)}{\left( \frac{K_p}{2}(\bar{\beta} - \bar{\alpha}) + K\beta - 1 \right)^2} \tag{S1.33}$$

This is a non-monotonic function with respect to $K_p$ that changes trend at the value $K_p^*$ given by:

$$K_p^* = \frac{\left(K(2\bar{\alpha} - \bar{\beta}) - 1\right)}{(\bar{\beta} - \bar{\alpha})} \tag{S1.34}$$

If $K_p^* > 0$, the strain energy and motor work will show a non-monotonic trend: for $K_p < K_p^*$, the strain energy (motor work) increases (decreases) with matrix stiffness, while for $K_p > K_p^*$ they exhibit the opposite trend (Figure S3C). If $K_p^* < 0$, this ensures that the strain energy always decreases, and motor work always increases for physically realistic values of $K_p > 0$ (Figure S3D). This corresponds to the conditions $K < \frac{1}{2\bar{\alpha} - \bar{\beta}}$ or $\bar{\beta} > 2\bar{\alpha}$. To physically understand this, recall that the strain energy considered here is the sum of cell and micropost strain energies. With increasing post stiffness, the post strain energy increases, while that of the cell decreases. Thus, the variation of the total strain energy with post stiffness will depend on the relative strengths of these two contributions. At low post stiffnesses, if the cell is very stiff, the post strain energy will dominate because the cell will not experience much strain, resulting in an increasing trend in strain energy. However, as the post stiffness increases, the opposite becomes true and the cell strain energy begins to dominate, producing a decreasing trend. Therefore, to ensure a monotonic trend we find the cell stiffness, $K$, must be kept sufficiently low so that even on soft microposts the post strain energy does not dominate. The maximum value of $K$ is determined by the parameters governing the development of cell contractility because these determine how much stress the cell exerts against the microposts and therefore the post strain energy. For example, as $\alpha$ increases, the amount of contractility developed in the cell increases and the more the cell and microposts are strained. The maximum $K$ that guarantees a monotonic trend in strain energy will therefore decrease. Furthermore, the parameter $\beta$ which penalizes motor recruitment is accordingly subtracted from $\alpha$ in the expression for maximum value of $K$. We choose $K$ such that this condition $K < \frac{1}{2\bar{\alpha}-\bar{\beta}}$ is always satisfied.

*1.3* *<u>Estimation of model parameters based on comparisons against experiments:</u>*

Mitrossilis et al. measured the force exerted by myoblasts suspended between a rigid plate and a deformable cantilever, as a function of the stiffness of the cantilever (*35*) .Using the 1D model described so far, Shenoy et al. (*34*) calculated the force exerted by the cell on the microposts as, $F = \sigma r^2$, where $r$ is the size of the cell which we assume to be $10\ \mu m$. and $\sigma$ is the steady state cytoskeletal stress in the cell (given by Equation S1.22). By adopting this simple relation, the effect of the mechano-sensitive signaling parameters can be described by the single parameter, $\alpha$. Now the force exerted by cells on microposts can be calculated from our model as follows:

$$F = \sigma_{ss} r^2 = \frac{\bar{k}_p \bar{F}}{\bar{k} + \bar{k}_p} \tag{S1.35}$$

where,

$$\bar{k} = \left(\frac{l_p}{2A_c}\right) r \frac{K\beta - 1}{\bar{\beta} - \bar{\alpha}} \text{ and } \bar{F} = \frac{\beta \rho_0 r^2}{\bar{\beta} - \bar{\alpha}} \left(\frac{l_p}{2A_c}\right), \text{ and } \bar{k}_p = \left(\frac{l_p}{2A_c}\right) r k_p \tag{S1.36}$$

Here, $\rho_0$ is the quiescent state contractility, $\beta$ the chemical stiffness parameter, $K$ is the elastic modulus of the cytoskeleton and $\bar{k}_p$ is the normalized post stiffness. The parameter $\bar{k}$ is an effective stiffness that describes the ratio between increase in the cell force and the cell length $r\epsilon$, while $\bar{F}$ is the stall force exerted by the cell on the plates. The elastic modulus of myoblasts is estimated to be $1\ kPa$. In a previous work, the static contractility for cardiac tissues (*70*) contracting against deformable micro-cantilevers was estimated to be $0.5\ kPa$. Similar values were reported for the contraction of fibroblasts (*58*) confined in a 3D collagen matrix. Hence, we estimate the isotropic contractility $\rho_0$ to be $0.5\ kPa$. The effective stiffness $\bar{k}$ and the stall force $\bar{F}$ were estimated by curve fitting (using a Least Squares fitting method) the force predicted by the model (*34*) against experimental measurements of force exerted by myoblasts on the deformable plate (*35*). Using these two quantities, we estimate the biophysical parameters as follows:

$$\alpha = \frac{\bar{F}\left(\frac{l_p A_c}{V_c}\right) - \rho_0 r^2}{\bar{F}K - \bar{k} r \rho_0}, \text{ and } \beta = \frac{\bar{F}}{\bar{F}K - \bar{k} r \rho_0}$$

(S1.37)

The model parameters were then tuned so that the steady state force predicted as a function of post stiffness matches the experimental trend measured by Mitrossilis et al. (*35*), and the chemical stiffness and feedback parameters were estimated to be: $\beta = 2.77\ kPa^{-1}$ and $\alpha = 2.33\ kPa^{-1}$. These values are listed in Table S3 and serve as guidelines in estimating the parameters for subsequent simulation of mesenchymal cell behavior in 2D and 3D micro-environments as described in SI Section 2.

### 2. A 3D chemo-mechanical model for the metabolic potential of adherent cells

We now extend the model for the cell between two microposts presented in SI Section 1 to a more generalized 3D framework. Within this framework, we consider (a) contractile stress fibers, (b) the cell cytoskeleton, and (c) the nucleus, all of which are important mechanical components. While the cytoskeleton and nucleus represent passive elements, the stress fibers are active, contractile elements consisting of myosin motors engaged to actin filaments. First, we briefly describe these cellular components and the energetic contributions associated with them.

#### *2.1. Cellular components and mathematical models that describe their behavior within a 3D framework:*

*2.1.1. Stress fiber distribution, represented by the contractility tensor $\rho_{ij}$ and the cell cytoskeleton:* The cytoskeleton is comprised of actin filaments that transfer the contractile force generated by the cell to the extracellular matrix. The contraction of myosin motors attached to the actin filaments induces tensile stresses in the cytoskeleton which in turn are transmitted to an extracellular environment by adhesion proteins such as integrin (Figure 1A). The deformation of the cell cytoskeleton is represented by the strain tensor, $\epsilon_{ij}$, while the stresses induced in the cell due to actomyosin contraction are represented by the stress tensor, $\sigma_{ij}$. The quantities $\sigma_{kk}(\epsilon_{kk})$ and $\tilde{\sigma}_{ij}(\tilde{\epsilon}_{ij})$ represent the volumetric and deviatoric part of the cell stress (strain), respectively. In this model, the average density of engaged motors in the stress fibers per unit volume is a symmetric tensor $\rho_{ij}$ that emerges from the fact that we treat the myosin motors as force dipoles (Figure S1). Denoting the coordinates of the myosin heads as $x_j$ and $x_j + \Delta x_j$, $\left|F_i^{(k)}\right|$ as the magnitude of the force dipole and $\left|\Delta x_j\right|$ as the distance between the myosin II motor heads (Figure S1), the mechanical work performed by a single myosin motor can be written as:

$$W_{dipole} = F_i u_i(x_j + \Delta x_j) - F_i u_i(x_j)$$

(S2.1)

where $u_i(x_j)$ and $u_i(x_j + \Delta x_j)$ are the displacements of the cytoskeleton at the myosin head domains $x_j$ and $x_j + \Delta x_j$, respectively. Assuming that there are $N$ bound motors per unit volume, $V$, the total volumetric work done by all the motors can be written as:

$$F_{motor} = \frac{1}{V}\sum_{k=1}^{N} F_i u_i\left(x_j^{(k)} + \Delta x_j^{(k)}\right) - F_i u_i\left(x_j^{(k)}\right)$$

(S2.2)

Now, using the definition:

$$\partial_j u_i = \frac{u_i\left(x_j^{(k)} + \Delta x_j^{(k)}\right) - u_i\left(x_j^{(k)}\right)}{\Delta x_j^{(k)}}$$

(S2.3)

Equation S2.2 can be written as:

$$F_{motor} = \frac{1}{V}\sum_{k=1}^{N} F_i^{(k)}\, \Delta x_j^{(k)} \partial_j u_i$$

(S2.4)

Assuming small strain theory, the strain can be written as:

$$\epsilon_{ij} = \frac{1}{2}(\partial_j u_i + \partial_i u_j)$$

(S2.5)

and we can rewrite Equation S2.4 as:

$$F_{motor} = \rho_{ij}\epsilon_{ij}$$

(S2.6)

Here, we note the definition of the contractility tensor:

$$\rho_{ij} = \frac{1}{V}\sum_{k=1}^{N} F_i^{(k)}\, \Delta x_j^{(k)}$$

(S2.7)

which must be symmetric for the total dipole moment to vanish and whose components represent cell contractility in different directions. The volumetric part of the contractility tensor, $\rho_{kk}$, correlates with the density of motors engaged on stress fibers while the deviatoric part, $\tilde{\rho}_{ij}$ is a measure of their polarization. At any point in the cell, the principal direction corresponding to the largest eigenvalue of the contractility tensor represents the predicted orientation of motors and, by extension, stress fibers at that point.

*2.1.2. Nucleus, modeled as a poroelastic solid:*

The nucleus is comprised of a thin elastic layer (lamina) that encompasses the chromatin and other important sub nuclear organelles along with a nucleoplasm that constitutes the fluid phase. The nuclear response to stress primarily comprises fluid redistribution and fluid efflux/influx. This depends on the timescales of fluid flow. However, as fluid flow is very rapid compared to cell spreading, it can be effectively treated as a compressible elastic material (*71,72*). The nucleus is modelled as an elastic spheroidal inclusion encapsulated within the cell with elastic properties $K_{nuc}$ and $\mu_{nuc}$, denoting its bulk and shear modulus, respectively, while the strain experienced by the nucleus is denoted by $\epsilon_{ij}^{nuc}$. The nucleus is considered to be a spheroidal inclusion stiffer than the cell.

### 2.2. Metabolic potential of the cell within a 3D framework:

#### 2.2.1. Tensorial descriptions of the conservative contributions and energy of ATP hydrolysis:

To define the metabolic potential, we sum the various contributions of the cell and microenvironment. First, we define the components of the conservative energy, $U_c$, as: $\int_o^{V_c}(F_{binding} + F_{strain} + F_{motor})dV_c$. Here $V_c$ is the cell volume while $F_{binding}$, $F_{strain}$ and $F_{motor}$ are the myosin binding, strain energy, and motor work densities, respectively, defined at every point in the cell. The quiescent state contractility is an isotropic tensor with zero deviatoric components and a magnitude of $\rho_0$ which represents the density of motors in stress fibers in the absence of external stress as determined by a balance between the entropic cost of recruiting a free myosin motor to an actin filament and the enthalpic energy decrease achieved by bond formation between actin and myosin. We define the myosin binding energy density as follows:

$$F_{binding} = \frac{\beta}{6}(\rho_{kk}(\boldsymbol{x}) - 3\rho_0)^2 + \frac{\beta}{2}\tilde{\rho}_{ij}(\boldsymbol{x})\tilde{\rho}_{ij}(\boldsymbol{x}), \tag{S2.8}$$

where $\beta$ is a chemical stiffness parameter while $\rho_{kk}$ and $\tilde{\rho}_{ij}$ represent the volumetric and deviatoric part of the contractility tensor. In 3D, the spatial dependence of contractility, stress, and strain must be considered, as denoted by the spatial coordinate $\boldsymbol{x}$. Note that Equation S2.8 ensures that when the cell is in the quiescent state and $F_{binding} = 0$, the contractility, $\rho_{ij}$, adopts the quiescent state value ($\rho_{kk} = 3\rho_0$ and $\tilde{\rho}_{ij} = 0$). The density of motor work in the cell is given as: $F_{motor} = \rho_{ij}\epsilon_{ij}$, which can be written in terms of $\rho_{kk}$ and $\tilde{\rho}_{ij}$ as follows:

$$F_{motor} = \frac{1}{3}\rho_{kk}(\boldsymbol{x})\epsilon_{kk}(\boldsymbol{x}) + \tilde{\rho}_{ij}(\boldsymbol{x})\tilde{\epsilon}_{ij}(\boldsymbol{x}) \tag{S2.9}$$

The strain energy density represents the energy of deformation associated with the passive components, i.e., the cell, the nucleus, and the ECM, per volume. The strain energy of the ECM in the case of 3D collagen will be described fully in SI Section 2.2.6, but for now the strain energy densities can be written as:

$$F_{strain}^{cell} = \frac{\mathrm{K}_{cell}}{2}(\epsilon_{kk}(\boldsymbol{x}))^2 + \mu_{cell}\tilde{\epsilon}_{ij}(\boldsymbol{x})\tilde{\epsilon}_{ij}(\boldsymbol{x}) \tag{S2.10a}$$

$$F_{strain}^{nuc} = \frac{\mathrm{K}_{nuc}}{2}(\epsilon_{kk}^{nuc}(\boldsymbol{x}))^2 + \mu_{nuc}\tilde{\epsilon}_{ij}^{nuc}(\boldsymbol{x})\tilde{\epsilon}_{ij}^{nuc}(\boldsymbol{x}) \tag{S2.10b}$$

$$F_{strain}^{ECM} = -\frac{1}{3}\int_0^{\epsilon_{kk}(\boldsymbol{x})}\sigma_{kk}(\boldsymbol{x})d\epsilon_{kk}(\boldsymbol{x}) - \int_0^{\tilde{\epsilon}_{ij}}\tilde{\sigma}_{ij}(\boldsymbol{x})d\tilde{\epsilon}_{ij}(\boldsymbol{x}) \tag{S2.10c}$$

The total strain energy of the passive components is found by integrating these energy densities over the volumes of the respective components:

$$U_{strain} = \int F_{strain}^{cell} dV_c + \int F_{strain}^{nuc} dV_n + \int F_{strain}^{ECM} dV_{ECM}, \tag{S2.11}$$

where $V_n$ is the nucleus volume and $V_{ECM}$ is the volume of the ECM.

We motivate the energy density of ATP hydrolysis $F_{ATP}$ in a similar manner as the cell between two microposts model outlined in SI Section 1, by first establishing the relationship between $F_{ATP}$ and the stress-dependent parameters $k_{on,ij}(\sigma_{ij}(\boldsymbol{x}))$ and $k_{off,ij}(\sigma_{ij}(\boldsymbol{x}))$ that determine the rate of stress fiber assembly and disassembly, respectively. Within a 3D framework, $k_{on,ij}$ and $k_{off,ij}$ are now tensor quantities that allow for anisotropy in the rate of stress fiber formation when there is anisotropy in the stress field. In other words, the model accounts for situations where a cell experiences upregulated stress fiber assembly in, for example, the 11-direction compared to the 22-direction ($k_{on,11} > k_{on,22}$) when the cell experiences higher stress in that direction ($\sigma_{11} > \sigma_{22}$), The exact functional dependence of the components of $k_{on,ij}$ and $k_{on,ij}$ on the components of $\sigma_{ij}$ is informed by the biophysical mechanisms that underlie motor activation and stress fiber formation. To start, we assume that stress fiber kinetics are linearly proportional to the average stress at every point such that the volumetric parts of the on and off rates ($k_{on}^v$ and $k_{off}^v$) are proportional to the volumetric stress:

$$k_{on}^v(\sigma_{kk}(\boldsymbol{x})) = \frac{1}{3}\left(k_{on,11}(\sigma_{kk}(\boldsymbol{x})) + k_{on,22}(\sigma_{kk}(\boldsymbol{x})) + k_{on,33}(\sigma_{kk}(\boldsymbol{x}))\right) \tag{S2.12a}$$

$$k_{off}^v(\sigma_{kk}(\boldsymbol{x})) = \frac{1}{3}\left(k_{off,11}(\sigma_{kk}(\boldsymbol{x})) + k_{off,22}(\sigma_{kk}(\boldsymbol{x})) + k_{off,33}(\sigma_{kk}(\boldsymbol{x}))\right) \tag{S2.12b}$$

while the deviatoric parts ($\tilde{k}_{on,ij}(\tilde{\sigma}_{ij}(\mathbf{x}))$ and $\tilde{k}_{off,ij}(\tilde{\sigma}_{ij}(\mathbf{x}))$) allow for upregulated stress fiber formation rates in the direction of larger stresses. This captures the fact that mechanosignaling and an increase in the total number of active motors may arise from an increase in stress along any direction, but the cell will experience a greater rate of stress fiber formation in the direction of higher stress. The rate tensors for stress fiber assembly and disassembly can be written as the sum of these components:

$$k_{on,ij} = \frac{1}{3} k_{on}^v(\sigma_{kk}(\mathbf{x}))\delta_{ij} + \tilde{k}_{on,ij}(\tilde{\sigma}_{ij}(\mathbf{x})) \tag{S2.13a}$$

$$k_{off,ij} = \frac{1}{3} k_{off}^v(\sigma_{kk}(\mathbf{x}))\delta_{ij} + \tilde{k}_{off,ij}(\tilde{\sigma}_{ij}(\mathbf{x})) \tag{S2.13b}$$

However, we have previously shown that contractility regulated solely by volumetric (average) stress is insufficient to explain experimentally observed increases in cell contractility in elongated cell morphologies characterized by anisotropic stress distributions (39). This discrepancy arises from the formation of anisotropic adhesions aligned with polarized stress fibers, which establish bidirectional mechano-chemical feedback between cell shape and signaling (63). To model this process, we introduce a non-linear term to the rate of stress fiber assembly, $k^a_{on,ij}$, that accounts for increases in cell contractility but only when the stress field is anisotropic. This is readily accomplished by expressing tensor quantities in terms of their principal components to identify stress anisotropy independent of any coordinate system. The principal components of stress, $\sigma_i(\boldsymbol{x})$, reveal stress anisotropy exists whenever the largest principal component of stress is greater than the second largest principal component of stress. For example, if we assume $\sigma_1(\boldsymbol{x}) > \sigma_2(\boldsymbol{x}) > \sigma_3(\boldsymbol{x})$, then we assign $k^a_{on,i}$ a functional dependence on the term $\left(\frac{\sigma_1(x)}{\sigma_2(x)} - 1\right)$ such that $k^a_{on,i}$ is nonzero and leads to increased stress fiber assembly only when the stress fields are sufficiently anisotropic (when $\sigma_1(\boldsymbol{x}) > \sigma_2(\boldsymbol{x})$). To provide a generalized framework where principal stresses are not necessarily ordered, we include $\varepsilon_{ijk}$, the Levi-Civita tensor, and $\theta$, which represents a unit step function centered on 0, when denoting the energy of ATP hydrolysis:

$$F_{ATP}(\sigma_i(\boldsymbol{x})) = -\sum_i \frac{k_{on,i}(\sigma_i(x)) + k^a_{on,i}\left(\sum_{jk}|\varepsilon_{ijk}|\theta(\sigma_i(x)-\sigma_j(x))\theta(\sigma_j(x)-\sigma_k(x))\theta\left(\frac{\sigma_i(x)}{\sigma_j(x)}\right)\left(\frac{\sigma_i(x)}{\sigma_j(x)}-1\right)\sigma_i(x)\right)}{k_{off,i}(\sigma_i(x)) + k_{on,i}(\sigma_i(x)) + k^a_{on,i}\left(\sum_{jk}|\varepsilon_{ijk}|\theta(\sigma_i(x)-\sigma_j(x))\theta(\sigma_j(x)-\sigma_k(x))\theta\left(\frac{\sigma_i(x)}{\sigma_j(x)}\right)\left(\frac{\sigma_i(x)}{\sigma_j(x)}-1\right)\sigma_i(x)\right)} \frac{\Delta G}{p_0}\rho_i(\boldsymbol{x}) = -\sum_i \frac{k_{on,i}(\sigma_i(x)) + k^a_{on,i}(\overline{\sigma_a}(x))}{k_{off,i}(\sigma_i(x)) + k_{on,i}(\sigma_i(x)) + k^a_{on,i}(\overline{\sigma_a}(x))} \frac{\Delta G}{p_0}\rho_i(\boldsymbol{x}) \tag{S2.14}$$

where $k_{on,i}(\sigma_i(\boldsymbol{x}))$ and $k_{off,i}(\sigma_i(\boldsymbol{x}))$ are the rate tensors given by Equation S2.13 expressed in principal coordinates, and $\overline{\sigma_a}(\boldsymbol{x}) = \sum_{jk}|\varepsilon_{ijk}|\theta\left(\sigma_i(\boldsymbol{x}) - \sigma_j(\boldsymbol{x})\right)\theta\left(\sigma_j(\boldsymbol{x}) - \sigma_k(\boldsymbol{x})\right)\theta\left(\frac{\sigma_i(x)}{\sigma_j(x)}\right)\left(\frac{\sigma_i(x)}{\sigma_j(x)} - 1\right)\sigma_i(\boldsymbol{x})$. Note how the combination of factors in the term $k^a_{on,i}(\overline{\sigma_a}(\boldsymbol{x}))$ ensure upregulation is proportional to the largest principal stress and impose the condition that this term is nonzero only when anisotropy exists in the stress field.

Lastly, in calculating the total metabolic potential in 3D we account for changes in interfacial energy between the cell and matrix that result from changing cell shape and changing ECM composition. As cell surface area or ECM ligand density changes, the number of integrin-ligand interactions change as does the cell membrane tension, leading to changing interfacial energy. We account for these contributions by adding to the metabolic potential the term $\Gamma$ which depends on the cell shape factor $\Theta$, a variable that may be interpreted as any quantity relating cell shape, such as cell body aspect ratio. Now, in terms of principal components, we can write the metabolic potential of the cell, $R$, as:

$$\begin{aligned} R(\rho_i(\boldsymbol{x}), \epsilon_i(\boldsymbol{x}), \Theta) &= \int_{V_C}\left[\frac{\beta}{6}\sum_i(\rho_i - 3\rho_0)^2 + \frac{\beta}{2}\sum_i\left(\rho_i(\boldsymbol{x}) - \frac{\rho_1(\boldsymbol{x}) + \rho_2(\boldsymbol{x}) + \rho_3(\boldsymbol{x})}{3}\right)^2 \right.\\ &+ \sum_i \rho_i(\boldsymbol{x})\epsilon_i(\boldsymbol{x}) \\ &\left. - \sum_i \frac{k_{on,i}(\sigma_i(\boldsymbol{x})) + k^a_{on,i}(\overline{\sigma_a}(\boldsymbol{x}))}{k_{off,i}(\sigma_i(\boldsymbol{x})) + k_{on,i}(\sigma_i(\boldsymbol{x})) + k^a_{on,i}(\overline{\sigma_a}(\boldsymbol{x}))}\frac{\Delta G}{p_0}\rho_i(\boldsymbol{x})\right] dV_c \\ &+ U_{strain}(\rho_i(\boldsymbol{x}), \epsilon_i(\boldsymbol{x}), \Theta) + \Gamma(\Theta) \end{aligned}$$

(S2.15)

Note that for convenience we refer to the sum of all terms in Equation S2.15 except $\Gamma(\Theta)$ as the *volumetric part of the metabolic potential* in order to highlight the competition between $\Gamma(\Theta)$ and all other terms in determining cell shape.

To predict the steady state optimum for a cell using this 3D implementation, the metabolic potential must be minimized with respect to contractility, $\rho_i(x)$, strain, $\epsilon_i(x)$, and cell shape, $\Theta$. In our simulation procedure we accomplish this by first taking the functional derivatives of $R$ with respect to $\rho_i(x)$ and $\epsilon_i(x)$ to generate PDEs that are solved using finite element simulations for fixed cell shape. We then repeat this process for incrementally changing cell aspect ratio to determine the cell shape which produces minimum $R$. For example, minimizing $R$ with respect to contractility and strain generates the following conditions:

$$\frac{\delta R\big(\rho_i(\boldsymbol{x}),\epsilon_i(\boldsymbol{x}),\Theta\big)}{\delta\rho_i(\boldsymbol{x})} = \frac{\delta U_c\big(\rho_i(\boldsymbol{x}),\epsilon_i(\boldsymbol{x}),\Theta\big)}{\delta\rho_i(\boldsymbol{x})} - \int\sum_i \frac{k_{on,i}(\sigma_i(\boldsymbol{x})) + k_{on,i}^a(\overline{\sigma_a}(\boldsymbol{x}))}{k_{off,i}(\sigma_i(\boldsymbol{x})) + k_{on,i}(\sigma_i(\boldsymbol{x})) + k_{on,i}^a(\overline{\sigma_a}(\boldsymbol{x}))}\frac{\Delta G}{p_0}dV_c = 0$$

(S2.16a)

$$\frac{\delta R\big(\rho_i(\boldsymbol{x}),\epsilon_i(\boldsymbol{x}),\Theta\big)}{\delta\epsilon_i(\boldsymbol{x})} = \frac{\delta U_c\big(\rho_i(\boldsymbol{x}),\epsilon_i(\boldsymbol{x}),\Theta\big)}{\delta\epsilon_i(\boldsymbol{x})} = 0$$

(S2.16b)

where $U_c$ is the sum of the conservative energies included in the volumetric part of the metabolic potential (everything except ATP hydrolysis energy). We show in SI Section 2.2.3 that, similar to the 1D micropost model, the above is equivalent to the steady state condition predicted by considering the kinetics of stress fiber assembly and disassembly in a 3D formulation.

*2.2.2. <u>Strain and contractility in 3D assuming a simplified dependence of the rate parameters on stress:</u>*

Similar to the cell between two microposts model (see Equation S1.22), we can assume the following simplified form for the energy of ATP hydrolysis of the cell but now account for a full tensorial implementation:

$$F_{ATP} = -\alpha_v\sum_i \sigma_i(\boldsymbol{x})\rho_i(\boldsymbol{x}) - \alpha_d\sum_i\left(\sigma_i(\boldsymbol{x}) - \frac{\sigma_1(\boldsymbol{x})+\sigma_2(\boldsymbol{x})+\sigma_3(\boldsymbol{x})}{3}\right)\sum_i\left(\rho_i(\boldsymbol{x}) - \frac{\rho_1(\boldsymbol{x})+\rho_2(\boldsymbol{x})+\rho_3(\boldsymbol{x})}{3}\right) - \alpha_a\sum_i \overline{\sigma_a}(\boldsymbol{x})\rho_i(\boldsymbol{x})$$

(S2.17)

$\alpha_v$ represents the intensity of upregulation in the number of stress fibers per unit volume due to volumetric stress. For the same increase in volumetric stress, $\sigma_i$, higher values of $\alpha_v$ lead to a greater number of stress fibers per unit volume, and consequently higher contractility. Similarly, $\alpha_d$ represents the intensity of polarization induced due to the presence of deviatoric stresses. A very small value of $\alpha_d$ leads to an unpolarized cell. The third term in S2.17 represents upregulation in stress fiber assembly due to stress anisotropy, and $\alpha_a$ is the parameter controlling the intensity of this upregulation. If one assumes that the principal stresses are always ordered such that $\sigma_1 \geq \sigma_2 \geq \sigma_3$ and $\sigma_1 \geq \sigma_2 > 0$, then the expression can be simplified to:

$$F_{ATP} = -\alpha_v \sum_i \sigma_i(\boldsymbol{x})\rho_i(\boldsymbol{x}) - \alpha_d \sum_i \left(\sigma_i(\boldsymbol{x}) - \frac{\sigma_1(\boldsymbol{x}) + \sigma_2(\boldsymbol{x}) + \sigma_3(\boldsymbol{x})}{3}\right) \sum_i \left(\rho_i(\boldsymbol{x}) - \frac{\rho_1(\boldsymbol{x}) + \rho_2(\boldsymbol{x}) + \rho_3(\boldsymbol{x})}{3}\right) - \alpha_a \sigma_a(\boldsymbol{x})\rho_1(\boldsymbol{x})$$

(S2.18)

where $\sigma_a(\boldsymbol{x}) = \left(\frac{\sigma_1(\boldsymbol{x})}{\sigma_2(\boldsymbol{x})} - 1\right)\sigma_1(\boldsymbol{x})$. To further simplify, we can also assume the strength of upregulation based on volumetric and deviatoric stresses are the same ($\alpha_v = \alpha_d = \alpha$), and the rate parameters can be related to stress as follows:

$$\sum_i \frac{k_{on,i}(\sigma_i(\boldsymbol{x})) + k_{on,i}^a(\overline{\sigma_a}(\boldsymbol{x}))}{k_{on,i}(\sigma_i(\boldsymbol{x})) + k_{on,i}^a(\overline{\sigma_a}(\boldsymbol{x})) + k_{off,i}(\sigma_i(\boldsymbol{x}))} \frac{\Delta G}{p_0} = \sum_i \alpha\sigma_i(\boldsymbol{x}) + \alpha_a \sigma_a(\boldsymbol{x})$$

(S2.19)

Simplifying the energy of ATP hydrolysis using Equation S2.19 and minimizing the metabolic potential with respect to strain and contractility (Equation S2.16), we obtain the following expressions for the steady state contractility and stress fields:

$$\rho_{kk}(\boldsymbol{x}) = \left(\frac{3\beta}{\beta - \alpha}\right)\rho_0 + \left(\frac{\alpha_a}{\beta - \alpha}\right)\sigma_a(\boldsymbol{x}) + \left(\frac{3K\alpha - 1}{\beta - \alpha}\right)\epsilon_{kk}(\boldsymbol{x})$$

(S2.20a)

$$\tilde{\rho}_{ij}(\boldsymbol{x}) = \left(\frac{2\mu\alpha - 1}{\beta - \alpha}\right)\tilde{\epsilon}_{ij}(\boldsymbol{x})$$

(S2.20b)

$$\sigma_{kk}(\boldsymbol{x}) = \left(\frac{3\beta}{\beta - \alpha}\right)\rho_0 + \left(\frac{\alpha_a}{\beta - \alpha}\right)\sigma_a(\boldsymbol{x}) + \left(\frac{3K\beta - 1}{\beta - \alpha}\right)\epsilon_{kk}(\boldsymbol{x})$$

(S2.21a)

$$\tilde{\sigma}_{ij}(\boldsymbol{x}) = \left(\frac{2\mu\beta - 1}{\beta - \alpha}\right)\tilde{\epsilon}_{ij}(\boldsymbol{x})$$

(S2.21b)

*2.2.3. <u>Energy of ATP hydrolysis and steady state dissipation rate governed by the kinetics of motor engagement:</u>*

Adherent cells remain contractile over long timescales (stress fiber disassembly timescales are of the order of several minutes to hours (*73*)). As the timescale of individual motor binding events is of the order of seconds (*74*), there must be a continuous cycle of myosin motor binding and motor unbinding that is responsible for force generation. Myosin contraction is accompanied by ATP hydrolysis, and, thus, in a time-averaged sense, ATP is continuously consumed (during the myosin work cycle) to maintain contractility. The rate of stress fiber assembly is denoted by a tensor $J_{on}$, whose individual components ($J_{on,ij}$) represent the time rate of change of the contractility ($\rho_{ij}$) in different directions. Thus, the constant energy input required to sustain this contractile state is $J_{on,ij}\,\frac{\Delta G}{p_0}$, where $\Delta G$ is the energy released by ATP hydrolysis and $p_0$ is the dipole strength of an individual myosin motor. Similarly, the rate of stress fiber disassembly is denoted by the tensor $J_{off,ij}$. The rate of change of the individual components of the contractility can be written in principal components as:

$$\frac{\partial \rho_i(\boldsymbol{x})}{\partial t} = J_{on,i}\,(\boldsymbol{x}) + J^a_{on,i}(\boldsymbol{x}) + J_{off,i}\,(\boldsymbol{x}) \tag{S2.22}$$

Note here we have split $J_{on,ij}$ into two parts to explicitly account for the rate of stress fiber assembly associated with feedback dependent on the average of stress ($J_{on,ij}$) and dependent on stress anisotropy ($J^a_{on,ij}$). A change in cell contractility involves a change in the conservative free energy density of the cell, $F_c$. $\frac{\partial F_c}{\partial \rho_i}$ represents the change in the conservative energy upon an incremental change in contractility, or equivalently, an incremental change in stress fiber density. The dissipation rate represents the heat dissipated per unit time per unit volume by motor activity and is calculated as the difference between the energy available through ATP hydrolysis and the energy converted into conservative forms. In terms of principal components, the dissipation rate per unit volume can be written as:

$$\begin{aligned}
\dot{D}(\boldsymbol{x}) = \sum_{i=1}^{3} &\underbrace{J_{on,i}(\boldsymbol{x})\frac{\Delta G}{p_0} + J^a_{on,i}(\boldsymbol{x})\frac{\Delta G}{p_o}}_{\substack{\textit{Energy available}\\ \textit{from ATP hydrolysis}}} \underbrace{- J_{on,i}(\boldsymbol{x})\frac{\delta F_c(\boldsymbol{x})}{\delta \rho_i(\boldsymbol{x})} - J^a_{on,i}(\boldsymbol{x})\frac{\delta F_c(\boldsymbol{x})}{\delta \rho_i(\boldsymbol{x})}}_{\substack{\textit{Conservative energy}\\ \textit{change upon}\\ \textit{stress fiber assembly}}} \\
&- \underbrace{J_{off,i}(\boldsymbol{x})\frac{\delta F_c(\boldsymbol{x})}{\delta \rho_i(\boldsymbol{x})}}_{\substack{\textit{Conservative energy}\\ \textit{change upon stress}\\ \textit{fiber disassembly}}} \\
= \sum_i &-J_{on\,,i}\,(\boldsymbol{x})\left(\frac{\delta F_c(\boldsymbol{x})}{\delta \rho_i(\boldsymbol{x})} - \frac{\Delta G}{p_0}\right) - J^a_{on,i}(\boldsymbol{x})\left(\frac{\delta F_c(\boldsymbol{x})}{\delta \rho_i(\boldsymbol{x})} - \frac{\Delta G}{p_0}\right) - J_{off,ij}\,(\boldsymbol{x})\frac{\delta F_c(\boldsymbol{x})}{\delta \rho_i(\boldsymbol{x})}
\end{aligned} \tag{S2.23}$$

In order to satisfy the second law of thermodynamics, i.e. $\dot{D} > 0$, we can choose the following forms for the rates:

$$J_{on,i}(\boldsymbol{x}) = -k_{on,i}(\sigma_i(\boldsymbol{x}))\left(\frac{\delta F_c(x)}{\delta\rho_i(x)} - \frac{\Delta G}{p_0}\right); \quad J^a_{on,i}(\boldsymbol{x}) = k^a_{on,i}(\overline{\sigma_a}(\boldsymbol{x}))\left(\frac{\delta F_c(x)}{\delta\rho_i(x)} - \frac{\Delta G}{p_0}\right);$$
$$J_{off,ij}(\boldsymbol{x}) = -k_{off,i}(\sigma_i(\boldsymbol{x}))\frac{\delta F_c(x)}{\delta\rho_i(x)}$$
(S2.24)

In the above equation, $k_{on,i}$, $k^a_{on,i}$, and $k_{off,i}$ are kinetic parameters (with units $Pa \cdot s^{-1}$) that govern the stress-regulated assembly and disassembly of stress fibers along different directions. At steady state $\frac{\partial\rho_i(x)}{\partial t} = 0$, and Equations S2.22 and S2.24 may be combined to yield the same result as Equation S2.16, revealing that *minimizing the metabolic potential is equivalent to the steady state obtained by considering stress fiber kinetics*.

At steady state, Equations S2.22 – S2.24 yield the following expression for the steady state heat dissipation per volume:

$$\dot{D}(\boldsymbol{x}) = \sum_{i=1}^{3} \frac{k_{on,i}(\sigma_i(\boldsymbol{x}))k_{off,i}(\sigma_i(\boldsymbol{x})) + k^a_{on,i}(\overline{\sigma_a}(\boldsymbol{x}))k_{off,i}(\sigma_i(\boldsymbol{x}))}{k_{off,i}(\sigma_i(\boldsymbol{x})) + k_{on,i}(\sigma_i(\boldsymbol{x})) + k^a_{on,i}(\overline{\sigma_a}(\boldsymbol{x}))} \frac{(\Delta G)^2}{{p_0}^2}$$
(S2.25)

the total steady state heat dissipated per unit time, denoted by $\dot{D}_{cell}$ is then found by integrating $\dot{D}$ over the cell volume as:

$$\dot{D}_{cell} = \int_{V_c} \sum_{i=1}^{3} \frac{k_{on,i}(\sigma_i(\boldsymbol{x}))k_{off,i}(\sigma_i(\boldsymbol{x})) + k^a_{on,i}(\overline{\sigma_a}(\boldsymbol{x}))k_{off,i}(\sigma_i(\boldsymbol{x}))}{k_{off,i}(\sigma_i(\boldsymbol{x})) + k_{on,i}(\sigma_i(\boldsymbol{x})) + k^a_{on,i}(\overline{\sigma_a}(\boldsymbol{x}))} \frac{(\Delta G)^2}{{p_0}^2} dV_c$$
(S2.26)

Note that, at steady state, there is no change in the net amount of stress fibers, as the assembly and disassembly rates are balanced (dynamic equilibrium). Thus, the contractility does not change with time at steady state. However, ATP is constantly consumed to maintain assembled stress fibers, and all the energy released by ATP consumption is dissipated as heat. Thus, Equation S2.26 not only yields the amount of heat released by a cell at steady state but also reveals the amount of power required to maintain the cell's contractile apparatus.

We may find real estimates of cell power if good estimates are known for the rates $k_{on_i}(\sigma_i(\boldsymbol{x}))$ and $k_{off_i}(\sigma_i(\boldsymbol{x}))$. For simplicity, we may assume $k_{on_i}(\sigma_i(\boldsymbol{x}))$ follows a linear relationship with stress such that $k_{on_i}(\sigma_i(\boldsymbol{x})) = k_{on_{0,i}} + c\sigma_i(\boldsymbol{x})$, where $k_{on_{0,i}}$represents stress fiber assembly rate in the absence of stress, $c$ is a constant relating the kinetics of stress fiber assembly to stress-dependent signaling. Here, we assume that stress fiber assembly rates in the absence of stress are very small i.e. $k_{on_{0,i}} \ll c\sigma_i(\boldsymbol{x})$, and hence we can simplify the equation to the form: $k_{on_i}(\sigma_i(\boldsymbol{x})) \approx c\sigma_i(\boldsymbol{x})$. This is a reasonable assumption supported by the fact that stress fibers are observed in vivo only under conditions of high mechanical tension (*75*). Similarly, $k^a_{on,i}$ is a rate parameter that depends on stress anisotropy, so we may assume it is of the form $k^a_{on,i}(\overline{\sigma_a}(\boldsymbol{x})) \approx c\left(\frac{\sigma_1(x)}{\sigma_2(x)} - 1\right)\sigma_1(\boldsymbol{x}) = c\sigma_a(\boldsymbol{x})$. From Equation S2.19, $k_{off_i}(\sigma_i(\boldsymbol{x}))$ can then be found as: $k_{off_i}(\sigma_i(\boldsymbol{x})) = \frac{c\sigma_i(x)+c\sigma_a(x)}{\alpha\sigma_i(x)+\alpha_a\sigma_a(x)}\left(\frac{\Delta G}{p_0} - (\alpha\sigma_i(\boldsymbol{x}) + \alpha_a\sigma_a(\boldsymbol{x}))\right)$. Altogether, Equation S2.26 simplifies to:

$$\dot{D}_{cell}(\boldsymbol{x}) = \int_{V_c} \sum_{i=1}^{3} \left(c\sigma_i(\boldsymbol{x}) + c\sigma_a(\boldsymbol{x})\right)\left(\frac{\Delta G}{p_0} - \left(\alpha\sigma_i(\boldsymbol{x}) + \alpha_a\sigma_a(\boldsymbol{x})\right)\right)\frac{\Delta G}{p_0}\, dV_c$$

(S2.27)

If we use the same value for the parameters $\alpha$ and $\alpha_a$ as determined and used elsewhere in our model (Table S3, SI Section 2.3), then it is only necessary to estimate $p_0$, $\Delta G$, and $c$ to estimate the power. Literature reports the energy of ATP hydrolysis in physiological settings as ~$12k_BT$, (~30 kJ/mol) and the dipole strength of a single myosin motor as ~5e-20 J (*62*). We estimate the value of $c$ here as 0.0008 $s^{-1}$, which represents the rate of stress fiber assembly (deduced by examining experimentally reported time lapse microscopy images that show stress fiber formation at around 15-20 min in human U2OS cells (*73*)). Using these values (summarized in Table S6), we extract from our finite element models the steady state cell power ($\dot{D}_{cell}$) by first calculating $D(\dot{\boldsymbol{x}})$ (Equation S2.25) at every point in the cell and then integrating over the cell volume. For example, we calculate the steady state power for a cell encapsulated in a collagen matrix of density 1 mg/mL is ~**2.2 fW**. Since **1 femtoWatt of power is roughly equivalent to 20,000 ATP molecules consumed per second** (62), this means the cell must consume **~44000 ATP molecules per second** to maintain this steady state configuration.

*2.2.4. <u>Hill's Law attained as a limiting case by considering the dynamics underlying metabolic potential</u>:*

As in the case of a cell on microposts, we now consider the temporal evolution of the metabolic potential, and to ensure that the Second Law of Thermodynamics is satisfied, we have the following evolution laws for the contractility and strain fields:

$$\frac{\partial \rho_{kk}(\boldsymbol{x})}{\partial t} = -k_\rho^v \int \left(\frac{1}{3}\epsilon_{kk}(\boldsymbol{x}) + \frac{1}{3}\beta(\rho_{kk}(\boldsymbol{x}) - 3\rho_0) - \frac{1}{3}\alpha\sigma_{kk}(\boldsymbol{x}) - \alpha_a\sigma_a(\boldsymbol{x})\right) dV_c$$

(S2.28a)

$$\frac{\partial \tilde{\rho}_{ij}(\boldsymbol{x})}{\partial t} = -k_\rho^d \int \left(\tilde{\epsilon}_{ij}(\boldsymbol{x}) + \beta\tilde{\rho}_{ij}(\boldsymbol{x}) - \alpha\tilde{\sigma}_{ij}\right) dV_c$$

(S2.28b)

$$\frac{\partial \epsilon_{kk}(\boldsymbol{x})}{\partial t} = -k_\epsilon^v \int \left(K\epsilon_{kk}(\boldsymbol{x}) + \frac{1}{3}\rho_{kk}(\boldsymbol{x}) - \frac{1}{3}\sigma_{kk}(\boldsymbol{x})\right) dV_c$$

(S2.29a)

$$\frac{\partial \tilde{\epsilon}_{ij}(\boldsymbol{x})}{\partial t} = -k_\epsilon^d \int \left(2\mu\tilde{\epsilon}_{ij}(\boldsymbol{x}) + \tilde{\rho}_{ij}(\boldsymbol{x}) - \tilde{\sigma}_{ij}(\boldsymbol{x})\right) dV_c$$

(S2.29b)

The above equation shows that the contractility increases with stress and that, while the contractility is isotropic for spherical cells, elongated cells experience an anisotropic stress field that polarizes the contractility as well. This is captured in the above equation by the presence of non-zero deviatoric terms (or anisotropy) for stress states that are not purely hydrostatic. In the limit that the rate of stress-fiber assembly is very fast (i.e. $k_\rho^v$, $k_\rho^d \rightarrow \infty$), the terms in the brackets of equation (S2.28a) vanish, and we find an expression for the volumetric part of the contractility as:

$$\rho_{kk}(\boldsymbol{x}) = 3\rho_0 + \frac{\alpha\sigma_{kk}(\boldsymbol{x})}{\beta} + \frac{3\alpha_a\sigma_a(\boldsymbol{x})}{\beta} - \frac{\epsilon_{kk}(\boldsymbol{x})}{\beta} \tag{S2.30}$$

Now, substituting above equation in (S2.29a) and simplifying, we obtain:

$$\frac{\sigma_{kk}(\boldsymbol{x})}{\sigma_m} - \frac{\dot{\epsilon}_{kk}(\boldsymbol{x})}{\dot{\epsilon}_m} = 1 \tag{S2.31}$$

Here $\sigma_m$ can be considered a 'stall stress' while $\dot{\epsilon}_m$ is the maximum rate of stress fiber assembly. These are given as:

$$\dot{\epsilon}_m = \frac{k_\epsilon^v V_c}{3}\left(\frac{3K\beta-1}{\beta-\alpha}\epsilon_{kk}(\boldsymbol{x}) + \frac{3\beta}{\beta-\alpha}\rho_0 + \frac{3\alpha_a}{\beta-\alpha}\sigma_a(\boldsymbol{x})\right),\ \sigma_m = \frac{3K\beta-1}{\beta-\alpha}\epsilon_{kk}(\boldsymbol{x}) + \frac{3\beta}{\beta-\alpha}\rho_0 + \frac{3\alpha_a}{\beta-\alpha}\sigma_a(\boldsymbol{x}) \tag{S2.32}$$

Equation S2.31 is a linear version of the Hill equation. Thus, Hill's relation is only valid when stress-fiber assembly rates are very rapid and for all other cases, the coupled equations need to be used to determine the time evolution of contractility and strain.

#### 2.2.5. *Interfacial Energy:*

We consider two main energetic contributions at the cell membrane that influence steady state cell shape separately from the volumetric part of the metabolic potential. These include cell membrane tension, $\gamma_0$, a surface tension-like term that imposes an energetic penalty to cell surface area increases, and receptor-ligand adhesive binding interactions, $\gamma_1$. The latter represents the reduction in energy achieved due to bond formation between receptors such as integrins at the cell-matrix interface and ligands present in the ECM. The relative strengths of these two quantities determine whether the interfacial energy of the cell increases or decreases with cell surface area. We can estimate the binding contribution $\gamma_1$ as follows:

$$\gamma_1 = \underbrace{\eta}_{\substack{fraction\\ of\\ bound\ receptors}} * \underbrace{d_{ad}}_{\substack{Receptor\\ density\\ per\ unit\ area}} * \underbrace{E_{RL}}_{\substack{energy\ of\ a\ single\\ receptor-ligand\ bond}} \tag{S2.33}$$

where $\eta$ is a parameter that represents the fraction of receptors on the cell membrane that are bound to ligands, $d_{ad}$ is the number of receptors available per unit cell surface area and $E_{RL}$ is the energy released due to bond formation between a single receptor and ligand. For the same cell spread area, the adhesive energy decreases in high density collagen due to the increased availability of ligands per unit area and consequent reduction in energy due to increased number of integrin-ligand interactions (increased $\eta$). When the ligand density is fixed but cell spread area increases, the number of receptor ligand bonds, $\eta$, again increases, leading to a decrease in the adhesive energy (Figure 2A).

As the membrane tension resists an increase in cell surface area, the energy penalty associated with cell elongation is given by $\gamma_0 S$, where $S$ is the surface area of the cell. Similarly, with increasing surface area, energy is released due to integrin-ligand bond formation, and this energy is calculated as $-\gamma_1 S$ . Hence, depending on the relative magnitudes of these two quantities, the total interfacial energy $\left((\gamma_0 - \gamma_1)S\right)$ will increase or decrease with cell surface area, $S$. Studies report typical membrane tension values for the cell are on the order of $\sim 10^{-4} - 10^{-5} N/m$ (*76*), even when measured near focal adhesions of an adhered cell (*77*). We roughly estimate the energy of the receptor-ligand energy $\gamma_1$ as follows: the receptor density was experimentally measured in HeLa cervical cancer cells to be $\sim 550\ \mu m^{-2}$ using a fluorescence correlation spectroscopy method (*78*). Assuming the energy associated with integrin-ligand bond (*79*) formation to be $5kT$ (where $k$ is the Boltzmann constant and $T$ is the standard temperature $298K$), and the fraction of bound receptors $\eta$ to be 0.5, we estimate $\gamma_1$ as $\sim 10^{-6} N/m$ using Equation S2.33, which is around 2 order of magnitude smaller than $\gamma_0$. Since the cell adhesion energy, $\gamma_1$, is around two orders of magnitude smaller than the membrane tension, $\gamma_0$, the effective interfacial energy is a positive quantity and increases with cell surface area. Cells develop larger and more stable focal adhesions on stiffer substrates, even if the number of available ligands on the substrate remains constant (*47*). This suggests the fraction of bound receptors increases with stiffness in 2D, and therefore, the total interfacial energy, $\Gamma_{2D} = \left((\gamma_0 - \gamma_1(E))S\right)$, is a function that decreases with substrate stiffness, $E$. However, in 3D collagen, reports suggest cell-matrix adhesion formation is coupled to the ability of the cell to degrade the matrix, and this decreases with matrix density (*44*). Therefore, total interfacial energy in 3D collagen must also include an increasing function of matrix density, $d$, to account for this, $\Gamma_{3D} = \left((\gamma_0 - \gamma_1(d) + \gamma_2(d))S\right)$. To simplify, we assume $\gamma_2 > \gamma_1$ such that $\Gamma_{3D} = \left((\gamma_0 + \gamma_2(d))S\right)$.

Following these expressions, in 2D simulations we account for cell membrane tension by considering the membrane tension both along the cell-air (CA) interface and along the cell-matrix (CM) interface ($\gamma_0 = \gamma^{CA}S^{CA} + \gamma^{CM}S^{CM}$) in addition to the binding energy of the cell-matrix interface ($\gamma_1 = \gamma^{CM}(E)S^{CM}$) so that the total interfacial energy is calculated as $\Gamma_{2D} = (\gamma^{CA}S^{CA} + \gamma^{CM}(E)S^{CM}) = (\gamma^{CA}S^{CA} + \gamma^{CM}(1 - f^{CM}(E))S^{CM})$. For simulations in 3D collagen, $\gamma_0 = \gamma S$ and $\gamma_2 = \gamma f(d) S$ so that the total interfacial energy is calculated as $\Gamma_{3D} = (\gamma(1 + f(d))S)$ The values used for the interfacial energy parameters in our simulations were determined through a sensitivity analysis (Figure S8) and are listed in Table S3.

*2.2.6. Fibrous constitutive model for 3D collagen:*

The mechanical behavior of collagen is modelled using a non-linear, anisotropic constitutive law for fibrous materials (*43*). The collagen matrix is comprised of two families of fibers: an isotropic distribution of randomly aligned fibers and a set of fibers that are aligned along the direction of maximum principal stretch (Figure S4A). The strain energy density of the collagen matrix at every point $\boldsymbol{x}$ is given as the sum of the two:

$$F_{strain}^{collagen}(\boldsymbol{x}) = W^I(\boldsymbol{x}) + W^f(\boldsymbol{x}) \tag{S2.34}$$

Here, $W^I$ represents the strain energy of the isotropic, randomly aligned fibers which are modelled as linearly elastic:

$$W^I(\boldsymbol{x}) = \frac{K_{ECM}}{2}(\epsilon_{kk}^{ECM}(\boldsymbol{x}))^2 + \mu_{ECM}\left(\tilde{\epsilon}_{ij}^{ECM}(\boldsymbol{x})\right)^2$$

(S2.35)

where, $\mu_{ECM}$ and $K_{ECM}$ are the initial shear and bulk moduli of the matrix, respectively, and $\epsilon_{ij}^{ECM}$is the strain in the matrix. The strain energy of the aligned fibers can be written as:

$$W^f(\boldsymbol{x}) = \sum_{a=1}^{3} f(\lambda_a(\boldsymbol{x}))$$

(S2.36)

where $\lambda_a$ are the principal stretches, and the energy function $f(\lambda_a)$ is chosen such that (i) the energy density contribution from the aligned fibers, $W^f$ vanishes below a critical value of tensile stretch, and (ii) the matrix stiffens only in the direction of (tensile) principal stretches when $\lambda_a$ is larger than the (tensile) stretch critical value, $\lambda_c$ (Figure S4B). The following form is used for $f(\lambda_a)$:

$$\frac{\delta f(\lambda_a(\boldsymbol{x}))}{\delta \lambda_a} = \begin{cases} 0, & \lambda_a < \lambda_1 \\ E_f \dfrac{\left(\dfrac{\lambda_a(\boldsymbol{x}) - \lambda_1(\boldsymbol{x})}{\lambda_2(\boldsymbol{x}) - \lambda_1(\boldsymbol{x})}\right)^n (\lambda_a(\boldsymbol{x}) - \lambda_1(\boldsymbol{x}))^2}{(n+1)(n+2)}, & \lambda_1 \leq \lambda_a \geq \lambda_2 \\ E_f \left(\dfrac{(1 + \lambda_a(\boldsymbol{x}) - \lambda_1(\boldsymbol{x}))^{m+2} - 1}{(m+1)(m+2)} + \dfrac{\lambda_2(\boldsymbol{x}) - \lambda_a(\boldsymbol{x})}{m+1} - \dfrac{(\lambda_2(\boldsymbol{x}) - \lambda_1(\boldsymbol{x}))^2}{(n+1)(n+2)}\right), & \lambda_a \geq \lambda_2 \end{cases}$$

(S2.37)

where $\lambda_1$ and $\lambda_2$ are the lower and upper boundaries, respectively, that define a smooth transition region about $\lambda_c$, $E_f$ is the modulus of aligned fibers, and $m$ and $n$ are exponents that further characterize the function. The stress in the collagen matrix is also found as the sum of the stresses due to isotropic and fibrous components. Accordingly, the stress due to fiber alignment is found as:

$$\sigma_{ij}^{fiber}(\boldsymbol{x}) = \sum_{a=1}^{3} \frac{\lambda_a(\boldsymbol{x})}{J} \frac{\delta f}{\delta \lambda_a(\boldsymbol{x})} (n_i^a(\boldsymbol{x}) \otimes n_j^a(\boldsymbol{x}))$$

(S2.38)

where $J = \det(F_{ij}(\boldsymbol{x}))$ is the Jacobian of the deformation gradient, $F_{ij}$, and $n_i^a$ are the unit vectors in the direction of the principal stretches. In our simulations, the fibrous contribution to matrix mechanics given by Equation S2.41 was implemented as an applied stress calculated at all points in the matrix, and the passive strain energy of the collagen matrix was calculated by taking the volume integral of Equation S2.34 over the full volume of the matrix. $K_{ECM}$ was estimated at different values of collagen density, $d$, by performing a logarithmic fit of scaled values of the equilibrium modulus of collagen gels as previously reported (*16*) (Table S2). $\mu_{ECM}$ was then estimated as one-tenth the value of $K_{ECM}$. Values of parameters used in the model are listed in Table S2.

*2.3 Simulation procedure for the generalized 3D model:*

The 3D model derived here is implemented in COMSOL (*80*) within a finite element framework. We solve the steady state equations for stress and contractility obtained by minimizing the metabolic potential (Equations S2.20-S2.21) to find the predicted contractility, stress, and strain fields for a given cell shape. We then iteratively perform simulations for changing cell shape (changing cell body aspect ratio) and changing matrix stiffness, calculating the metabolic potential for each. The optimum shape at a given matrix stiffness is determined by the shape with the lowest metabolic potential. Cells were modelled using tetrahedral elements with a minimum element size of 0.225μm. The mechanical equilibrium equations, $\partial\sigma_{ij}/\partial x_i = 0$, are then solved at every integration point within the cell and matrix along with appropriate boundary conditions to determine the stress and strain fields (Figure S4C). Auxiliary variables for principal stresses and strains are utilized to handle the cyclical dependence introduced by $\sigma_a$ in Equation S2.21(a). To improve model convergence towards a solution in the case of some cell shapes, an estimate for $\sigma_a$ was used, $\sigma_a = \frac{1}{10}\left(AR_{xy}\right)^{0.3}\sigma_1(\boldsymbol{x})$, where $AR_{xy}$ is the in-plane cell aspect ratio. This form for $\sigma_a$ was utilized instead of the generalized $\sigma_a = \left(\frac{\sigma_1(x)}{\sigma_2(x)} - 1\right)\sigma_1(\boldsymbol{x})$ because in the specific case of the cell shapes modeled here, $\sigma_a$ scales with $AR_{xy}$ in a similar manner. The collagen matrix is modeled as a cylinder that ensconces the cell, and the 2D substrate is modeled as cylinder whose top surface is in contact with the bottom of the cell. Each ECM has a radius greater than 10 times that of the cell to reduce boundary effects.

The model parameters used to model a cell between microposts (Table S1) served as guidelines for choosing the biophysical parameters in these simulations. Iterative analysis was performed where model predictions of cell elongation and contractility at steady state was compared against corresponding experimental measurements in 2D and 3D micro-environments for MDA MB-231 cells, and the material parameters that give the best match were used as listed in Table S3. The mechanical properties of the nucleus are chosen such that it is stiffer than the cell.

3. Model for steady state ATP replenishment in cells

*3.1. Predicting steady state concentrations of adenosine nucleotides considering mechanosensitive activation of AMPK:*

In this section, we describe the model for ATP replenishment based on the mechanosensitive kinetics of AMPK activation (phosphorylation). We first describe the ATP:ADP ratio at different stages of cell spreading for cells on soft and stiff matrices:

- *Cell just seeded on a substrate*: It is assumed that cells have the same ATP:ADP ratio just after being seeded on the substrate (see Figure 4A), which is typically in the range of 1-10 (*43*).
- *Before cell reaches steady state*: As the cell spreads, ATP consumption is higher on the stiffer substrate, which leads to a lower ATP:ADP (and higher AMP:ATP) ratio for cells on stiff substrates that are not fully spread (Figure 4A).
- *At steady state:* The lower ATP:ADP ratio on stiff substrates also leads to the activation of more AMPK*, and consequently higher ATP:ADP ratio in cells at steady state (Figure 4B).

Experimental measurements conducted at different stages of cell spreading also show a similar trend and further validate the proposed mechanosensitive ATP replenishment pathway (*18*). While the dynamics of ATP consumption as the cell spreads is essential to reinforce the idea of a mechanosensitive ATP replenishment pathway activated before the cell reaches steady state, we will restrict attention in this study to predicting only the steady state ATP:ADP ratio as a function of matrix stiffness. The adenylate kinase reactions that govern ATP synthesis and consumption, the reaction rate constants and the biophysical parameters they depend on are listed in Table S4.

The rate constants that govern ATP hydrolysis are $k_1^+$ and $k_1^-$, while the conversion of AMP to ADP is governed by $k_2^+$ and $k_2^-$, and the synthesis of activated AMPK is governed by $k_3^+$ and $k_3^-$. Further, we assume that the total concentration of the nucleotides is constant and that the sum of $[AMPK]$ and $[AMPK^*]$ is constant, i.e.:

$$[ATP] + [ADP] + [AMP] = [ATOT] \tag{S3.1}$$

$$[AMPK] + [AMPK^*] = [AMPKTOT] \tag{S3.2}$$

The time evolution of concentration of ATP, AMP and AMPK* can be written as:

$$\frac{d[ATP]}{dt} = -k_1^+[ATP] + k_1^-[ADP][P_i] \tag{S3.3}$$

$$\frac{d[AMP]}{dt} = -k_2^+[AMP][P_i] + k_2^-[ADP] \tag{S3.4}$$

$$\frac{d[AMPK^*]}{dt} = k_3^+[AMP][AMPK] - k_3^-[AMPK^*] \tag{S3.5}$$

To determine the steady state levels of ATP, ADP and AMP, we set $\frac{d[ATP]}{dt} = \frac{d[AMP]}{dt} = \frac{d[AMPK^*]}{dt} = 0$. This condition along with equations (S3.1) and (S3.2), leads to a set of five equations that we solve simultaneously to determine $[ATP]$, $[ADP]$, $[AMP]$, $[AMPK]$ and $[AMPK^*]$ at steady state.

To solve, we first note the rate of ATP consumption in the cell due to active motors, $k_1^+$, is a function of the total volumetric stress found by integrating $\sigma_{kk}(\boldsymbol{x})$ over the cell volume, $k_1^+ = k_1^+(\int \sigma_{kk}(\boldsymbol{x})dV_c)$. To simplify notation, we denote total volumetric stress as $\sigma_{kk} = \int \sigma_{kk}(\boldsymbol{x})dV_c$ to avoid explicitly writing the volume integral every time it appears. We assign $k_1^+$ the functional form $k_1^+ = r_0\sigma_{kk}^l$ where $r_0$ is a constant and $l$ is an exponent that determines the sensitivity of the rate constant governing ATP consumption to stress. Additionally, the rate of ATP production, $k_1^-$, is a function of the amount of phosphorylated AMPK, $[AMPK^*]$. This allows us to write:

$$\frac{[ATP]}{[ADP][P_i]} = \frac{k_1^-}{k_1^+} = \frac{r_1[AMPK^*]}{r_0\sigma_{kk}^l} \tag{S3.6}$$

where we have assumed a linear dependence of $k_1^-$ on $[AMPK^*]$, the strength of which is determined by the constant $r_1$. Next, we can combine Equations S3.2 and S3.5 to yield:

$$[AMPK^*] = \frac{[AMP][AMPKTOT]k_3^+}{(k_3^- + k_3^+[AMP])}$$

(S3.7)

If we assume that the rate of AMPK dephosphorylation is significantly larger than the rate of AMPK phosphorylation ($k_3^- \gg k_3^+$), then we may simplify Equation S3.7 to:

$$[AMPK^*] = k_3[AMP][AMPKTOT]$$

(S3.8)

where $k_3 = \frac{k_3^+}{k_3^-}$. Combining Equations S3.8 and S3.6 yields:

$$\frac{[ATP]}{[ADP][P_i]} = \frac{r_1 k_3}{r_0 \sigma_{kk}^l}[AMP][AMPKTOT].$$

(S3.9)

Since the rate constant governing AMPK activation through mechanosensitive calcium signaling, $k_3$, is also an increasing function of cytoskeletal stress ($\sigma_{kk}$), we assign it the form $k_3(\sigma_{kk}) = r_2 {\sigma_{kk}}^m$. Like with $k_1^+$, this form is chosen to allow for a more generalized dependence of $k_3$ on $\sigma_{kk}$ as determined by the constants $r_2$ and $m.$ Next, we note that if we assume the ratio between the rate constants of the adenylate kinase reactions governing the interconversion of ADP and AMP is constant:

$$\frac{[AMP][P_i]}{[ADP]} = \frac{k_2^-}{k_2^+} = \gamma$$

(S3.10)

we may combine Equation S3.9 with Equations S3.4 and S3.10 to obtain:

$$[ATP] = \frac{r_1 r_2 \gamma [AMPKTOT][ADP]^2}{r_0}\sigma_{kk}^{m-l}$$

(S3.11)

Lastly, we make use of Equation S3.1 to yield:

$$[ADP] = \frac{[ATOT] - [ATP]}{(1+\gamma')}$$

(S3.12)

where $\gamma' = \frac{\gamma}{[P_i]}$, and we assume $[P_i]$ to be a constant here. Substituting Equation S3.12 into S3.11 leads to a quadratic equation that can be solved for $[ATP]$ and subsequently $[ADP]$ and $[AMP]$:

$$[ATP] = [ATOT]\left(\frac{c}{\sigma_{kk}^n} + 1 - \sqrt{\left(\frac{c}{\sigma_{kk}^n} + 1\right)^2 - 1}\right)$$

(S3.13a)

$$[ADP] = \frac{[ATOT]}{(1+\gamma')}\left(\sqrt{(\frac{c}{\sigma_{kk}^{n}}+1)^2 - 1} - \frac{c}{\sigma_{kk}^{n}}\right)$$

(S3.13b)

$$[AMP] = \frac{\gamma'}{(1+\gamma')}[ATOT]\left(\sqrt{(\frac{c}{\sigma_{kk}^{n}}+1)^2 - 1} - \frac{c}{\sigma_{kk}^{n}}\right)$$

(S3.13c)

from which $\frac{[ATP]}{[ADP]}$ can be found. The final expression for the steady state ATP:ADP ratio is:

$$\frac{[ATP]}{[ADP]} = \frac{(1+\gamma') * \left(\frac{c}{\sigma_{kk}^{n}} + 1 - \sqrt{\left(\frac{c}{\sigma_{kk}^{n}}+1\right)^2 - 1}\right)}{\sqrt{(\frac{c}{\sigma_{kk}^{n}}+1)^2 - 1} - \frac{c}{\sigma_{kk}^{n}}}$$

(S3.14)

where $c = \frac{r_0(1+\gamma')^2}{r_1 r_2 \gamma [AMPKTOT][ATOT]}$ and $n = m - l$. Since the reported value for the steady state ratio of $[AMP]/[ADP]$ is ~0.3, we assign $\gamma'$ (the constant representing this ratio (see Equation S3.10)) the value of 0.3 (*81*). Using model predictions of $\sigma_{kk}$ in optimally shaped cells at various 3D collagen densities, we perform a fit of this function to experimentally measured ATP:ADP ratios using PercevalHR (*16*) and determine the constants $c$ and $n$ using the linear least squares method. Results are reported in Table S5 and Figure S15. All 3D model calculations for cells in 3D collagen, on 2D substrates, and on confining micropatterns are performed using these values. Note that since experimental data is reported as normalized values of PercevalHR signal, the fit was performed using these normalized values as opposed to true concentrations.

Interestingly, Equation S3.14 predicts that the steady state ATP:ADP ratio will increase with increasing stress (or increasing ECM stiffness) as long as the parameters $c$ and $n$ are both positive. Since $c$ is comprised of other factors that are all positive quantities by definition, $c$ will always be positive regardless of how its factors such as $r_1$, $r_2$, or $\gamma$ may change in value. However, $n$ describes the relative strength of the change in the rate parameters $k_1^+$ and $k_3$ in response to a change in stress, which means it will only be positive if $k_3$ (the rate of calcium-dependent AMPK activation) shows a larger increase in response to stress than $k_1^+$ (the rate of ATP consumption by motors). Therefore, an increasing trend in ATP:ADP ratio with increasing ECM stiffness is obtained only when cells experience a larger increase in the rate of AMPK activation than in the rate of ATP consumption by motors. Likewise, Equation S3.14 predicts that ATP:ADP ratio will show a decreasing trend with ECM stiffness if the opposite is true and the rate of ATP consumption increases faster than the rate of AMPK activation with increasing stiffness.

#### 3.2. <u>Comparing model predictions with experimental measurements of cellular energetics</u>

##### *3.2.1. <u>Model prediction of [AMPK*]:</u>*

Our model allows for the quantitative prediction of the concentration of activated (phosphorylated) AMPK within a cell dependent on mechanosensitive signaling with few assumptions. At steady state, Equation S3.5 leads to the following prediction for the concentration of activated AMPK, [AMPK*]:

$$[AMPK^*] = \frac{k_3^+}{k_3^-}[AMPK][AMP] \tag{S3.15}$$

Combining with Equation S3.5, we find

$$[AMPK^*] = [AMPKTOT]\left(\frac{k_3[AMP]}{1 + k_3[AMP]}\right). \tag{S3.16}$$

If we assume $k_3[AMP] \ll 1$, then Equation S3.8 simplifies to:

$$[AMPK^*] = k_3[AMP][AMPKTOT]. \tag{S3.17}$$

Equation S3.17 can be used to find the normalized value of $[AMPK^*]$ at different 3D collagen matrices or different 2D polyacrylamide substrates by plugging in the predicted values for $[AMP]$ (Equation S3.13c) and the functional form for $k_3$ ($k_3 = \sigma_{kk}^m$). The normalized [AMPK*] value can be determined as:

$$\frac{[AMPK^*]}{[AMPKTOT][ATOT]} = \frac{\gamma'}{(1+\gamma')}\left(\sqrt{(\frac{c}{\sigma_{kk}^n} + 1)^2 - 1} - \frac{c}{\sigma_{kk}^n}\right)\sigma_{kk}^m \tag{S3.18}$$

One must assume a value for $m$ when calculating $[AMPK^*]$, and we have assumed its value as $m = 2n$ when calculating $[AMPK^*]$ for cells in 3D collagen and on confining micropatterns, and as $m = 1.5n$ when calculating $[AMPK^*]$ for cells on 2D substrates. Note that unlike the ATP:ADP ratio (Equation S3.14), the predicted AMPK activation increases with $\sigma_{kk}$ regardless of the values chosen for parameter $n$. The values of $[AMPK^*]$ predicted by Equation S3.18 are used to compare with experimentally measured amounts of phosphorylated AMPK (Figure 4F) as well as with other measures of cell energetics such as glucose uptake (Figures 4G,H) and mitochondrial membrane potential (Figure 5B, Figure S16, and Figure S17) by normalizing by the value of $[AMPK^*]$ at the stiffest substrate condition (2D) or highest collagen density (3D), or the most confining micropattern. To obtain predicted values, we assume $[AMPKTOT]$ and $[ATOT]$ are constant. We note that the predicted values of the change in mitochondria membrane potential in cells on micropatterns with fixed elongation and changing spread area are higher than the experimentally measured values (Figure S17). This discrepancy suggests the involvement of additional biochemical pathways that may limit mitochondrial activity when cells adopt a spread but not elongated morphology.

*3.2.2.* *Model prediction of ATP consumption rate:* As outlined in SI Section 2.2.3, the rate at which the cell consumes energy to maintain its contractility at steady state is equivalent to the rate at which the cell dissipates heat because there are no associated changes in conservative energy. We therefore compute the cell's total ATP consumption rate with Equation S2.27 by integrating the spatially varying heat dissipation rate over the cell volume in optimally shaped cells determined by minimizing the metabolic potential.

*3.2.3.* *Simulating myosin motor inhibition:* To simulate lower activity of myosin motors, we use a smaller value for $\alpha$, the parameter that reflects the average lifetime of stress fibers (Equation S2.19). This is equivalent to lowering the motor binding rate, $k_{on,ij}$, and/or increasing the motor unbinding rate, $k_{off,ij}$, to reflect a shift in the dynamic equilibrium of motors towards the unbound, inactive state that arises with experimental inhibition of myosin. To simulate control cells, we use the same value of $\alpha$ as in the rest of our simulations, $\alpha = 2.33\ kPa^{-1}$ (Table S3), while cells having been treated with myosin inhibitor are simulated with the much smaller value of $\alpha = 0.01\ kPa^{-1}$. Consequently, the predicted steady state contractility and stress developed in the cell is lower (Figure S12), as is the predicted ATP:ADP ratio (Figure 4J).

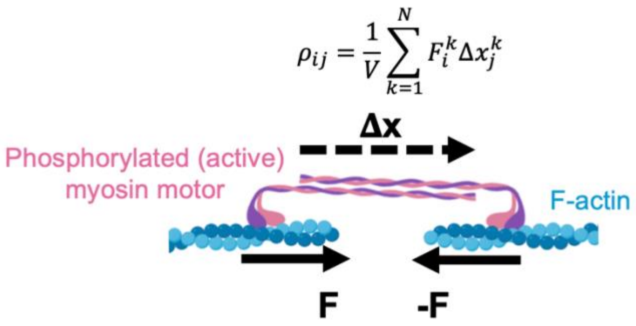


**Figure S1: Active myosin motors are modelled as force dipoles.** Contractility generated by myosin motors is represented as a tensor, $\rho_{ij}$, in our model whose components represent the motor density in respective directions.

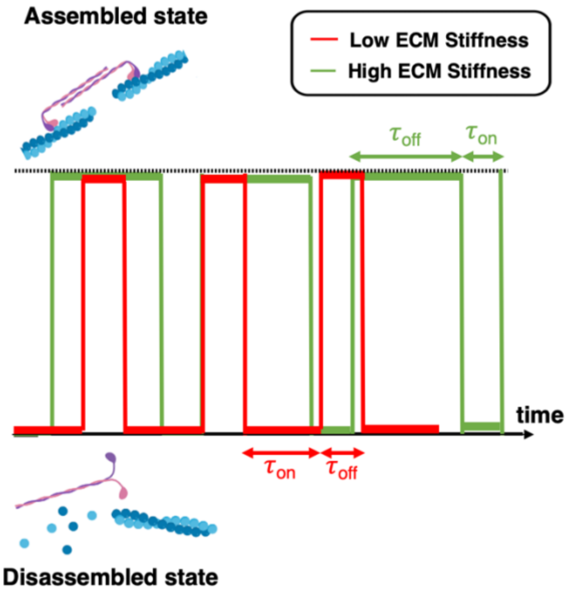


**Figure S2: ECM stiffness regulates stress fiber assembly/disassembly kinetics.** Actin stress fibers in cells undergo a continuous cycle of assembly and disassembly. This cycle can be represented by considering the lifetime of stress fibers, $\tau_{off}$, and the time required for stress fiber formation, $\tau_{on}$. These times also relate to the rates of stress fiber formation such that $k_{on}(\sigma_{ij}) = 1/\tau_{on}(\sigma_{ij})$ and $k_{off}(\sigma_{ij}) = 1/\tau_{off}(\sigma_{ij})$. The dependence on stress, $\sigma_{ij}$, reflects the mechanosensitive nature of the pathways governing assembly (Figure 1A). In low stiffness environments, the ratio of time spent in the assembled state, $\tau_{on}$, relative to the time spent in the disassembled state, $\tau_{off}$, is higher than for high stiffness environments in which mechanosensitive feedback upregulates stress fiber formation.

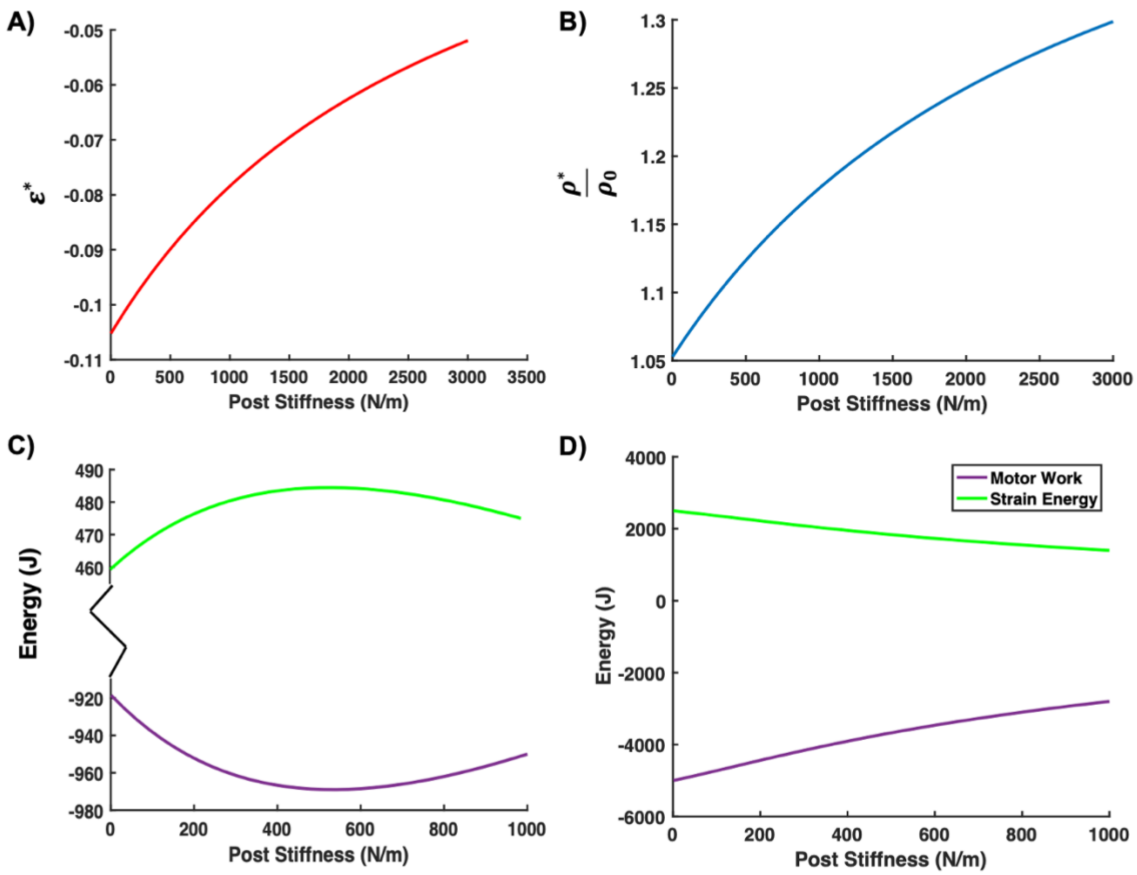


**Figure S3: Steady state strain and contractility for a cell between microposts.** Variation of the steady state (**A**) cell strain and (**B**) contractility as a function of micropost stiffness. (**C**) Non-monotonic variation in strain energy and motor work with post stiffness showcased for parameter values $\alpha = 7e^{-3}\ Pa^{-1},\ \beta = 10e^{-3}\ Pa^{-1}, K = 450\ Pa$. (**D**) Monotonic variation in strain energy and motor work with post stiffness for parameter values $\alpha = 7e^{-3}\ Pa^{-1},\ \beta = 10e^{-3}\ Pa^{-1}, K = 200\ Pa$.

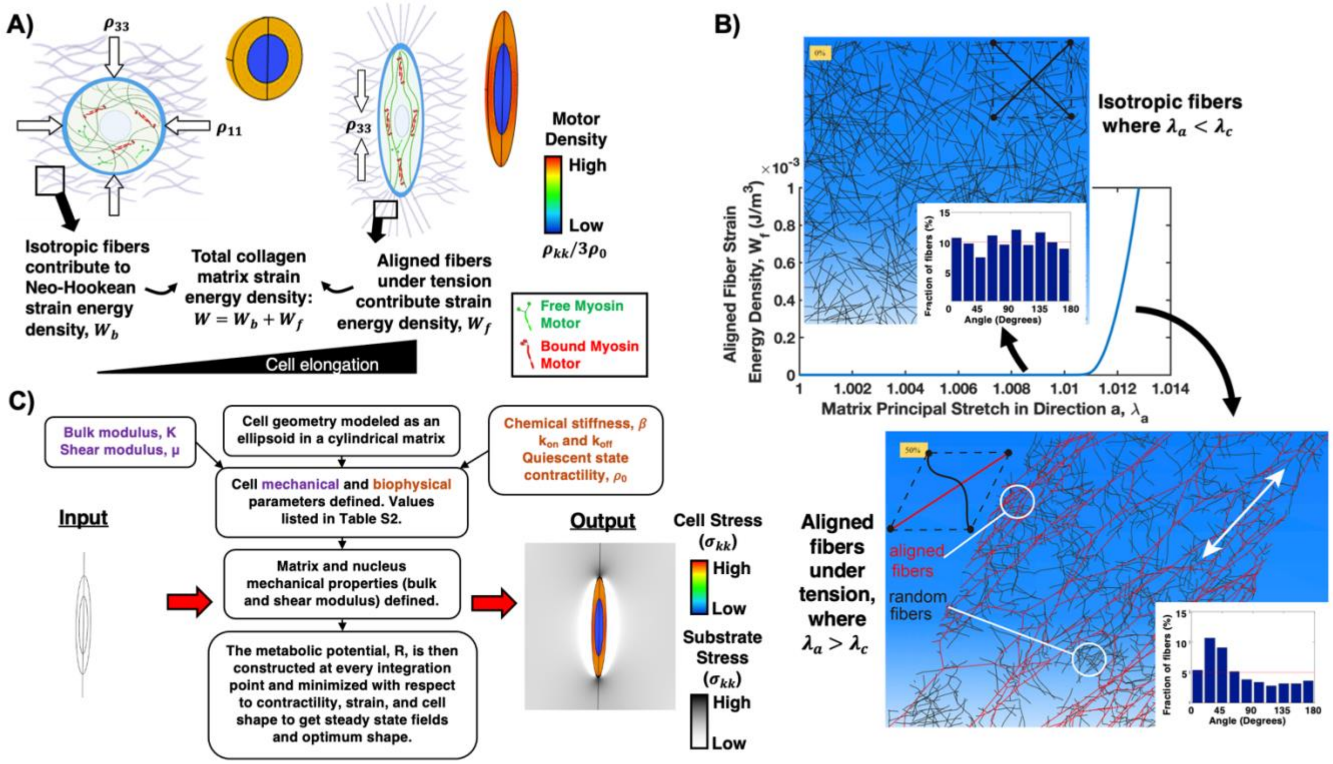


**Figure S4: Implementation of 3D collagen model in a finite element framework.** (**A**) Depiction of the tensorial nature of contractility and a schematic illustrating the non-linear fiber model used to represent collagen which captures the preferential alignment of fibers along the direction of maximum principal stretch $\lambda_a$, (**B**) Graphical illustration of the strain energy density function $f(\lambda_a)$ along with representation of fiber alignment beyond a critical stretch $\lambda_c$ as reported from discrete fiber simulations (*47*), and (**C**) Flowchart depicting the method of solution employed along with workflow for determining the steady state strain and contractility fields.

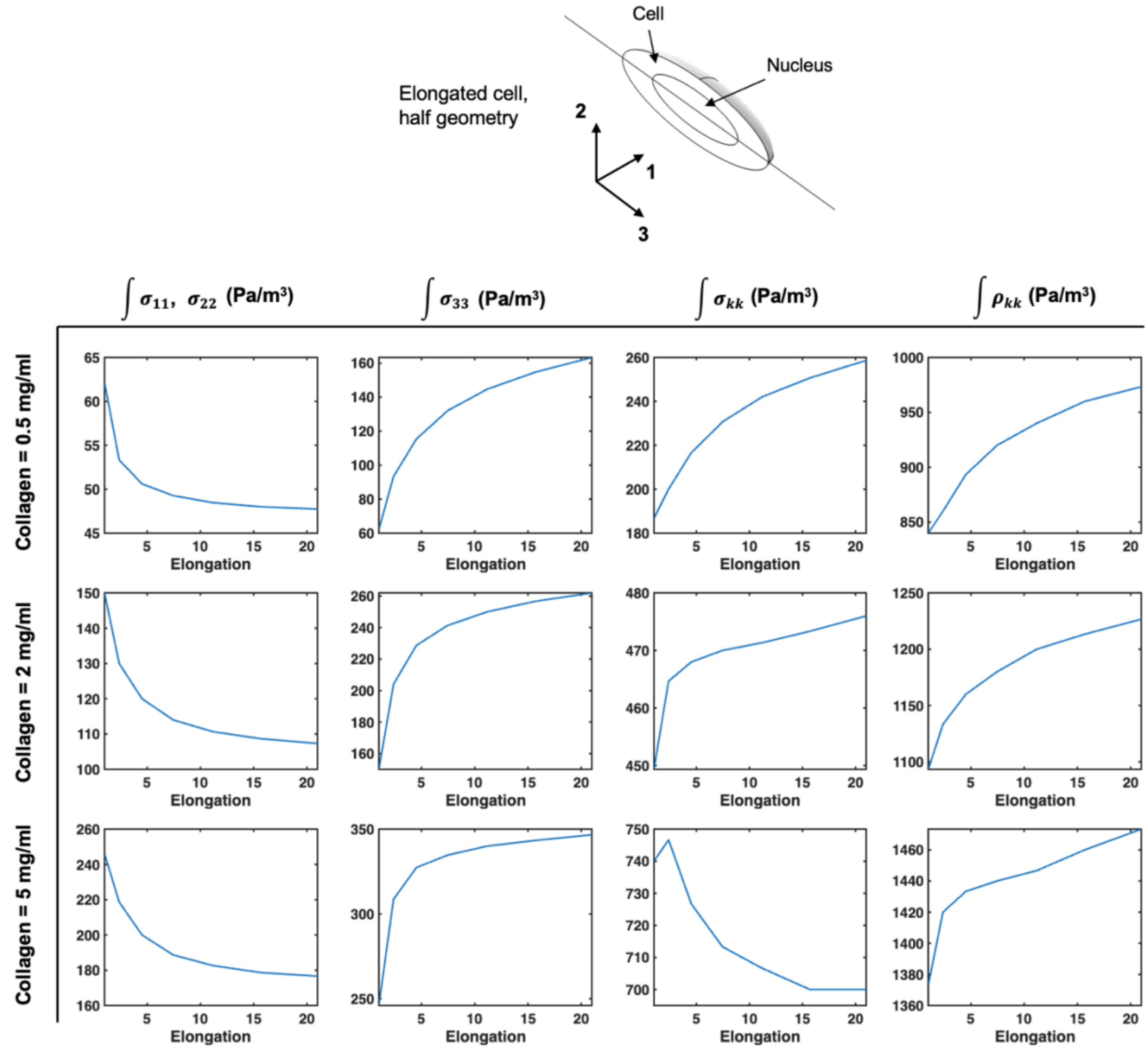


**Figure S5: Stress components as a function of cell elongation.** Variation of the stress components with cell elongation for low (0.5 mg/ml), intermediate (2 mg/ml), and high (5 mg/ml) collagen densities.

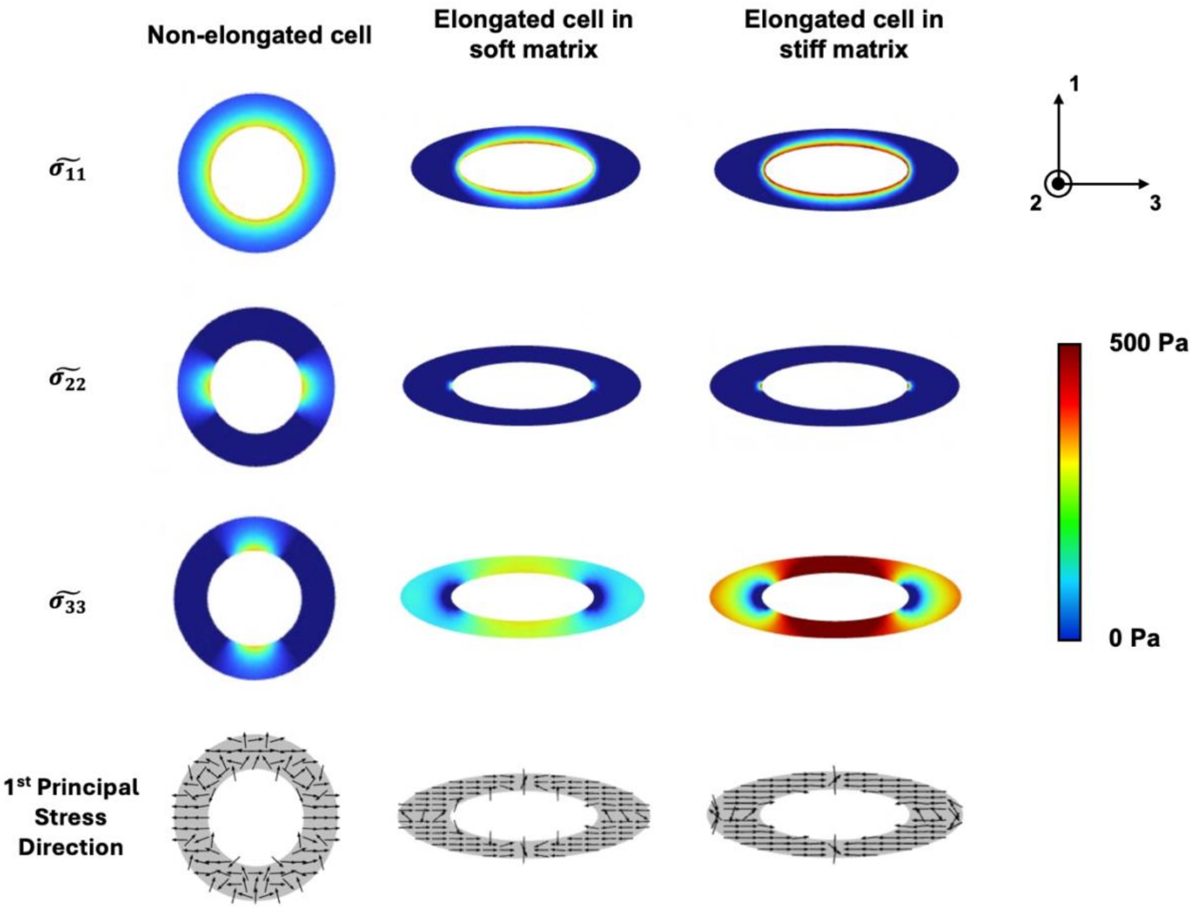


**Figure S6: Cross-section contour plots of deviatoric stress components of cells in 3D collagen.** Apart from small differences in stress concentration around the nucleus (central region of the cell, excluded on these plots), only the deviatoric stress along the direction of elongation (the 3-direction, $\widetilde{\sigma_{33}}$) shows significant increases throughout the cell body as the cell elongates, indicating preferential polarized alignment of stress fibers in this direction. Plotting the direction of first principal stress (shown by black arrows) throughout the cytosol (shown in gray) provides further confirmation that stress primarily develops along the direction of elongation.

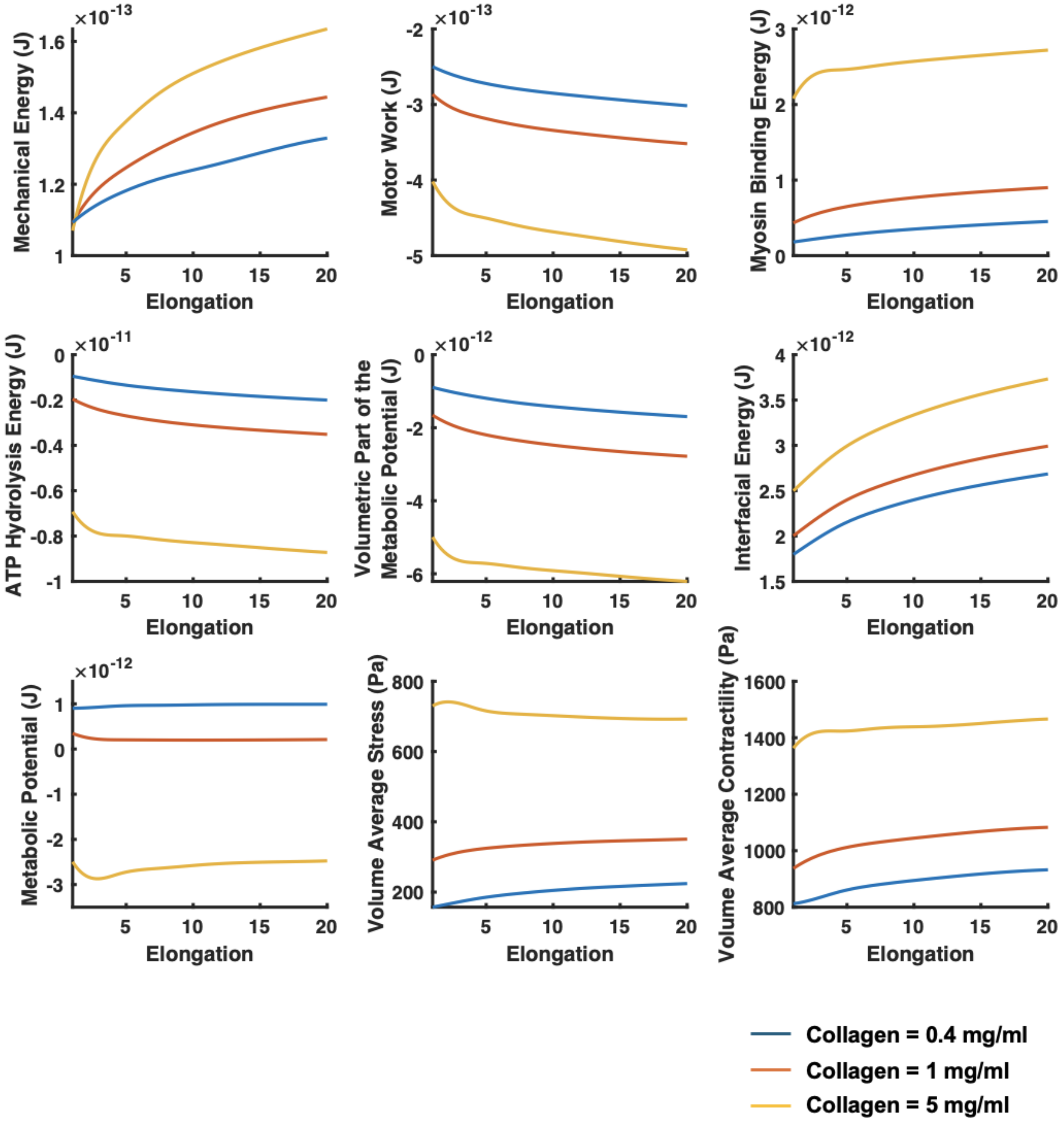


**Figure S7: Trends in energy, stress, and contractility in 3D collagen.** Breakdown of the different energy contributions as a function of cell elongation and matrix density for cells in 3D collagen.

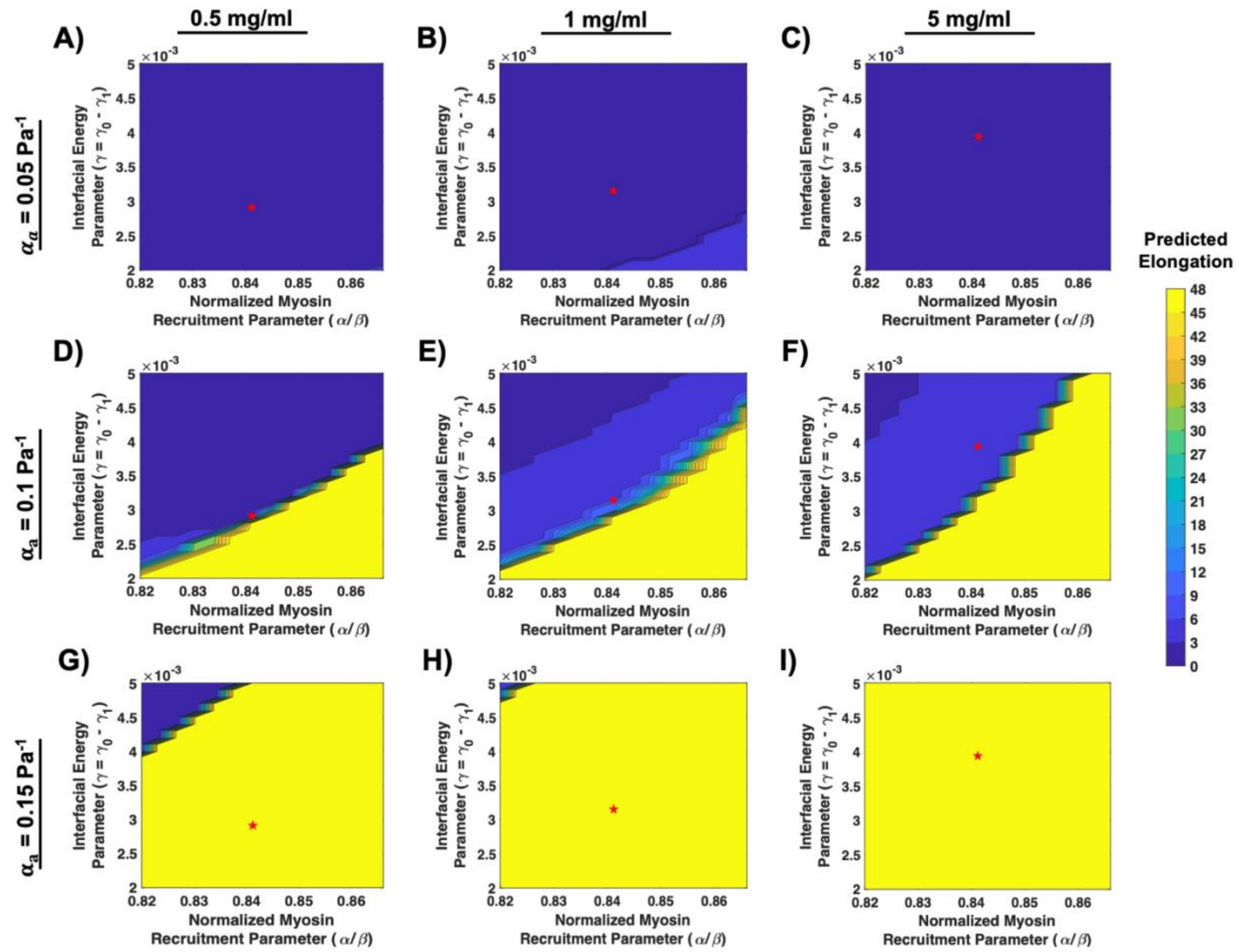


**Figure S8: Sensitivity analysis for interfacial energy parameter.** Sensitivity analysis done to determine the functional dependence of interfacial energy on other model parameters for collagen densities of 0.5 (**A,D,G**), 1 (**B,E,H**) and 5 (**C,F,I**) mg/ml and $\alpha_a$ values of 0.05 (**A,B,C**), 0.1 (**D,E,F**), and 0.15 (**G,H,I**) $Pa^{-1}$. Red stars indicate the parameter value combinations that give a good match with experimentally observed cell elongations at $\alpha_a$ = 0.1 and $\alpha/\beta$=0.841. These values were used to construct the functional dependence of the interfacial energy parameter on collagen density and were used in all subsequent simulations. Note that elongations >48 were not simulated as cell shapes begin to become extremely thin and unphysical.

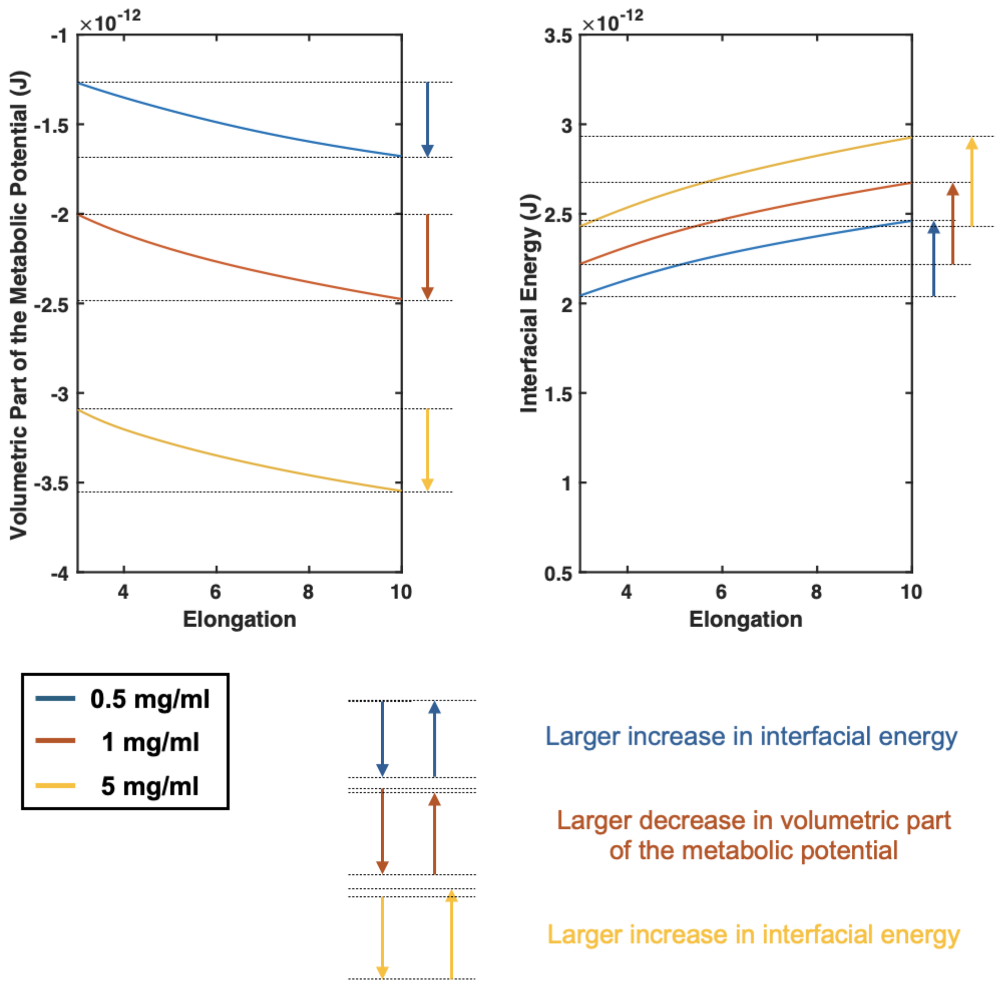


**Figure S9**: **Visualization of the competing magnitudes of the volumetric part of metabolic potential and interfacial energy in elongating cells.** A detailed view and comparison of the variation in the magnitudes of the volumetric part of the metabolic potential and interfacial energy at low (0.5 mg/ml), intermediate (1 mg/ml) and high (5 mg/ml) matrix densities. Only at intermediate densities is the decrease in the volumetric part of the metabolic potential with cell aspect ratio larger than the increase in interfacial energy, which promotes cell elongation.

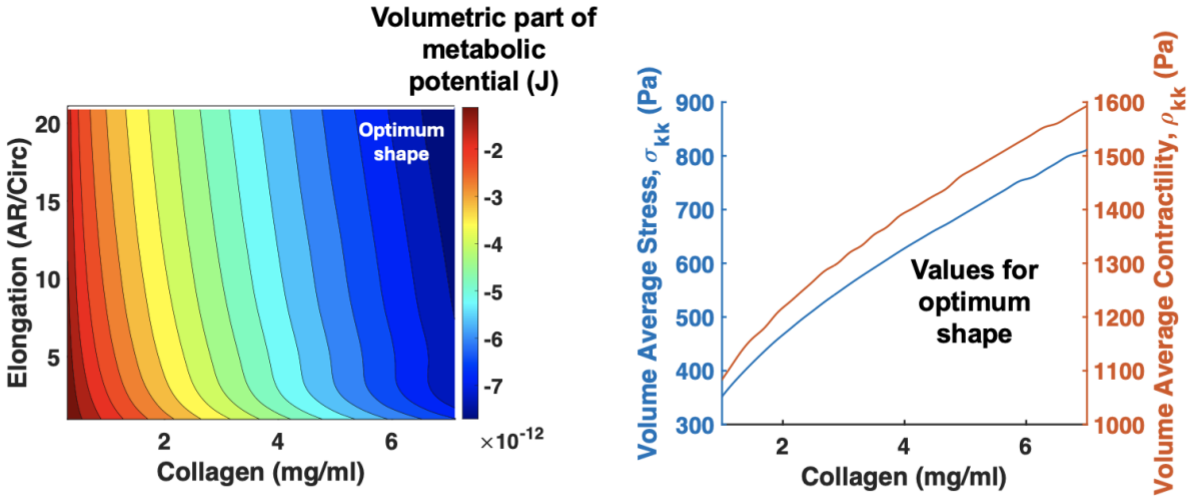


**Figure S10: Volumetric part of the metabolic potential predicts extremely elongated optimum cell shapes without interfacial energy.** Cell elongation in collagen matrices predicted by the volumetric part of the metabolic potential without considering interfacial energy predicts highly elongated optimum shapes at all collagen densities and monotonic increases in stress and contractility.

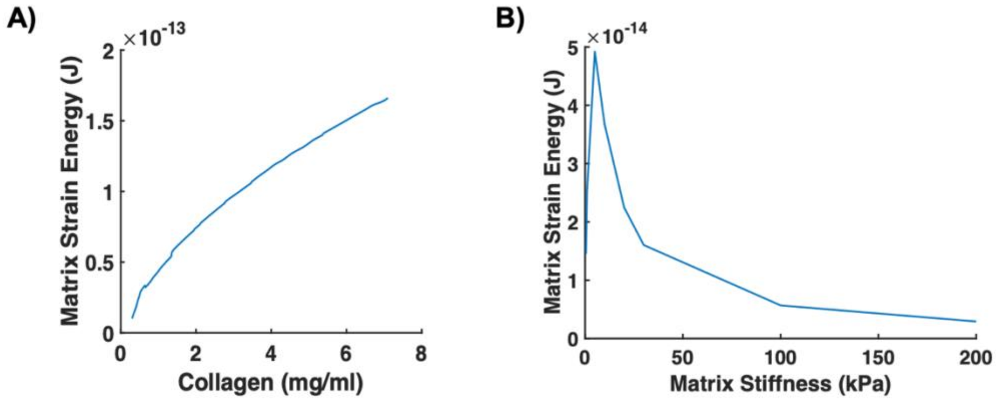


**Figure S11: Strain energy of the ECM around cells in 2D and 3D:** Model prediction of total ECM strain energy for optimum shaped cells in (**A**) 3D collagen and (**B**) on 2D PA substrates.

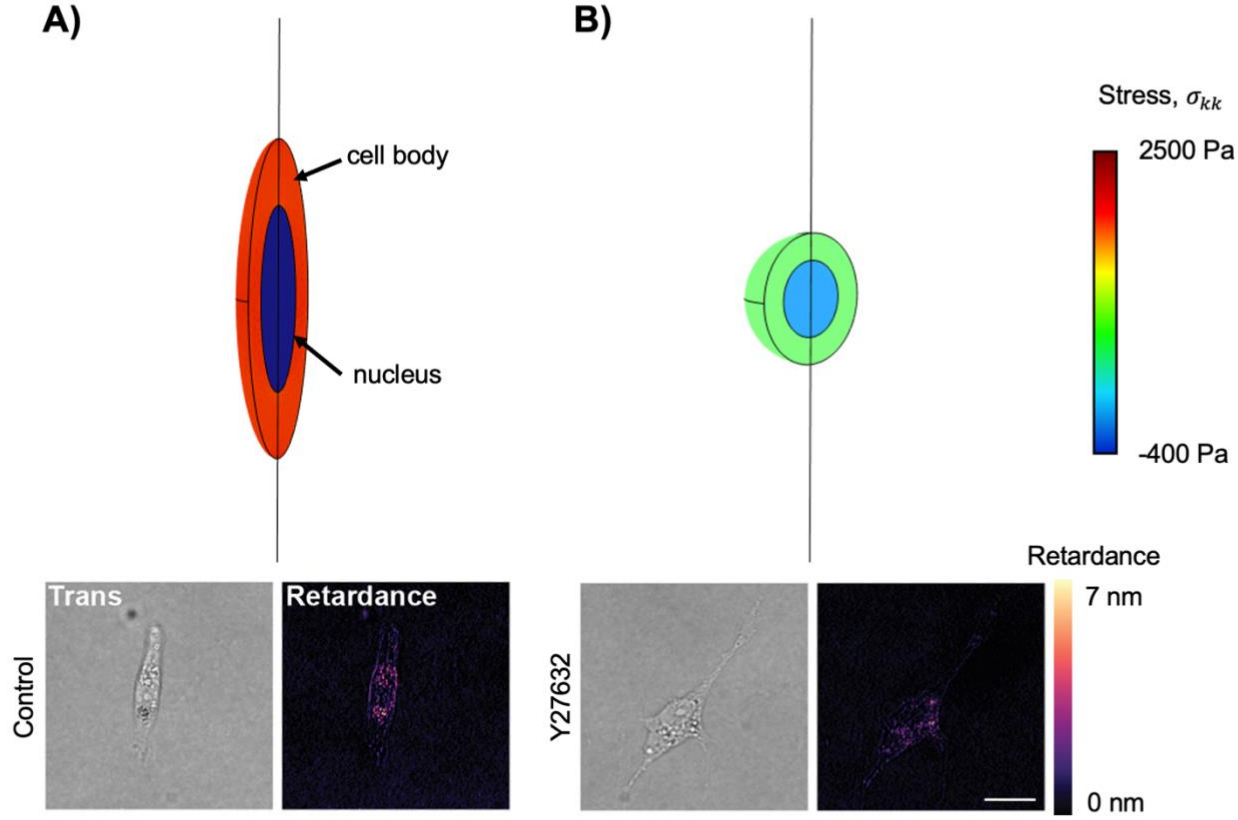


**Figure S12: Stress distribution in optimally shaped cells after inhibition of myosin recruitment**: Model prediction of stress throughout the cell body in 1.5 mg/ml collagen for (**A**) a typical value of stress fiber assembly rate ($\alpha = 2.33\ kPa^{-1}$) and (**B**) a lower value ($\alpha = 0.01\ kPa^{-1}$) simulating myosin inhibition along with experimental measurements of optical retardance revealing cell contractility from (*45*). Scale bar = 25 µm.

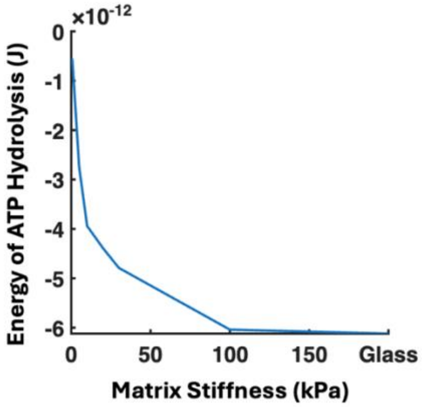


Energy available from ATP Hydrolysis = $-\int_0^{V_c} F_{ATP} dV_c$

**Figure S13: Energy available from ATP hydrolysis for cells on 2D substrates.** Energy of ATP hydrolysis available in optimum shaped cells at steady state as a function of matrix stiffness.

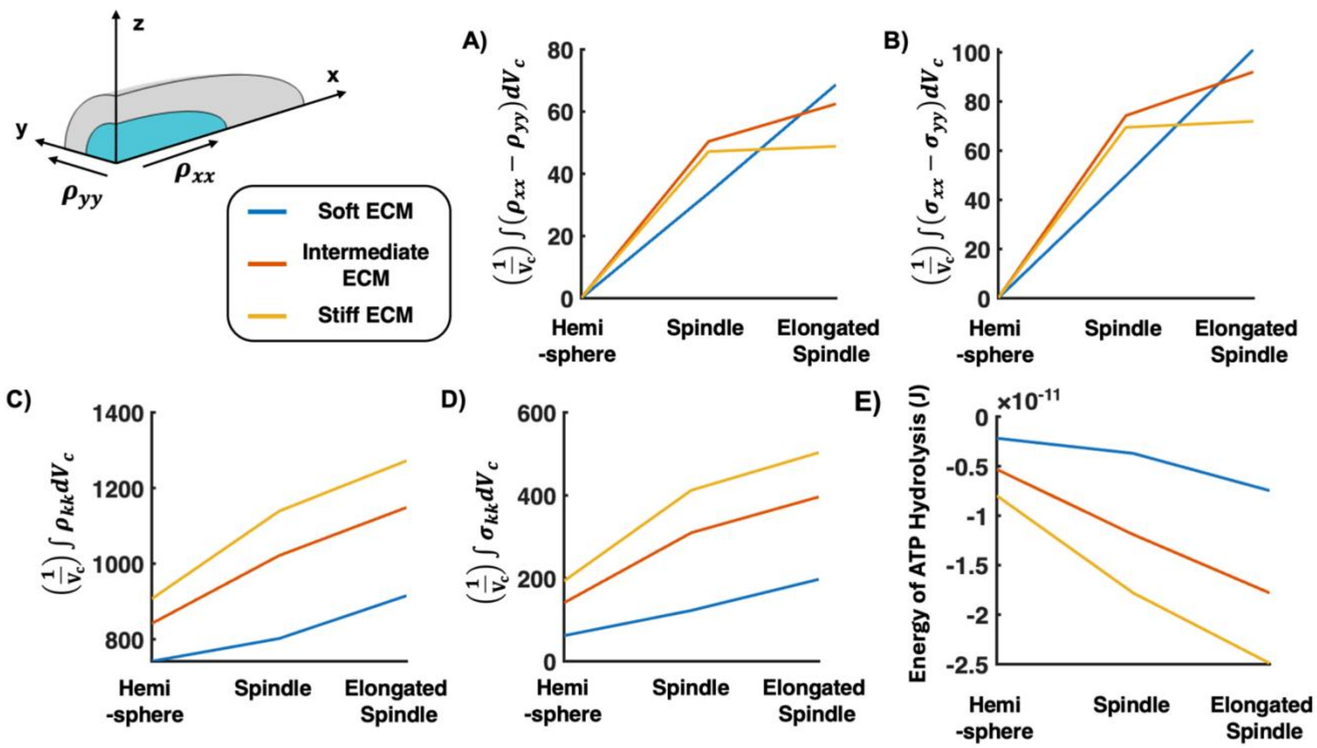


**Figure S14: Cell polarization, contractility and cytoskeletal stress as a function of cell shape and matrix stiffness on 2D substrates.** (**A**) Polarization in cells as a function of cell shape for different matrix stiffness levels, and (**B**) Polarization of stress components as function of cell configuration for soft, intermediately stiff, and stiff ECM. Variation of (**C**) normalized contractility (represents number density of myosin motors engaged on stress fibers), (**D**) normalized cytoskeletal stress, and (**E**) energy of ATP hydrolysis with cell shape and ECM stiffness.

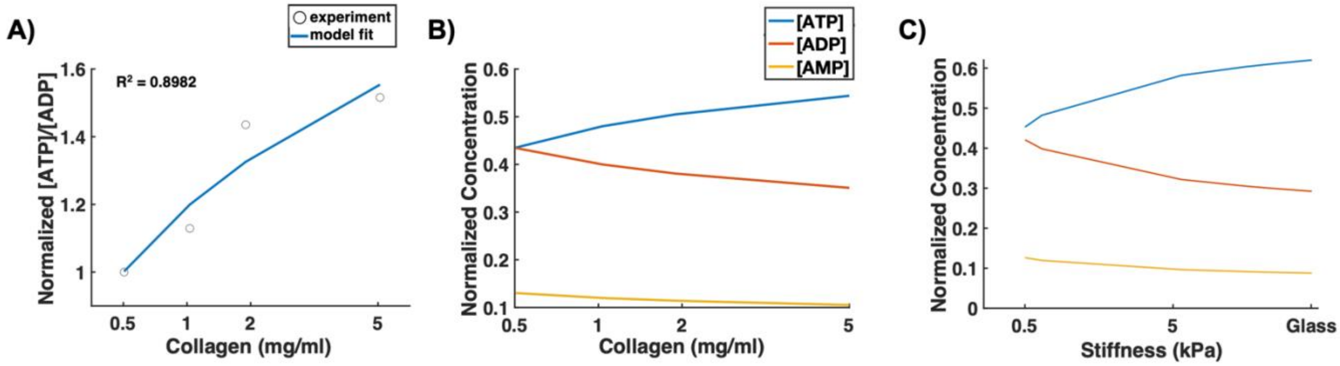


**Figure S15: Least squares fitting to determine model parameters** (**A**) Result of nonlinear least squares fit of model for [ATP]:[ADP] against experimentally obtained mean values using the PercevalHR sensor from (*16*). Predicted [ATP], [ADP], and [AMP] (normalized to total adenosine nucleotide concentration, [ATOT]) as a function of (**B**) 3D collagen density and (**C**) 2D matrix stiffness.

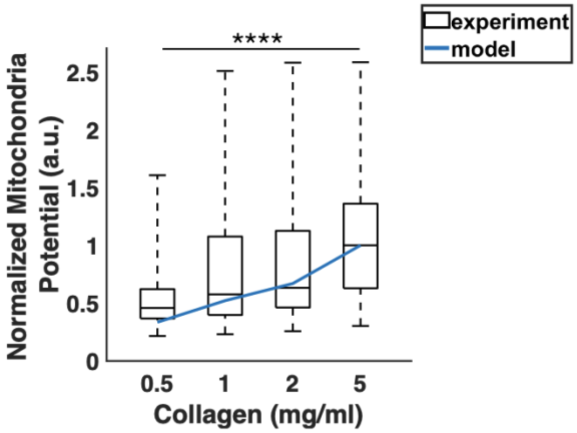


**Figure S16: Normalized mitochondria membrane potential for cells in 3D collagen.** Comparison of the normalized mitochondria membrane potential predicted by theory with experimental values obtained from TMRM measurements in MDA-MB-231 cells in 3D collagen (*16*). Box plots show median, 25$^{th}$/75$^{th}$ percentiles, and 5$^{th}$/95$^{th}$ percentiles *p<0.05, **p<0.01, ***p<0.001, ****p<0.001 with Kruskal-Wallis one-way ANOVA.

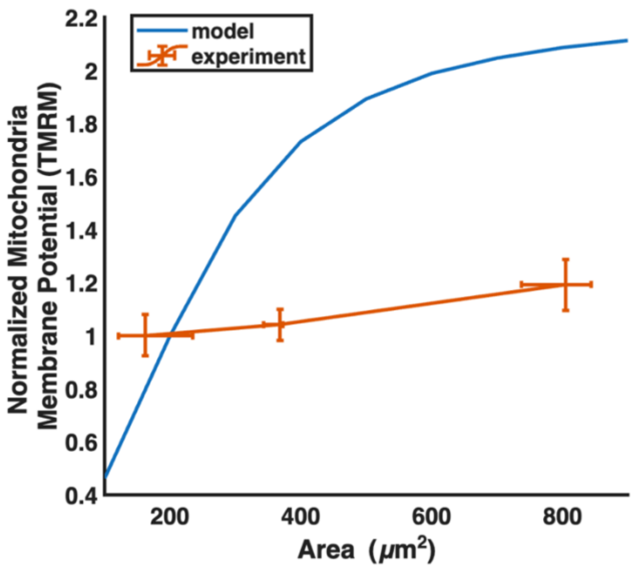


**Figure S17: Normalized mitochondria membrane potential for cells on confining micropatterns.** Comparison between the normalized mitochondria membrane potential (TMRM) measured in cells on micropatterns with fixed elongation and model predictions. Plot shows mean ±SD. Scale bar = 50 μm. Experimental data obtained from (*16*).

| Parameter | Value | Description |
|---|---|---|
| $\beta$ | $20\ kPa^{-1}$ | Chemical Stiffness |
| $\rho_0$ | $100\ kPa$ | Quiescent state contractility |
| $\tilde{\tau} = \frac{k_{on}(\sigma)}{k_{on}(\sigma) + k_{off}(\sigma)} = \alpha\sigma\left(\frac{p_0}{\Delta G}\right)$ | $\alpha = 7\ kPa^{-1}$ | Average stress fiber lifetime |
| $K$ | $100\ Pa$ | Cytoskeleton bulk modulus |
| $K_p$ | $2000\ Pa$ | Micropost bulk modulus |

**Table S1.** Mechanical and biophysical parameters used in simulating a cell between two microposts.

| Parameter | Value | Description |
|---|---|---|
| $E_f$ | 400 Pa | Modulus of aligned fibers |
| $\lambda_c$ | 1.001 | Critical stretch parameter beyond which fiber alignment is observed |
| $\lambda_t$ | $0.25\lambda_c$ | Transition region parameter |
| $\lambda_1$ | $\lambda_c - \frac{\lambda_t}{2}$ | Transition region lower limit |
| $\lambda_2$ | $\lambda_c + \frac{\lambda_t}{2}$ | Transition region upper limit |
| $m$ | 10 | Strain stiffening exponent |
| $n$ | 15 | Transition region exponent |
| $K_{ECM}$ | $693.58ln(d) + 920.65$ Pa | Initial collagen bulk modulus |
| $\mu_{ECM}$ | $\frac{K_{ECM}}{10}$ Pa | Initial collagen shear modulus |

**Table S2**: List of material parameters used to model behavior of fibrous collagen.

| Parameter | Value (3D) | Value (2D) | Description |
|---|---|---|---|
| $V_c$ | $1500\ \mu m^3$ | $1500\ \mu m^3$ | Cell Volume |
| $\beta$ | $2.77\ kPa^{-1}$ | $2.77\ kPa^{-1}$ | Chemical Stiffness |
| $K$ | $1.67\ kPa$ | $1.67\ kPa$ | Bulk Modulus of Cell |
| $\mu$ | $0.357\ kPa$ | $0.357\ kPa$ | Shear Modulus of Cell |
| $\rho_0$ | $1\ kPa$ | $1\ kPa$ | Quiescent State Contractility |
| $K_{nuc}$ | $2.5\ kPa$ | $2.5\ kPa$ | Bulk Modulus of Nucleus |
| $\mu_{nuc}$ | $0.536\ kPa$ | $0.536\ kPa$ | Shear Modulus of Nucleus |
| α | $2.33\ kPa^{-1}$ | $2.33\ kPa^{-1}$ | Average lifetime of stress fibers determined by stress-dependent signaling |
| $\alpha_a$ | $10\ kPa^{-1}$ | $10\ kPa^{-1}$ | Average lifetime of stress fibers determined by stress anisotropy-dependent signaling |
| $\gamma(d)$ | $\gamma(d) = \gamma(1+f(d))S$<br>$f(d) = g_0 d^{g_1}$<br>$\gamma = 0.00158$ J/m$^2$<br>$g_0 = 1$<br>$g_1 = 0.25$ | | Interfacial energy function (3D) |
| $\gamma^{CA}$ | | 0.018 J/m$^2$ | Interfacial energy parameter (cell-air) (2D) |
| $\gamma^{CM}(E)$ | | $\gamma^{CM}(E) = \gamma^{CM}(1-f^{CM}(E))$<br>$f^{CM}(E) = g_0 E^{g_1}$<br>$\gamma^{CM} = 0.001$ J/m$^2$<br>$g_0 = 0.25$<br>$g_1 = 0.2$ | Interfacial energy along cell-matrix interface (2D) |

**Table S3**: List of mechanical and biophysical model parameters used in simulating the response of MDA MB-231 cells in 2D and 3D micro-environments.

| Chemical Reaction | Biophysical parameters that govern reaction rate |
|---|---|
| (1) $ATP + H_2O \xrightarrow{k_1^+} ADP + P_i + \Delta G^*$ | $k_1^+ = f\left(k_{on}(\sigma_{kk})\right) = r_0 \sigma_{kk}^J$ |
| (2) $ADP + P_i \xrightarrow{k_1^-} ATP$ | $k_1^- = f_1([AMPK^*]) = r_1[AMPK^*]$ |
| (3) $AMP + P_i \underset{k_2^-}{\overset{k_2^+}{\Leftrightarrow}} ADP$ | $\gamma' = \frac{k_2^-}{k_2^+}$ |
| (4) $AMP + AMPK \underset{k_3^-}{\overset{k_3^+}{\Leftrightarrow}} AMPK^*$ | $k_3^+ = f_2([AMP], [Ca^{2+}])$ |

**Table S4:** Important biophysical parameters in the ATP replenishment model. Here, $\Delta G^*$ is the energy released due to hydrolysis of 1 mol of ATP.

| Parameter | Value | Description |
|---|---|---|
| $n$ | 0.504 | Measure of sensitivity of calcium signaling and Rho-ROCK signaling to stress |
| $c$ | 1.053E-7 | Factor incorporating other model constants including $r_0$, $r_1$, $r_2$, $[ATOT]$, $[AMPKTOT]$, and $\gamma$ |

**Table S5.** List of parameters required for prediction of ATP:ADP ratio determined by least-squares fitting.

| Parameter | Value | Description |
|---|---|---|
| $\alpha$ | 2.33e-3 1/Pa | Parameter representing strength of stress-dependent motor recruitment or average lifetime of stress fibers |
| $\alpha_a$ | 0.01 1/Pa | Parameter representing strength of stress anisotropy-dependent motor recruitment or average lifetime of stress fibers |
| $\Delta$G | -5.06e-20 J | Represents the energy of a single ATP bond |
| $V_c$ | 1.5E-15 $m^3$ | Volume of the cell |
| $p_0$ | 5e-20 J | Dipole strength of a single myosin motor |
| $c$ | 0.0008$s^{-1}$ | Estimated rate for stress fiber assembly |

**Table S6:** Parameters used to calculate the steady state cell power.
